# Spatiotemporal dynamics of tumor microenvironment remodeling

**DOI:** 10.1101/2025.07.15.662972

**Authors:** Kamil Lisek, Tancredi Massimo Pentimalli, Ilan Theurillat, Svea Beier, Daniel León-Periñán, Anastasiya Boltengagen, Anna Antonatou, Tasneem Cheytan, Serafima Dubnov, Marion Müller, Florian Hubl, Artemis Xhuri, Hanna Romanowicz, Beata Smolarz, Elodie Montaudon, Sandra Raimundo, Anca Margineanu, Marie Schott, Séverine Kunz, Elisabetta Marangoni, Nikos Karaiskos, Dana Ishay Ronen, Stefan Florian, Mor Nitzan, Walter Birchmeier, Nikolaus Rajewsky

## Abstract

During tumorigenesis, interactions between tumor and stromal cells progressively remodel the tumor microenvironment (TME) towards pro-tumoral functions. Understanding early TME remodeling dynamics is therefore crucial for developing interceptive therapies. However, clinical samples typically provide isolated, late tumorigenesis snapshots. To overcome this limitation, we generated triple-negative breast cancer mice that develop multifocal, asynchronous tumors along a continuous luminal-to-basal transdifferentiation trajectory. Ordering spatial transcriptomes from 100+ ducts along this trajectory reveals the spatiotemporal dynamics and molecular mechanisms of TME remodeling, identifying early stromal programs and signaling axes as potential targets for disease interception before establishment of invasive niches. The stromal spatiotemporal remodeling dynamics are conserved across genetically heterogeneous patient-derived xenograft models and treatment-naïve human TNBC cohorts, supporting the translational relevance of our findings. Our study shows that temporal ordering of spatially-resolved disease snapshots unravels some of the molecular “forces” that, starting from the cell-of-origin, propel cells/microenvironments along a disease trajectory.

## Introduction

Solid tumors are complex ecosystems where malignant cells are embedded in a specialized extracellular matrix (ECM) and interact with surrounding stromal cells. During tumorigenesis, cell-cell and cell-ECM interactions initiate and sustain the changes in cellular composition and molecular phenotypes that ultimately remodel homeostatic tissues towards a tumor-permissive microenvironment^1,2^. Given their critical role in tumor progression, interactions within the tumor microenvironment (TME) have emerged as an attractive therapeutic target across diverse tumor entities^3–5^.

While single-cell RNA sequencing has transformed our understanding of tumor and stromal molecular phenotypes^6,7^, the lack of spatial context hinders the systematic mapping and understanding of cellular interactions in the TME. Pioneering analyses based on H&E morphology^8^ and classical antibody staining^9,10^ – alongside recent spatial omics approaches - have revealed that specific TME architectures correlate with tumor progression^11,12^ and response to therapy^5,13,14^. Very recently, high-resolution spatial transcriptomic (ST) technologies have enabled comprehensive gene expression profiling within intact tissue sections at single-cell and even subcellular resolution^15,16^. By capturing cell–cell and cell–ECM interactions in their native spatial context, these approaches have provided further insights into the mechanisms that stabilize multicellular niches or drive tumor progression^17^. It has been proposed that these and other spatially resolved data hold significant potential for informing mechanism-based, personalized therapeutic strategies in clinical oncology, including interception of disease trajectories^2,18,19^.

Nevertheless, spatial omics only provide a static snapshot, limiting insight into which interactions coordinate TME remodeling dynamics over time. Moreover, patient tissues are often late-stage and heterogeneous, reflecting variable genetics, disease trajectories, and treatment histories. In contrast, *in vitro* systems (*i.e*. organoids and 2D cultures) and genetically engineered mouse models enable studying tumorigenesis along genetically-defined trajectories. While *in vitro* systems may not fully recapitulate immune and stromal components of the TME^20^, mouse models can reveal the dynamic interplay between epithelial transformation and TME remodeling within native, immunocompetent tissues^21^.

Even in mouse models, collecting tumor samples to profile early tumorigenesis can be challenging. For example, if tumor development occurs as a single lesion and several weeks after oncogenic induction, early lesions – being particularly small – can be difficult to localize. Furthermore, numerous animals would need to be analyzed in order to reach adequate spatiotemporal resolution. Therefore, we set out to design a mouse model that would combine (1) a relation to an aggressive human cancer where the TME represents an attractive therapeutic target, such as triple-negative breast cancer (TNBC)^22–27^, (2) genetically-defined tumor development to facilitate ordering individual tumors along a shared trajectory^21,28,29^, (3) fast tumor development minimizing the impact of somatic mutations and (4) genesis of multiple asynchronous tumors within the same gland to facilitate high-resolution, spatiotemporal TME profiling.

We reasoned that such a model could be generated by introducing three oncogenic mutations, commonly altered in human TNBC ^30–33^, activating p53, PI3K, and WNT pathways into luminal epithelial cells of the mammary gland. This combined oncogene activation jumpstarted tumorigenesis, inducing multiple, spatially-distinct tumors within the same gland and enabling the parallel analysis of dozens of healthy ducts, early and advanced tumors and their microenvironments in individual samples. Using high-resolution, unbiased spatial transcriptomics (Open-ST^34^), we generated a unique dataset encompassing 100+ ducts, which we complemented with single-nucleus RNA sequencing from independent animals. Ordering individual ducts along a shared tumor progression trajectory revealed TME remodeling dynamics from tumor initiation to tumor invasion. At the tumor-stromal interface, stromal phenotypes were organized in multicellular niches, which were dynamically remodeled closely following changes in intraductal tumor phenotypes and the emergence of transient cell–cell interactions.

Our spatiotemporal analysis identified a sequential, coordinated remodeling in which early TGF-β2 signaling from the cell-of-origin aligns with the emergence of inflammatory cancer-associated fibroblasts (iCAFs), which in turn are followed by the accumulation of myofibroblastic CAFs (myCAFs) at the invasive tumor border. Co-transplantation of tumor cells in syngeneic mice confirmed the ability of myCAFs to steer tumor trajectories towards invasive phenotypes and demonstrated that perturbation of early signaling axes can alter the subsequent trajectory of stromal remodeling.

Finally, the stromal remodelling dynamics identified here were conserved in response to human TNBC cells with different mutation profiles using patient-derived xenografts (PDX) models and independent cohorts of treatment-naive TNBC patients, supporting their translational relevance for targeting the TME in human TNBC.

Altogether, mapping cell-cell interaction dynamics in space and time provided insight into the drivers of stromal remodeling and identified therapeutic candidates for disease interception.

## Results

### A murine model to study TME remodeling dynamics in TNBC

To model TME dynamics in TNBC, we generated murine models conditionally activating *Pik3ca^H1047R^* and *Trp53^R172H^* oncogenic alleles^35,36^, together with an oncogenic, stabilized form of β-catenin (*Ctnnb1^Δex3^*)^37^ in the mammary epithelium. Oncogenes were activated in luminal epithelial cells using the Cre recombinase under control of the *Wap* promoter (WAPiCre)^38^. Recombined cells were co-labeled with enhanced yellow fluorescent protein (YFP) via a *Rosa26*^eYFP^ reporter allele^39^, allowing tumor cell tracing (**Fig. 1A**).

**Figure 1.**
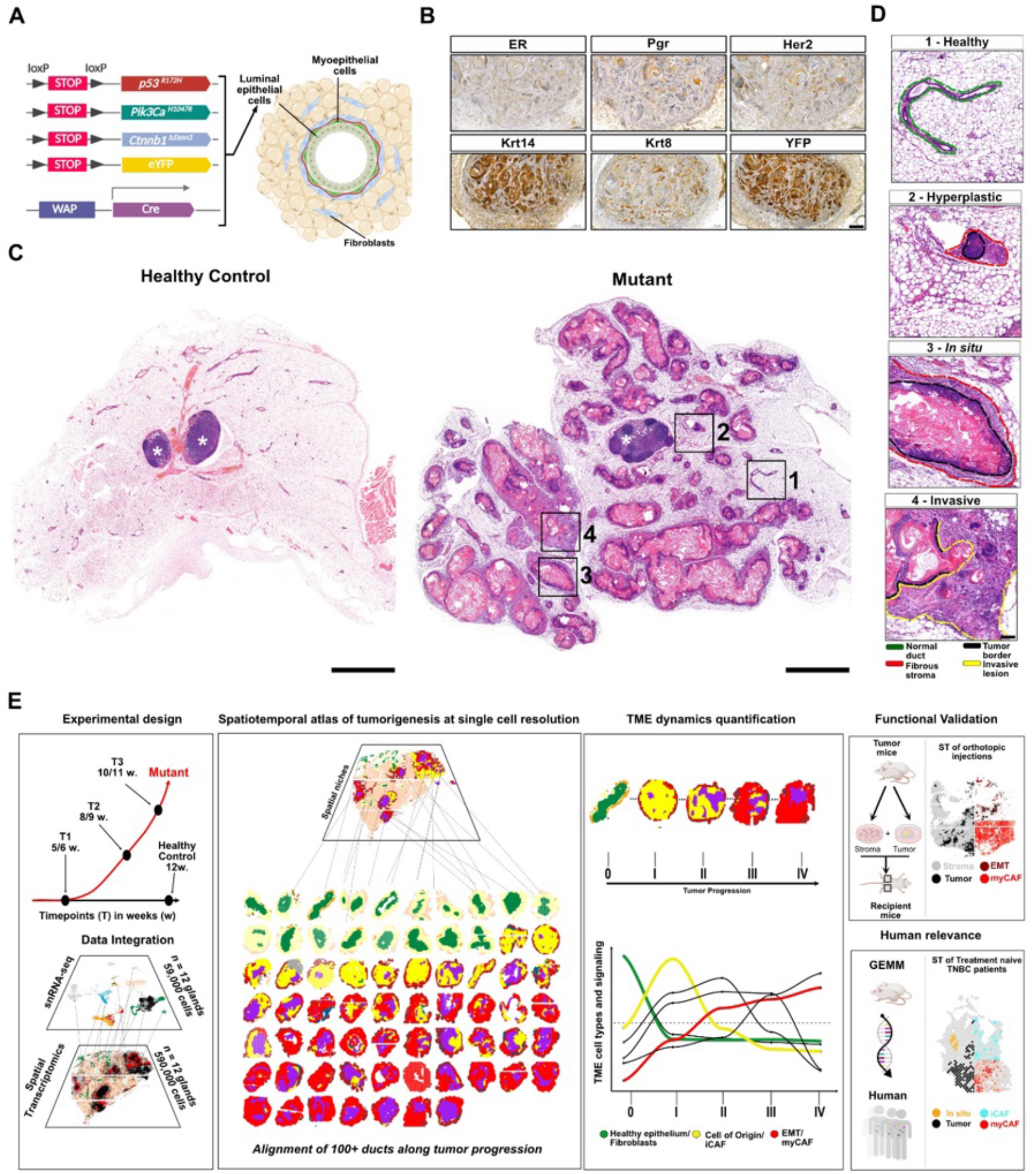
Multiple asynchronous tumors arise in a triple-negative breast cancer (TNBC) model. (A The TNBC mouse model. Using a WAP-iCre system, cells in the luminal epithelium of the mammary glands express three oncogenic drivers (mutant *Trp53^R172H^*, *Pik3ca^H1047R^*, and a stabilized form of β-catenin (*Ctnnb1^Δex3^*). Each of the pathways are frequently activated, sometimes jointly, in human patients. Recombined cells are tagged with YFP. (B) Tumors are triple-negative and basal-like at study endpoint. Immunohistochemistry staining of mammary tumors at 9 postnatal weeks. Observed: low levels of the hormone receptor markers ER, PR, Her2 and luminal keratin Krt8; high level of basal keratin, Krt14 and recombined cell reporter, YFP. Additional replicates in SFig. 1C Scale bar: 200 µm. (C) Multiple tumors arise asynchronously in individual glands. Hematoxylin and eosin (H&E) staining of abdominal mammary glands from mutant and control animals at postnatal week 8. White stars: intramammary lymph nodes. Scale bars: 1 cm. (D) Ducts in individual glands co-exist at different stages of transformation. Higher magnification H&E images from the mutant glands shown in (C). Scale bar: 100 µm. (E) Framework to study tumor progression and TME remodeling: 1) Tumor growth is detectable after 6 postnatal weeks. Spatial Transcriptomics and snRNA-seq were performed on healthy control and three tumor stages (T1–T3) spanning early to advanced. 2) Ducts were computationally segmented to resolve events at the duct– stromal interface. 100+ spatially sequenced ducts were computationally ordered along tumor progression. 4) Cellular, ligand-receptor, and ECM dynamics were analyzed across stages of tumor evolution 5) Functional validation of the role of myCAF in TME remodeling and promoting the emergence of invasive tumor phenotypes 6) The stromal spatiotemporal remodeling dynamics are recovered in human TNBC data.

Activation of single oncogenes led to tumor development in all animals between 25 and 41 postnatal weeks (**SFig. 1A**). Instead, control animals, which carried all oncogenic alleles but lacked WAPiCre, never developed tumors. In triple-mutants, combined oncogene activation anticipated tumor development. Analysis of gland weight identified the first evidence of tumor onset at 5 postnatal weeks, with fast tumor growth between 5 and 10 weeks when animal welfare endpoints required termination (**Fig. 1B, SFig. 1B, C)**. At study termination, tumors featured a TNBC immunohistochemistry profile, lacking estrogen, progesterone and HER2 receptors (**Fig. 1B, SFig. 1D**). Accordingly, advanced lesions exhibited basal features, namely high basal (Krt14) and low luminal (Krt8) markers, and histopathological features consistent with squamous metaplastic carcinoma, including abundant intraluminal keratinization and invasion into a desmoplastic stroma. Instead, YFP detection was restricted to Krt8+ cells in 5-week-old mice (**Fig. 1B, S****Fig. 1**E), confirming tumor initiation in the luminal compartment and suggesting the acquisition of basal features with tumor progression. At study termination, whole-gland hematoxylin and eosin (H&E) staining revealed multiple ducts at various stages of transformation separated by the adipocyte-rich stroma typical of healthy glands (**Fig. 1C**). In the same gland, ‘healthy’ ducts lined by a thin layer of fibroblasts (1) could still be detected together with ‘hyperplastic’ ducts (2), ‘*in situ*’ keratinized ducts confined by a fibrotic layer (3) and ‘invasive’ lesions where nests of tumor cells infiltrated the fibrotic stroma (4) (**Fig. 1D**). Electron microscopy further confirmed progressive remodeling of the tumor–stroma interface. Whereas healthy ducts were surrounded by a continuous basement membrane enclosing cuboidal epithelial cells, advanced lesions displayed loss of basement membrane integrity together with elongated fibroblasts and infiltrating myeloid cells surrounding tumor cells with irregular nuclear morphology and keratin deposition (**SFig. 1F**). Consistent with locally invasive disease, whole-gland immunofluorescence (IF) highlighted nests of tumor cells in the intramammary lymph node when included in the same tissue section (**SFig. 1G**).

In summary, we generated a fast, reproducible and aggressive model of TNBC to study the dynamics of tumor and microenvironment remodeling from healthy ducts to invasive lesions.

### Spatiotemporal study of TME remodeling at single cell resolution

To study tissue remodeling throughout tumorigenesis, we combined snRNAseq with high-resolution, genome-wide (polyA-based) spatial transcriptomics (Open-ST^34^) on 24 abdominal mammary gland samples across one control and three tumor timepoints. The sampling strategy was designed to span the entire disease course, from the earliest detectable lesions at tumor onset (T1, 5-6 weeks), through intermediate tumor expansion (T2, 8-9 weeks), to the experimental endpoint when tumors reached the maximum size permitted by the animal protocol (T3, 10-11 weeks), as reflected by the progressive increase in mammary gland weight (**Fig. 1G, SFig. 1H**).

Unbiased clustering of 58,742 single-nuclei revealed 15 cell types (**SFig. 2A-B**) spanning epithelial, stromal, and immune compartments (**Fig. 2A, STable1**). The epithelial compartment included luminal, hormone sensing and myoepithelial cells as well as transcriptionally distinct tumor cell states not detected in control samples. Stromal populations consisted of fibroblasts, pericytes, endothelial cells, and adipocytes. The immune compartment included both lymphoid (B and T cells) and myeloid lineages, including macrophages, monocytes, plasmacytoid and migratory dendritic cells.

**Figure 2.**
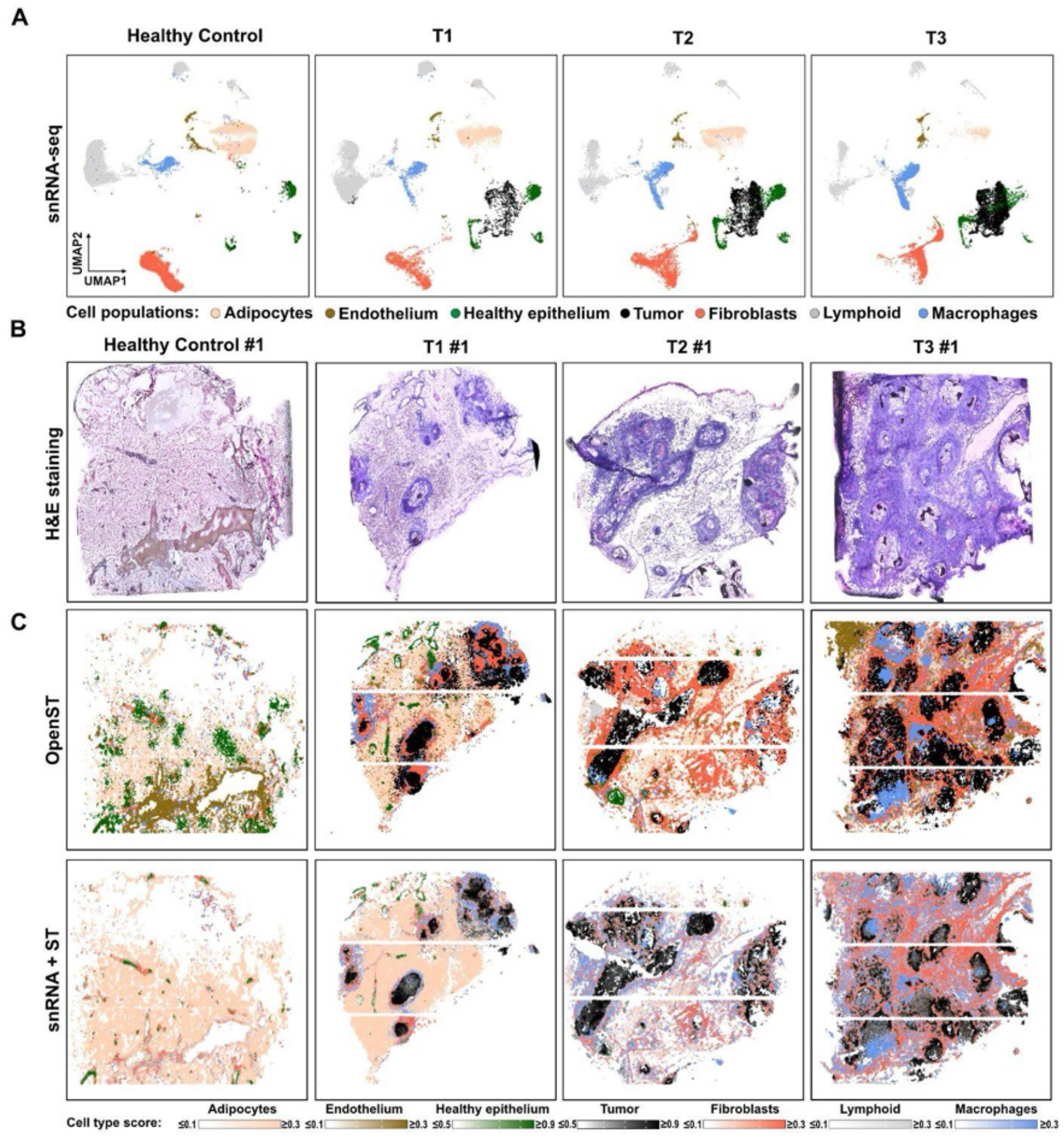
Spatiotemporal quantification of tumorigenesis at single cell resolution. (A) Epithelial and stromal cell types are remodeled in the tumor microenvironment (TME). Uniform manifold approximation and projection (UMAP) of single-nucleus RNA-seq data isolated from healthy and mutant mammary glands along tumor progression (Timepoints T1–T3). RNA from *N* = 58,742 nuclei was sequenced (12 samples, 3 biological replicates/timepoint, Methods). Cell types in SFig. 2A. (B) Spatial transcriptomics captures the progressive remodeling of TME molecular architecture. Top: H & E images of tissue slices from glands acquired for spatial transcriptomics. Bottom: Clustering of the genome-wide, high-resolution spatial transcriptomics data (Open-ST). RNA from *N* =587,288 pseudocells was sequenced (12 samples, 3 biological replicates/timepoint, Methods). Additional replicates in SFig. 2C. Scale bars: 500 µm. (C) Top: Clustering of the genome-wide, high-resolution spatial transcriptomics data (Open-ST). RNA from N =587,288 pseudocells was sequenced (12 samples, 3 biological replicates/timepoint, Methods). Additional replicates in SFig2C. Scale bars: 500 µm Single-nucleus RNA-seq mapped to spatial transcriptomics. Bottom: As in top but colored by “Robust Cell Type Deconvolution” scores (RCTD) for selected cell types captured by single-nucleus RNA sequencing. RCTD is a supervised learning approach to decompose cell type mixtures in pseudocells using profiles learned from single-nuclei RNAseq after correcting for platform-specific effects. Each cellular population gets a probabilistic score (0 to 1) in each spatial location and scores in each spatial location sum up to 1.

To investigate TME molecular organization, we utilized our ST data. Open-ST provides genome-wide transcriptomic profiling at subcellular (0.6 μm) spatial resolution^34^. To enable robust identification of cell states, we aggregated neighboring barcodes into ∼120 μm² hexagonal pseudocells (Methods). Unbiased clustering of 587,288 pseudocells identified regions characterized by epithelial, stromal, and immune marker gene expression. The resulting molecular architecture closely matched the cell populations identified independently by snRNAseq (**Fig. 2B**, **SFig. 2C**).

To map snRNAseq populations to their tissue locations, we leveraged Robust Cell Type Deconvolution^40^ (**Fig. 2C**). RCTD probabilistically assigns cell-state compositions to each spatial location by integrating the single-nucleus reference with spatial gene expression while accounting for platform-specific differences. Our integrated analysis captured the progressive remodeling of gland architecture along tumorigenesis. While control samples only featured healthy ducts surrounded by a thin layer of fibroblasts and an adipocyte-rich stroma, gene expression profiling confirmed that healthy ducts co-existed with multiple transformed ducts in tumor samples. From T1 to T3, transformed ducts increased in size and were surrounded by abundant fibroblasts and macrophages, which replaced adipocytes in the periductal stroma. Macrophages also accumulated in proximity and inside transformed ducts. Of note, none of the ducts in ST sample T1#3 showed evidence of tumor transformation, suggesting that tumor initiation may not be fully synchronized across all animals (**SFig. 2C**). Surprisingly, while lymphocytes composed ∼30% of dissociated cells, they were rare in the periductal stroma. In fact, analysis of individual single-nuclei samples revealed that several immune and stromal populations were either always present or absent together (**SFig. 3A**). Compatible with intramammary lymph nodes being at times captured upon tissue dissociation, these immune and stromal populations mapped exclusively to the intramammary LN captured in ST sample T2#3 (**SFig. 3B**). Consistent with an immune desert TME, CD3 gene expression and T cell RCTD scores both localized to the intramammary lymph node (**SFig. 3C-D**) and CD3 immunostaining confirmed that T cells were largely absent from the tumor-associated stroma (**SFig. 3E**). Therefore, integrating ST and snRNA-seq data allowed to map the organization of cellular populations in the TME and excluded potential confounding effects brought about by either approach alone.

**Figure 3.**
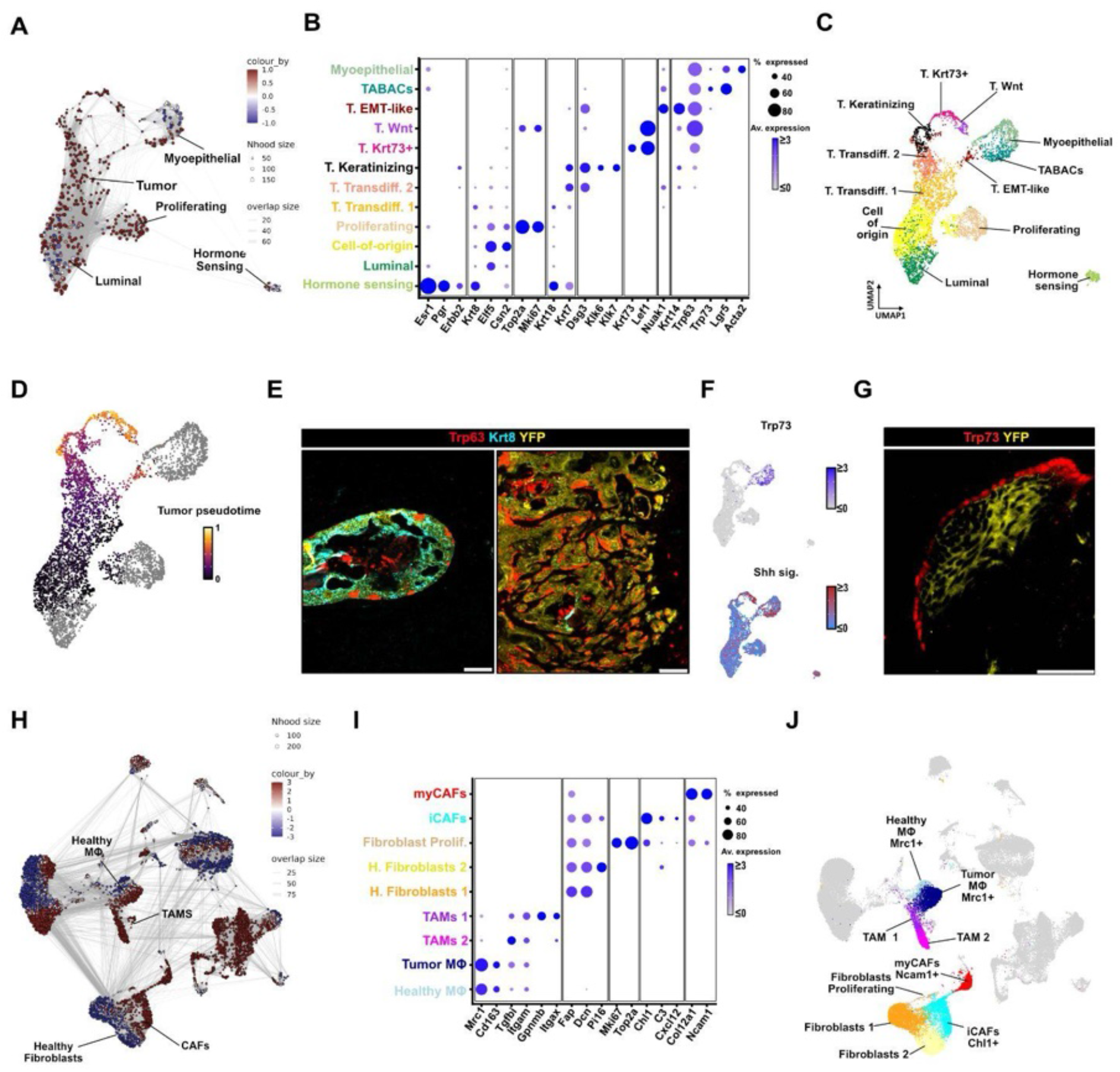
Epithelial and stromal cells undergo extensive remodeling in the TME. (A) Epithelial cells are remodeled after oncogene activation. Differential abundance analysis (“MILO”, Methods) was performed on the epithelial single-nucleus RNA-seq dataset to quantify changes in neighborhood abundance between healthy and tumor-bearing (T1–T3) samples. Each node represents a neighborhood of transcriptionally similar cells in the k-nearest-neighbor graph. Nodes are colored by the log fold change in neighborhood abundance between tumor and healthy samples. Red/blue: enrichment in tumor-bearing/healthy samples. Node size is proportional to the number of cells in each neighborhood, and edge thickness reflects neighborhood overlap. *N* = 5,428 nuclei integrated from 12 samples (3 biological replicates/timepoint). Color scale clipped between 1 and −1. (B) Marker gene expression supports epithelial annotations. Dot size: percentage of cells (≥20% shown), for each cell state, where ≥1 transcript of a specific gene was detected. Color: Z-scores of mean gene expression values per cell state. T: Tumor, Av: average. (C) As in (A, D, F) but colored by cluster annotations (see B). (D) Pseudotime captures tumor progression after oncogene activation. As in (A), but colored by pseudotime scores computed by probabilistic modelling of tumor cell trajectories anchored to a Cell-of-origin (“Palantir”, Methods). Grey: Luminal and myoepithelial cells. Hormone sensing not shown. (E) Tumor cells (YFP) down/up-regulate luminal/basal markers during tumor progression (Krt8 and Trp63, respectively). IF staining of early (left) and invasive (right) lesions. Scale bars: 50 µm. (F) Markers of Tumor-associated basal cells (TABACs). As in (A, C) but colored by Z-scores for Trp73 expression (top) and AU Cell enrichment for the ‘Hedgehog Signaling Pathway’ gene set (bottom; Methods). (G) TABACs (Trp73^+^) form a layer of non-recombined cells around transformed cells (YFP^+^). IF staining of an intermediate lesion. Scale bars: 50 µm. (H) Stromal cells are remodeled in the TME. Differential abundance analysis of cell states was performed as in (A) to quantify neighborhood abundance changes between healthy and tumor-bearing (T1–T3) samples. *N* =58,742 nuclei from 12 samples (3 biological replicates/timepoint). Color scale clipped between 3 and −3. (I) Dot plot as in (B) but of marker genes supporting stromal cell state annotations. (J) Stromal molecular phenotypes. As in (H) but colored by cell state annotations (see (I)).

### Epithelial and stromal cells undergo extensive remodeling in the TME

To study the phenotypic remodeling of cellular populations in the TME, we proceeded to investigate epithelial and stromal transcriptional states captured by snRNAseq throughout tumorigenesis.

Unbiased clustering of 5,428 epithelial cells identified 12 populations, which we annotated based on their enrichment in CTRL vs mutant samples (**Fig. 3A, SFig. 4A**), the expression of canonical marker genes (**Fig. 3B**, **STable2**) and reference-based label transfer from a murine healthy and transformed mammary gland atlas^41^ (**SFig. 4B**). While CTRL samples featured hormone-sensing (*Esr1+ Pgr+ Erbb2+*), luminal (*Elf5*+), and myoepithelial cells (*Acta2*+), multiple populations were only detected in mutant samples despite data integration (**SFig. 4C**). *Wap* expression, which drives oncogene activation, was specific to *Csn2*+ alveolar luminal cells (**SFig. 4D**). This population, which we annotated as ‘cell-of-origin’, was already present but rare in CTRL samples and expanded during tumorigenesis. Again, label transfer scores and *Krt8* expression confirmed tumor initiation in the luminal compartment. Instead, further tumor populations showed decreasing levels of luminal and increasing level of basal markers (i.e. *Krt14* and *Trp63*). Despite their shared genotype, basal-like tumor populations acquired diverse phenotypes, distinguished by keratinization, Wnt signaling and EMT (epithelial-to-mesenchymal transition) signatures (**Fig. 3C, SFig. 4E**). Pseudotime analysis starting from the cell-of-origin (Methods) ordered tumor cells along continuous trajectory and revealed divergent endpoints, which corresponded to the activation of different molecular pathways in basal-like tumor cells (**Fig. 3D**). Consistent with such luminal-to-basal (L2B)^42,43^ transition, IF showed widespread Krt8+/YFP+ luminal tumor cells in hyperplastic lesions, while YFP+/Trp63+/Krt8- basal tumor cells characterized invasive lesions (**Fig. 3E**, **SFig. 4F**). In contrast to hormone-sensing cells, which persisted as a stable, non-transformed population, myoepithelial cells were remodeled in mutant samples. Compared to myoepithelial cells isolated from CTRL samples, tumor-associated basal cells (TABACs) upregulated *Trp73* expression and sonic Hedgehog pathway activity (**Fig. 3F**), downregulated *Acta2* while retaining the expression of canonical basal markers (i.e. Trp63) (**SFig. 4G**). Consistent with non-recombined, tumor-reprogrammed myoepithelial cells, IF identified a monolayer of YFP-/Trp73+ cells surrounding tumor lesions (**Fig. 3G, SFig. 4H**). Altogether, oncogene activation gradually transformed luminal alveolar cells into divergent basal-like tumor states and led to myoepithelial cell remodeling into TABACs.

**Figure 4.**
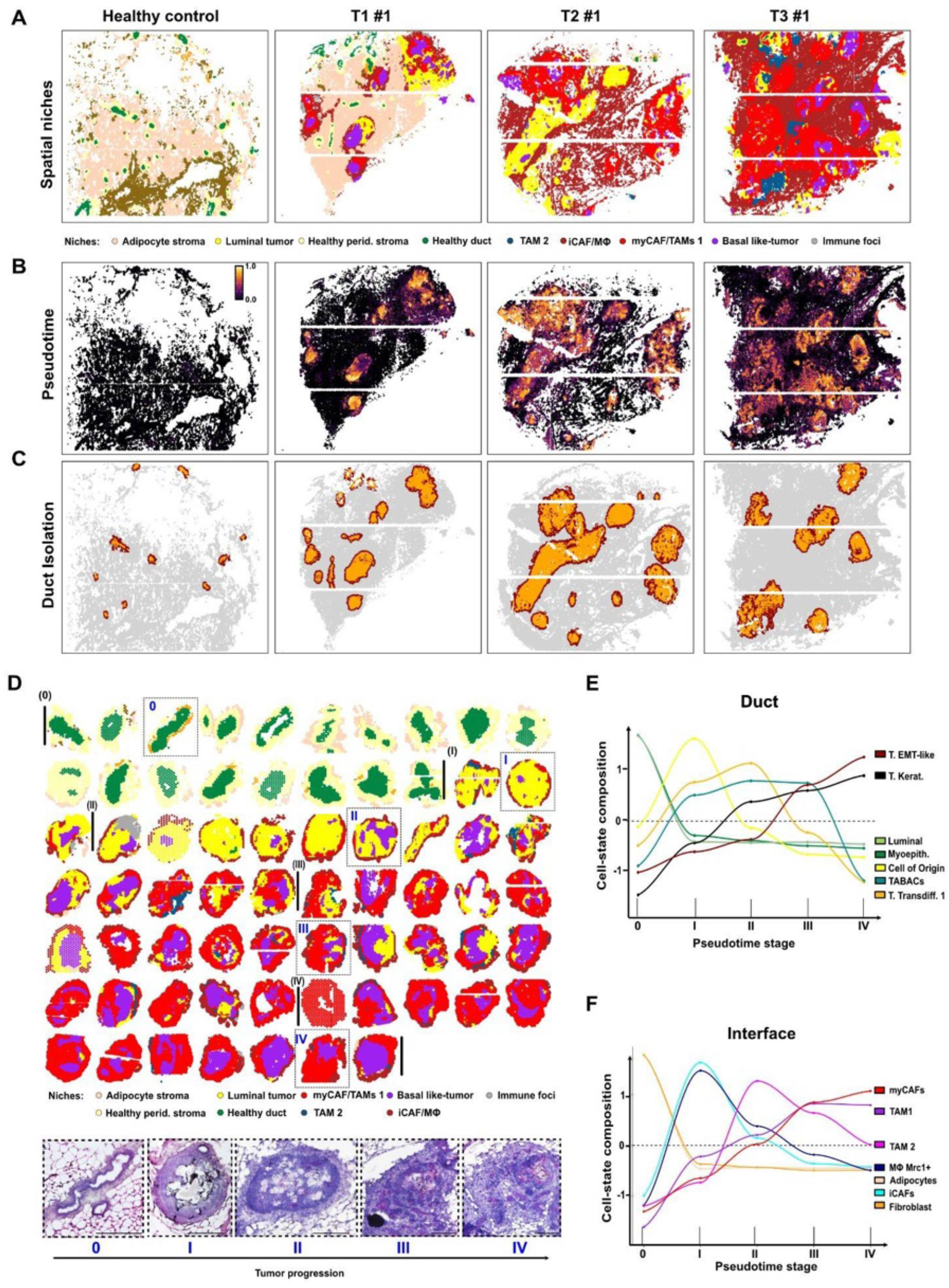
Spatiotemporal dynamics of tumor-stromal interface remodeling. (A) Spatial mapping of epithelial and stromal niches during tumorigenesis. Open-ST samples (as in Fig. 2B) are colored by niche assignment. Average RCTD cell-state compositions per cellular neighborhood (k=30) were grouped into recurrent spatial niches by unsupervised clustering (see SFig. 4A-D, Methods). Additional replicates in SFig. 6D. (B) Spatial mapping of tumor pseudotime scores (see Fig. 3C) inferred by label transfer from the single-nucleus epithelial trajectory to Open-ST data. As in (C) but colored by pseudotime label transfer scores (Methods). Higher pseudotime values correspond to progressively advanced epithelial tumor states. Additional replicates in SFig. 6E. (C) Manual segmentation of individual epithelial ducts and adjacent periductal interface regions used for duct-resolved analyses. Intraductal regions are shown in orange and surrounding duct–stromal interface regions are shown in yellow. Colors indicate duct regions and do not represent different biological duct classes. Ducts were manually segmented based on epithelial boundaries after RCTD-based spatial mapping; ducts with unclear boundaries or less than 250 cells were excluded. Additional replicates are shown in SFig. 6F. (D) Ordering 100+ ducts along increasing tumor pseudotime reveals a continuous progression from healthy ducts to advanced invasive lesions. Each duct is colored according to its spatial niche composition as shown in (A). Ducts with < 250 pseudocells were removed (44 out of 111). Representative hematoxylin and eosin (H & E)-stained images illustrate the corresponding histopathological progression from normal mammary ducts (stage 0), through in situ (stages I) and keratinizing lesions (stage II-III) and invasive carcinoma (stage IV). (E) Dynamics in epithelial cell-state composition across tumor pseudotime. Curves represent scaled RCTD-inferred abundances of epithelial cell states within intraductal regions, revealing the progressive loss of normal luminal and myoepithelial populations and the emergence of transdifferentiating, keratinizing, and EMT-like tumor states. (F) Dynamics in stromal cell-state composition at the tumor interface across tumor pseudotime. As in (E) but for stromal and immune populations, showing the sequential emergence of iCAFs, tumor-associated macrophages, and myCAFs within the duct-stromal interface during tumor progression.

Besides epithelial cells, we sought to identify which stromal phenotypes emerged in the TME. Compatible with their tissue-resident role, two fibroblast (‘Fibroblasts 1’ and ‘Fibroblasts 2’) and one macrophage population (‘Macrophages/MΦ healthy’) were enriched in CTRL samples (**Fig. 3H, SFig. 5A**) and showed similarity to reference populations in the Bach *et al* dataset (**SFig. 5B**). Conversely, fibroblasts in mutant samples expressed cancer-associated signatures^44^ (**SFig. 5C**), supporting their annotation as inflammatory cancer-associated fibroblasts (iCAFs) and myofibroblastic CAFs (myCAFs). Similarly, two macrophage populations featured the upregulation of a tumor-associated macrophage (TAM) signature whereas a population of Mrc1+, tumor-enriched macrophages (‘Macrophages tumor’) did not express TAM signatures likely representing macrophages being recruited but not yet reprogrammed in the TME. Rather than representing M1 and M2 macrophages, TAM1 and TAM2 were both enriched for the TREM2/SPP1 tumor-infiltrating macrophage signature^45,46^, while ‘Macrophages tumor’ for the FOLR2/LYVE1 tissue-resident one (**SFig. 5D**). Differential gene expression and gene set enrichment analysis further confirmed that ‘Macrophages tumor’ were relatively similar to healthy macrophages compared to TAMs and were enriched for interferon-response pathways (**SFig. 5E-F**). In contrast, TAM1 and TAM2 featured a shift in gene expression and showed enrichment for Hallmark TGFβ signaling with TAM2 further upregulating hypoxia, glycolysis and TNFα/NFκB signaling pathways. Finally, Chl1, Ncam1, Gpnmb emerged as specific markers for iCAFs, myCAFs and TAMs in our model and were used for downstream validations (**Fig. 3I**, **SFig. 5G**, STable3).

**Figure 5.**
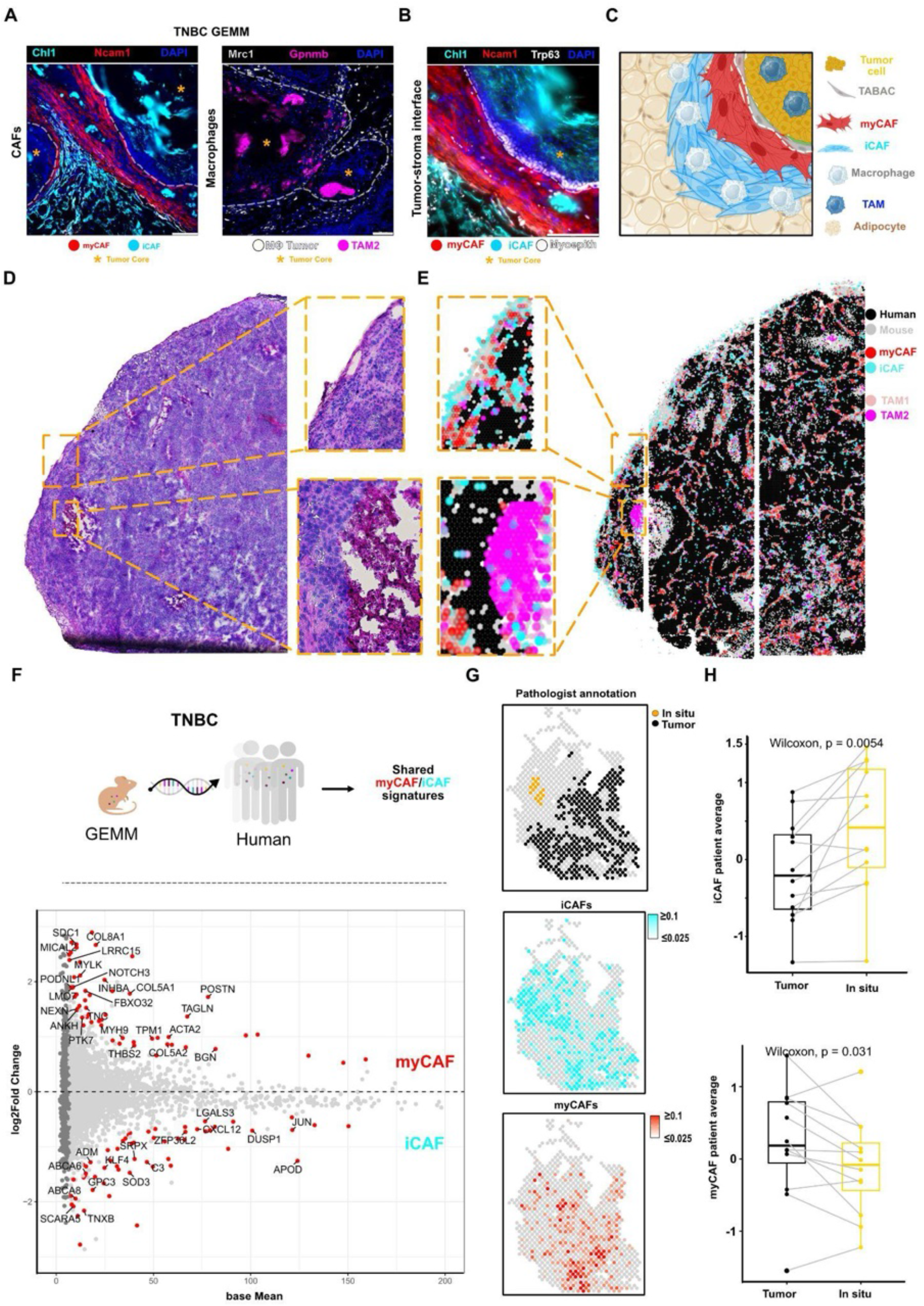
Spatiotemporal stromal dynamics are conserved in human TNBC. (A) Cell type distribution at the duct-stromal interface. IF staining of iCAFs (Chl1⁺), myCAFs (Ncam1⁺), macrophages (Mrc1⁺), and TAMs (Gpnmb⁺) in a T2 8 week tumor. Scale bars: 50 µm. (B) TABACs (Trp63^+^) form a layer between tumor cells (YFP^+^) and myCAFs (Ncam1^+^) interface. IF staining showing the tumor–stroma interface in a primary tumor. Scale bars: 100 µm. (C) Spatial cell types organization at the tumor-stromal interface. (D) H & E staining of the profiled TNBC PDX section (HBCx-14). Zooms indicate the tumor boundary (top) and a keratinizing region (bottom). (E) Species-aware transcript assignment distinguishes human tumor-cell regions from mouse stromal regions. RCTD deconvolution scores localize myCAFs (red) and TAM 1 (purple) to the tumor-stromal interface, iCAFs (cyan) and ‘Macrophages tumor’ (darkblue) to the peritumoral stromal and TAM 2 to the keratinizing region (magenta). Individual plots shown in SFig.8 E-H. (F) Cross-species identification of conserved CAF signatures. Top: Graphical representation. Bottom: Differential expression analysis of CAFs from treatment-naïve human TNBC (Methods). Pseudobulk analysis of single-cell RNA-seq data (Wu et al., 2021) was used to identify human iCAF (bottom) and myCAF (top) marker genes for comparison with the corresponding mouse populations. Red: genes with adjusted p-value < 0.05 across TNBC patients. Labeled: conserved genes with adjusted p-value < 0.05 in both TNBC patients and GEMM, used for spatial mapping in (G) and SFig.9. (G) Spatial projection of conserved stromal signatures in human TNBC. Pathologist’s annotation of in situ and invasive tumor regions (top) together with spatial AU Cell scores of conserved iCAF (middle) and myCAF (bottom) signatures (representative section from patient TNBC7; Methods). Low-resolution spatial transcriptomics data from Wang et al 2024. 11 additional patients shown in SFig.9. (H) Stromal remodeling dynamics are conserved in human TNBC progression. Paired analyses demonstrate iCAF signature enrichment in pathologist-annotated in situ lesions compared to invasive tumor regions within the same section. N = 12 patients. Points indicate patient mean iCAF/myCAF AUCell scores in tumor/in situ regions. Lines connect the same patient. Boxes indicate the interquartile range, center lines the median, whiskers the 1.5× interquartile range. P values: Two-sided Wilcoxon signed-rank.

Overall, single-cell analysis captured the progressive remodeling and the acquisition of diverse molecular phenotypes in transformed glands. While oncogenes drive the remodeling of luminal epithelial cells, the emergence of diverse tumor (**Fig. 3C**) and stromal states (**Fig. 3J**) suggests a role for local interactions in shaping molecular phenotypes in the TME.

### Tumor and stromal states are spatially organized in dynamic multicellular niches

To investigate the organization of tumor and stromal phenotypes in the TME, we mapped single-nuclei populations onto the spatial transcriptomic data using RCTD (**SFig. 6A**).

**Figure 6.**
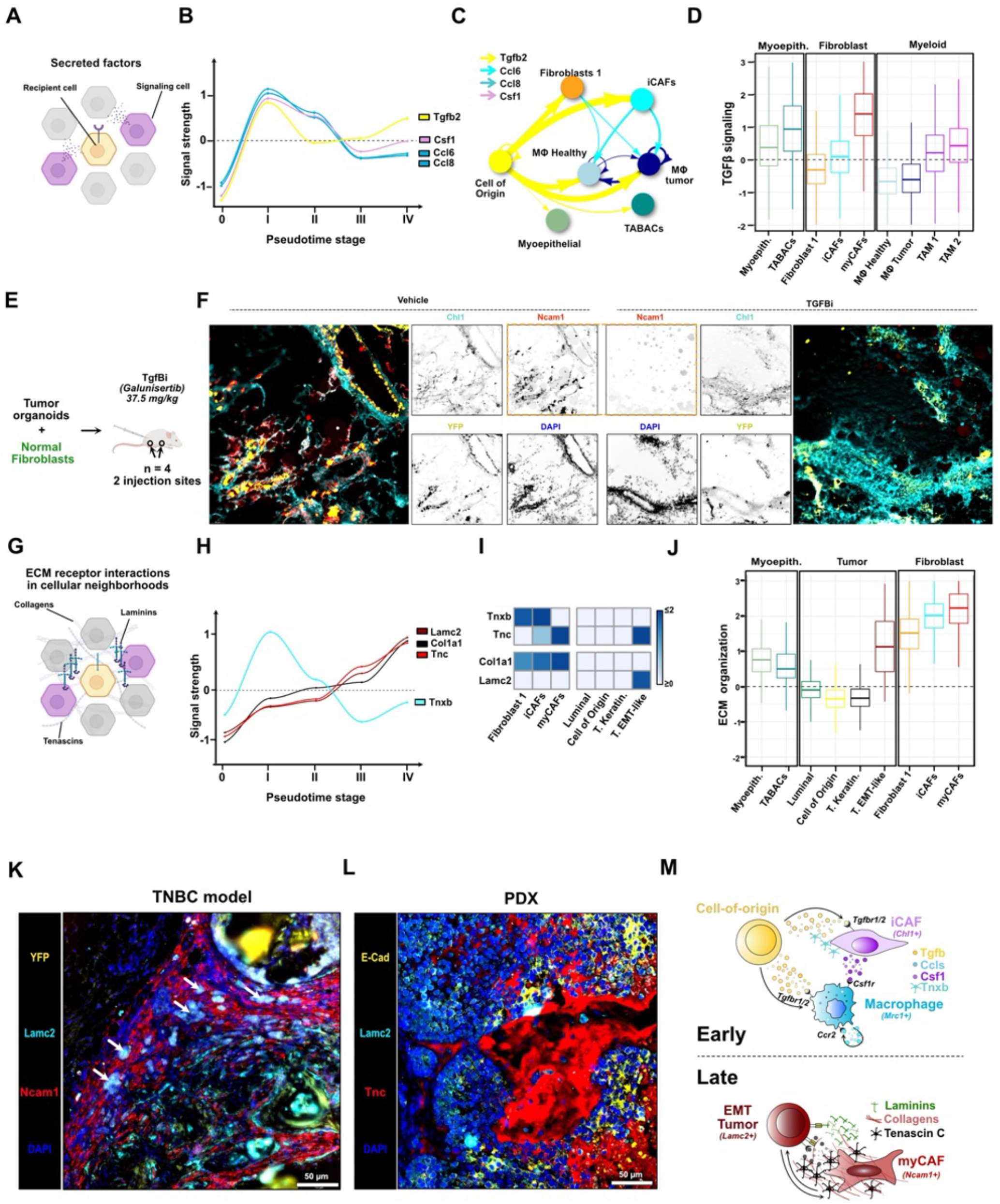
Coordinated and transient signaling axes orchestrate early interface remodeling. (A) Spatial analysis of secreted ligand-receptor interactions in cellular neighborhoods. Ligand spatial activity is computed based on co-expression of ligand and its receptors in neighboring cells (Methods). Pairs of ligand and receptor pairs from CellChat. (B) Transient induction of signaling axes at the early (Stage I) tumor-stromal interface. As in Fig. 5B but showing Z-scores of mean spatial ligand activity, computed for pseudocells at the tumor-stromal interface. Additional Stage I and other stage-enriched ligands shown in SFig. 6A. (C) Tgfb2 signaling from the cell-of-origin is received by early-interface populations. snRNA-seq data is used to identify the directionality of stage-specific interface interactions (see (B)). Arrow width is proportional to geometric of ligand/receptor expression in the sender/receiver populations mapping to the early duct-stromal interface (Stage I). Arrows shown for receptor and ligand expression detected in >30% of cells. (D) Intracellular targets of TGFβ signaling are upregulated in early-interface populations. Box plots showing Z-scores of AU Cell enrichment for HALLMARK ‘TGFβ signaling’ gene set in snRNA-seq populations mapping to the duct-stromal interface. (E) Experimental design for orthotopic co-transplantation. Tumor cells were injected with healthy fibroblasts into the mammary fat pad of immunocompetent FVB/N mice. and treated either with Vehicle or pan TGFβ inhibitor *Galunisertib* (F) TGFβ inhibition affects the Ncam1 myCAFs accumulation around the injected tumor cells. Immunofluorescence merged image and single-channel images for Ncam1, Chl1 YFP and DAPI are shown in treated samples described in (E). (G) Spatial analysis of ECM-receptor interactions in cellular neighborhoods. As in (A) but focusing ECM ligand–receptor pairs. (H) Progressive ECM remodeling at the tumor-stromal interface. As in (B) but showing the dynamics of selected ECM ligands. All stage-enriched ligands are shown in SFig. 10. (I) Heatmap: mean ligand expression across selected populations from snRNA-seq data. (J) myCAFs orchestrate ECM organization. Z-score for AU Cell enrichment of ‘ECM organization’ REACTOME gene set in selected snRNA-seq populations. (K) EMT-like tumor cells invade in a remodeled ECM at the tumor-stromal interface. IF staining in a T3 tumor showing individual Lamc2⁺/YFP⁺ tumor cells (arrows) in proximity to Ncam1⁺ myCAFs. Scale bar: 50 µm. (L) Lamc2⁺ tumor cells infiltrate in a remodeled ECM in PDX samples. IF staining of a PDX sample showing Lamc2⁺/Ecad⁺ tumor cells in a Tnc⁺ stroma. Scale bar: 50 µm. (M) Graphical representation of signaling events in early and late tumorigenesis.

Unbiased clustering of average RCTD deconvolution scores in cellular neighborhoods (S**Fig. 6B**) identified 13 multicellular niches, ranging from tissue-resident to tumor-specific microenvironments (**SFig. 6C**). As a positive control, hormone sensing, luminal and myoepithelial cells colocalized in a specific niche, which we annotated as ‘healthy duct’. Similarly, endothelial cells and pericytes colocalized in the ‘blood vessel’ niche. Vascular cells and adipocytes were enriched in the ‘adipocyte stroma’ and -together with fibroblasts 1- in the ‘healthy periductal stroma’ niches. Instead, Fibroblasts 2 were particularly abundant in the ‘fibrous stroma’ niche. Spatial mapping (**Fig. 4A, SFig. 6D**) confirmed the alignment of multicellular niches with healthy ducts, blood vessels, adipocyte stroma evident from tissue histology (**Fig. 2B, SFig. 2C**) and associated the ‘fibrous stroma’ niche with the gland capsule captured in T1#2 (**SFig. 6D**). Integrating single-nuclei and ST revealed how tumor-remodeled fibroblast and myeloid populations were closely associated in space, with iCAFs and tumor macrophages colocalizing in the ‘iCAF/macrophage niche’ and myCAFs and TAMs 1 in the ‘myCAF/TAMs1 niche’. Instead, TAMs 2 were enriched in a specific ‘TAM 2 niche’. As expected, immune populations mostly colocalized with lymph node-related stromal cell types to the ‘lymph node niche’. Of note, monocytes, B and dendritic cells also colocalized to discrete ‘immune foci’ infrequently detected in tumor samples. Furthermore, the spatial organization of tumor states closely followed their pseudotime. Early states including cell-of-origin and transdifferentiating tumor cells were enriched in the ‘Luminal tumor niche’, while late, basal phenotypes (i.e. keratinizing, Krt63+ and Wnt basal tumor populations) in the ‘Basal-like tumor niche’ mapped to ducts whose enlarged lumen was filled by keratin pearls. Instead, EMT-like tumor cells did not localize to the basal-like tumor niche but rather with TABACs, myCAFs and TAMs 1 in the myCAFs/TAMs 1 niche, compatible with a role for the local microenvironment in promoting invasive tumor phenotypes.

To understand the dynamics of niche remodeling during tumorigenesis, we mapped tumor pseudotime scores in space (Methods). As a control, healthy ducts and the periductal stroma featured near-zero pseudotime, while a range of positive pseudotime scores was detected across transformed ducts (**Fig. 4B, SFig. 6E**). We thus proceeded to segment (**Fig. 4C, SFig. 6F**) 111 individual ducts and sort them along their average pseudotime scores. The independent mapping of tumor pseudotime and multicellular niches highlighted a continuum of duct transformation (**Fig. 4D**), partitioned in 5 stages: ‘Stage 0’ (Average duct pseudotime from 0,0 to 0,15), I (0,16-0,31), II (0,32-0,47), III (0,48-0,62) and IV (0,63-0,78). Remarkably, molecular stages were coupled with histological tumor progression. Stage 0 ducts displayed normal mammary architecture, Stage I ducts corresponded to epithelial hyperplasia and carcinoma in situ, Stage II–III ducts to progressively disorganized keratinizing tumors, whereas Stage IV contained invasive lesions characterized by disruption of ductal architecture and tumor cell infiltration into the surrounding stroma (**Fig. 4D**).

In summary, spatial mapping of epithelial and stromal phenotypes revealed their colocalization in multicellular niches. Pseudotime scoring and duct segmentation then revealed progressive niche remodeling along tumor progression.

### Spatiotemporal dynamics of tumor-stromal interface remodeling

Having ordered 100+ ducts along a shared tumorigenesis trajectory, we proceeded to investigate how the duct-stromal interface was remodeled throughout tumor progression.

Supporting the robustness of spatial mapping, cell state dynamics (**SFig. 7A**) closely followed marker gene expression levels (**SFig. 7B**). Epithelial state transitions in the duct core followed tumor pseudotime defined at the single-cell level (**Fig. 4E**). Namely, luminal and myoepithelial populations characterized Stage 0 ducts (i.e. healthy), while cell-of-origin cells peaked upon tumor initiation (Stage I). With tumor progression, cell-of-origin cells were rapidly replaced by transdifferentiating tumor phenotypes first (Stages II-III), and by keratinizing and EMT phenotypes later (Stages III-IV). TABACs were detected from tumor initiation throughout tumor progression (Stages I-III) but were absent in Stage IV ducts, compatible with the loss of basement membrane integrity in advanced tumors. We then analyzed how stromal phenotypes were remodeled in response to intraductal epithelial changes (**Fig. 4F**). While healthy ducts were surrounded by fibroblasts and adipocytes (Stage 0), the interface was rapidly populated by iCAFs and macrophages upon tumor initiation (Stage I) and by myCAFs and TAMs 1 with tumor progression (Stages II-IV). Instead, TAMs 2 were detected at the tumor-stromal interface only transiently in Stage III tumors. In advanced tumors, however, we observed TAMs 2 accumulation in the duct core. Furthermore, iCAFs and macrophages remained abundant, but were predominantly located in the periductal stroma distal to the myCAF–TAM interface (**SFig. 7C**).

**Figure 7.**
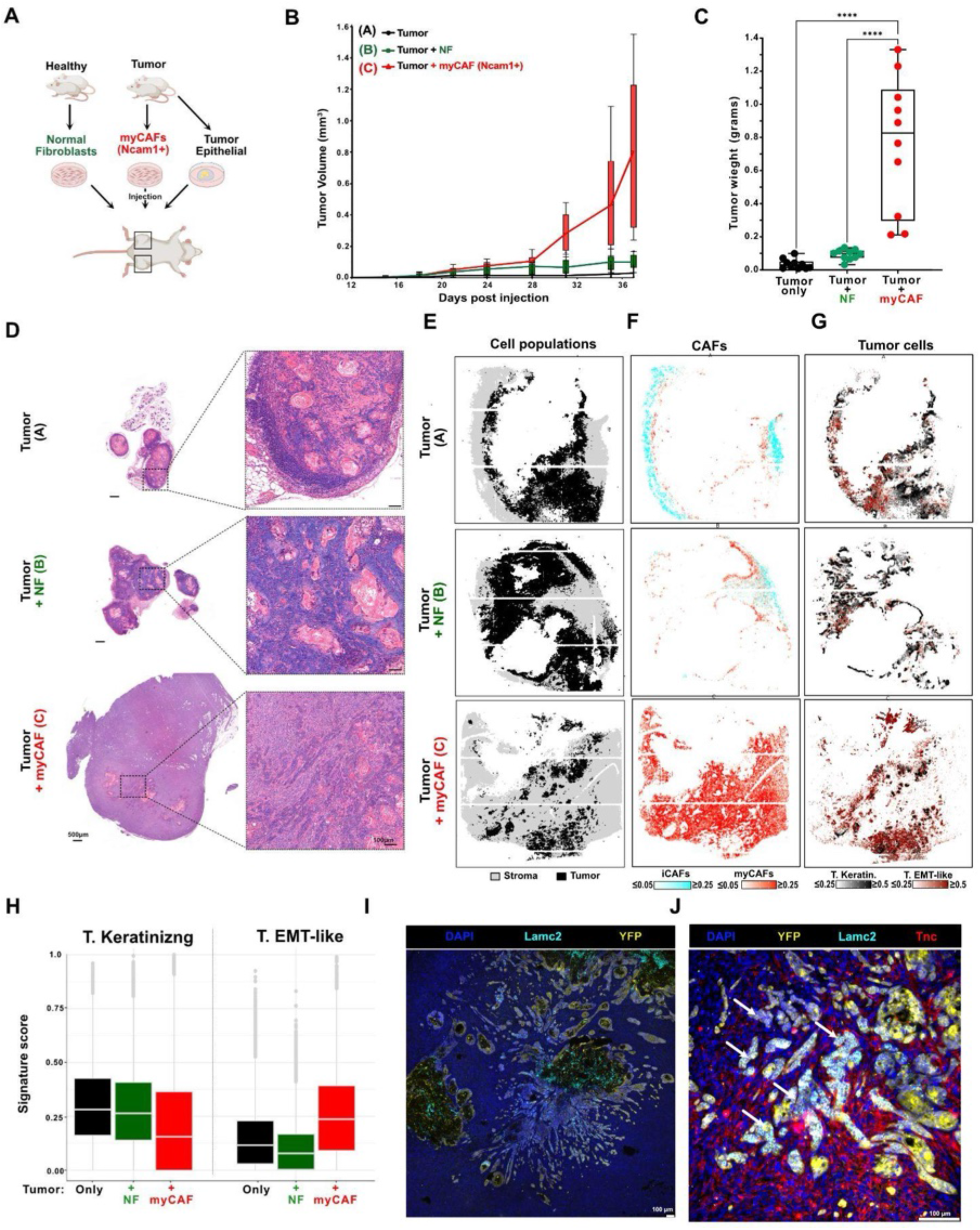
Myofibroblasts are sufficient to promote tumor growth and invasion. (A) Experimental design for orthotopic co-transplantation. Tumor cells were injected alone (Group A), with healthy Ncam1⁻ fibroblasts (Group B), or with tumor-derived Ncam1⁺ myCAFs (Group C) into the mammary fat pad of immunocompetent FVB/N mice. (B) myCAFs promote tumor growth when co-transplanted with tumor cells in healthy immunocompetent mice. Tumor volume was measured daily from day 15 to day 37, the experimental endpoint defined by maximum allowable tumor size. Tumor growth at individual injection sites is shown over time in mm³ (N=10 per group). (C) myCAFs co-injection leads to increased tumor weight at study termination. Fatpad weights (in gram) at the day of study termination for Groups A, B, and C. Bars represent endpoint tumor mass in grams. *N* =10. (D) myCAF co-transplant promotes disorganized tumor histology. Hematoxylin and eosin (H & E) staining of representative tumors from Groups A–C. Scale bars: 500 µm. (E) Spatial transcriptomic (Open-ST) profiling of transplanted tumor derived cells. Spatial mapping (RCTD deconvolution as in Fig. 2C tumor snRNA-seq populations to Open-ST data from one tumor/group. (F) Spatial mapping (RCTD deconvolution as in Fig. 2C and SFig. 5G) of iCAF and myCAFs snRNA-seq populations to Open-ST data from one tumor/group. Immune populations shown in SFig. 7H-J. (G) myCAFs promote invasive tumor phenotypes. As in (C) but showing RCTD deconvolution scores for EMT-like and keratinizing snRNAseq tumor populations. Tumor pseudotime and additional tumor populations shown in SFig. 7K-L. (H) myCAFs shift tumor phenotypes towards EMT. Quantification of keratinizing and EMT tumor phenotypes across conditions. Boxplot of RCTD scores per tumor pseudocells (tumor pseudotime > 0.3), related to (G). (I) EMT-like tumor cells invade in a myCAF-rich, remodeled ECM tumor-stromal interface. IF staining of a Group C tumor. Left: overview of the tumor area showing Lamc2 and YFP magnified region displaying colocalization of Lamc2, YFP, and Tnc. Arrows indicate sites of colocalization. Scale bars: 100 µm (J) Magnified region from I displaying colocalization of Lamc2, YFP, and Tnc. Arrows indicate sites of colocalization. Scale bars: 100 µm

We thus sought to quantify distances of stromal phenotypes to the duct center along tumor progression (Methods). This confirmed how stromal remodeling occurred along a reproducible spatiotemporal gradient: first iCAFs and macrophages were recruited and then populated the distal periductal stromal with tumor progression as myCAFs and TAMs 1 replaced them at the duct border (**SFig. 7D**). On the other hand, the distance of TAMs 2 to the duct border increased with tumor progression, both towards the duct center and in the distal periductal stromal, compatible with their localization to TAM 2 niches. To confirm the remodeling of stromal states, we analyzed their dynamics in individual cellular neighborhoods using phase diagrams (Methods). Paired abundances at the tumor-stromal interface revealed the progressive transition from a macrophage-high/iCAF-high state in tumor initiation (Stage I) towards a myCAF/TAM rich state during tumor progression (Stages II-III) (**SFig. 7E**). Together, these analyses demonstrate that epithelial evolution and stromal remodeling proceed through highly coordinated spatial transitions that can be quantitatively reconstructed by ordering individual ducts along a shared tumor trajectory.

### Stromal remodeling dynamics are conserved across mouse and human TNBC

To validate the stromal states identified by single-nucleus and spatial transcriptomics, we first performed immunofluorescence on primary tumors. Consistent with the spatial atlas, Chl1+ iCAFs localized to the outer stromal compartment in an intermediate tumor gland, whereas Ncam1+ myCAFs accumulated immediately adjacent to the tumor border (**Fig. 5A, SFig. 8A**). Similarly, Gpnmb+ TAMs localized preferentially within the tumor-associated stroma, while Mrc1+ macrophages were distributed more broadly throughout the surrounding tissue.

High-resolution imaging of the tumor–stroma interface further demonstrated that Ncam1+ myCAFs directly surrounded Trp63+ basal cells, whereas Chl1+ iCAFs occupied the outer stromal compartment (**Fig. 5B, SFig. 8B**), validating the spatial organization identified by spatial transcriptomic analyses (**Fig. 5C**).

Patient-derived xenograft (PDX) models complement mouse models by preserving key molecular and histological features of human tumors, offering a direct link to clinical heterogeneity and therapeutic response ^47,48^. We therefore investigated whether tumor cells from treatment-naïve TNBC patients would elicit a similar stromal reaction using PDX models^47^. Human and mouse transcripts could be readily distinguished by OpenST, allowing simultaneous investigation of tumor and stromal compartments (**SFig. 8E**). RCTD mapping of mouse-derived stromal states in the PDX microenvironment identified spatially segregated iCAFs and myCAFs, closely recapitulating the organization observed in the GEMM (**Fig. 5D-E, SFig. 8E-H**). Likewise, TAM1 and TAM2 occupied distinct regions of the tumor microenvironment, supporting conservation of macrophage heterogeneity across independent tumor models.

IF staining of 9 additional PDX models further confirmed that Ncam1⁺ myCAFs tightly wrapped around PDX tumors (**SFig. 8I**) including but not limited to lines harboring the same genetic alterations (**STable4**). Chl1⁺ iCAFs and Mrc1+ macrophages colocalized in the peritumoral stroma (**SFig. 8C,D**).

Hence, the remodeling of the tumor-stromal interface was shared between primary mouse and PDX tumors regardless of tumor genetic drivers.

To determine whether mouse stromal programs are also present in human TNBC, we analyzed two independent human TNBC cohorts. We analyzed single-cell RNA-seq data from treatment-naïve TNBC patients^25^ to derive human CAF signatures while accounting for inter-patient heterogeneity. This identified 58 iCAF and 57 myCAF marker genes in human TNBC through pseudobulk differential expression across 11 patients (Methods). Comparison with the corresponding mouse populations revealed substantial cross-species conservation, with 16 conserved iCAF markers and 23 conserved myCAF markers shared between mouse and human datasets (**Fig. 5F**, Fisher’s Exact test odds ratio= 5.28 p=0.000001).

To investigate whether conserved stromal programs exhibit the spatiotemporal organization predicted by our mouse trajectory, we analyzed an independent cohort of 92 treatment-naïve human TNBC samples profiled with low-resolution spatial transcriptomics with pixel-level annotations of the matched HE image ^26^. 12 tumors contained both pathologist-annotated carcinoma in situ and invasive regions within the same tissue section, enabling paired comparisons while controlling for inter-patient variability. Spatial mapping of conserved human–mouse CAF signatures confirmed the known enrichment of myCAF signatures in invasive tumor regions (P = 0.031, paired Wilcoxon test) and revealed the enrichment of iCAF signatures (P=0.0054) in pathologist-annotated in situ lesions (**Fig. 5G-H**, **SFig. 9**).

Together, these analyses demonstrate that both the molecular identity of iCAFs and myCAFs and the directionality of their spatial remodeling during tumor progression are conserved between treatment-naïve murine and human TNBC. This prompted us to investigate which communication events underlie such stromal dynamics.

### Coordinated and transient signaling axes orchestrate early interface remodeling

To identify which cell–cell interactions orchestrate tumor–stromal interface remodeling, we quantified the co-expression of known ligand-receptor (LR) pairs in cellular neighborhoods^49^ (**Fig. 6A**, Methods).

Focusing on 390 secreted interactions, we first identified 49 stage-specific ligands enriched at the duct-stromal interface (**SFig. 10A**). We then quantified LR expression in single-nuclei populations to identify potential senders and receivers of such interactions and reconstruct stage-specific communication networks. Upon tumor initiation (Stage I), our integrated communication analysis revealed the coordinated induction of *Tgfb2*, *Csf1* and *Ccl6* and *Ccl8* signaling at the tumor-stromal interface (**Fig. 6B**). Compatible with early remodeling, ligand activity peaked (Stage I) and quickly dropped following tumor initiation (Stage II-IV). In our single-nuclei data, *Tgfb2* ligand expression was specific to the cell-of-origin, while Tgfb receptor (*Tgfbr1*, *Tgfbr2*) expression was detected across all stromal populations (**SFig. 10B**). Instead, Csf1 signaling was specific to periductal fibroblasts and iCAFs (*Csf1*+) and macrophages (*Csf1r*+). In turn, macrophages acted both as senders and receivers of Ccl6 and= 8 signaling, suggesting a positive feedback loop reinforcing macrophage recruitment^50^ (**Fig. 6C**). After peaking at the early tumor interface (Stage I), Tgfb2 activity decreased with tumor progression (Stages II-III) to increase again in Stage IV. In fact, *Tgfb2* expression was downregulated in intermediate tumor states but then detected again in EMT tumor cells. Furthermore, myCAFs also acted as sources of TGF-β at the advanced tumor-stromal interface, where TAMs 1 emerged as the main TGF-β recipients (**SFig. 10C**). To assess intracellular responses to TGF-β signaling^51^, we quantified TGF-β pathway activity in single-nuclei populations (Methods). Gene set expression increased from fibroblasts to iCAFs and myCAFs, from tumor macrophages to TAMs and from healthy myoepithelial cells to TABACs (**Fig. 6D**). Therefore, TGF-β signaling was upregulated in all interface populations as they transitioned from healthy to the tumor-specific phenotypes, supporting the central role of TGF-β in the remodeling of cellular phenotypes at the tumor-stromal interface.

To determine whether the predicted TGFβ signaling contributes to stromal remodeling, we orthotopically co-transplanted tumor organoids with healthy mammary fibroblasts into immunocompetent mice and treated animals with the TGFβ receptor inhibitor Galunisertib (**Fig. 6E**, Methods). Treatment did not affect endpoint tumor weight (**SFig. 10D**), allowing evaluation of stromal remodeling independently of differences in tumor burden. TGFβ receptor inhibition markedly reduced accumulation of Ncam1+ myCAFs surrounding transplanted tumor cells while preserving the surrounding Chl1+ fibroblast population (**Fig. 6F**). These observations support the computational prediction that early TGFβ signaling promotes progression toward the myCAF-rich stromal architecture reconstructed from the spatial trajectory.

### EMT-like tumor cells invade in a myCAF-rich, remodeled ECM tumor-stromal interface

Beyond secreted signals, cell-ECM interactions play a central role in TME remodeling^52^. We thus extended our spatial communication analysis to ECM ligand-receptor pairs^49^ (**Fig. 6G**).

This revealed dynamic changes in 50 ECM interactions at the tumor-stromal interface (**SFig. 10E**). In particular, Tenascin-X (Tnxb) activity, which is linked with tumor restraining properties^53^, peaked upon tumor initiation (Stage I) and declined sharply with tumor progression (Stages III-IV), while Tenascin-C (Tnc) - its tumor-specific counterpart^54,55^ - linearly increased together with Col1a1 and Lamc2 signaling, peaking at the interface of advanced tumors (Stage IV) (**Fig. 6H**). While Col1a1 was expressed by all fibroblast states - although at highest levels in myCAFs -, we observed a shift in tenascin expression from Tnxb in fibroblasts and iCAFs to Tnc in myCAFs. EMT-like tumor cells also featured Tnc expression and emerged as the only population expressing Lamc2, a recently reported marker of invasive tumor cells^56^ (**Fig. 6I**). To assess cellular contributions to ECM remodeling, we quantified ‘ECM organization’ pathway activity in snRNAseq populations. Signature scores increased at the transition from fibroblasts to iCAFs and peaked in myCAFs, supporting their central role in ECM remodeling (**Fig. 6J**).

Among tumor phenotypes, EMT-like tumor cells displayed the highest pathway scores, compatible with their active role in shaping the ECM in advanced tumors. In contrast, macrophages, including TAMs, contributed minimally to this process. In the TME, ECM organization was spatially restricted to the tumor-stromal interface (**SFig. 10F**). Consistent with these transcriptomic analyses, second harmonic generation (SHG) imaging, a label-free method to visualize fibrillar collagen in tissue sections^57^,revealed collagen-rich regions around transformed ducts and within the dysplastic stroma in a T3 gland (**SFig. 10G-H**). IF staining of the consecutive section then confirmed the colocalization of Ncam1+ and TNC+ myCAFs in these collagen-rich regions(**SFig. 10I-K**).

Interestingly with IF we validated the presence of infiltrating Lamc2+/YFP+ tumor cells within a dense network of Ncam1⁺ stromal cells (**Fig. 6K**, **SFig. 11A**), confirming the tight spatial association of myCAFs with invasive, EMT-like tumor cells. Furthermore, we found Tnc deposition around YFP positive cells at the tumor-stromal interface in primary tumors, especially around those invading surrounding stroma (**SFig. 11 B-C**). Notably, we observed the same remodeling pattern in PDX tumors. ECM signaling has been highest in regions of the tumor occupied by myCAFs (**SFig. 11D**). We observed disseminating, tumor: -Lamc2 (**Fig. 6L, SFig. 11E,F**) and -Ecad (**Fig. 6L, SFig. 11G**) positive cells surrounded by TNC-rich stroma were found as consistent phenotype across genotypes.

Overall, while in the tumor initiation TGF-β emerged as a main transforming factor of TME, myCAFs emerged as the principal drivers of ECM remodeling at the tumor–stroma interface, depositing a collagen- and Tnc-rich matrix that encapsulates invasive tumor cells (**Fig. 6M**).

### Myofibroblasts are sufficient to promote tumor growth and invasion

To determine whether Ncam1+ myCAFs are sufficient to promote tumor progression, we orthotopically transplanted tumor cells^58^ into the mammary fat pad of immunocompetent, syngeneic recipient mice either alone (Group A), together with Ncam1⁻ fibroblasts from healthy controls (Group B), or tumor-derived Ncam1⁺ myCAFs (Group C) (**Fig. 7A**, Methods). Co- transplantation of Ncam1+ myCAFs markedly accelerated tumor growth compared with tumors transplanted alone or with healthy fibroblasts (**Fig. 7B-C**). Histological analysis revealed that tumors from Groups A and B resembled early-stage disease, whereas Group C tumors displayed a disorganized architecture with nests of epithelial cells invading the surrounding stroma (**Fig. 7D**). Co-transplantation of tumor cells with iCAFs (Cd34⁺)^101^ also promoted tumor growth, although less efficiently than Ncam1⁺ myCAFs, indicating that both CAF populations support tumor progression but differ in their ability to establish the late invasive microenvironment (**SFig. 12A-C**).

To investigate how myCAFs reshape TME molecular architecture, we profiled representative tumors from each group using OpenST (**Fig. 7E**). While iCAFs were abundant in the peritumoral stroma in Group A and, to a lesser extent, Group B tumors, they were rare in Group C (**Fig. 7F**). Instead, myCAFs featured the opposite pattern: they were rare in Group A, limited to a peritumoral ring in Group B, while they were abundant and intermixed with tumor cells in Group C tumors. Of note, B cells were abundant in the myCAF-poor periductal stroma of Group A tumors but were absent in Group B and C (**SFig. 12D, SFig. 13A**), suggesting immunosuppressive effects of myCAFs on the TME of the injected mice. Similar to their localization in primary tumors, macrophages were abundant in the peritumoral stroma of Groups A and B (**SFig. 12E, SFig. 13B**), while TAMs 1 occupied the tumor-stromal interface and TAMs 2 formed hotspots in the tumor core across all groups, being particularly abundant in Group C (**SFig. 12F, SFig. 13B**). This stromal remodeling was accompanied by a shift in epithelial cell states, with increased EMT-like tumor populations and reduced keratinizing tumor cells following myCAF co-transplantation (**Fig. 7G-H**). Tumor cells featured late pseudotime in all groups (**SFig. 12G**), however Groups A and B featured abundant Keratinizing and Wnt⁺ phenotypes (**SFig. 12H**), while tumor cells in Group C upregulated EMT signatures (**Fig. 7G-H**). ECM organization pathway activity also increased in Group C and colocalized with myCAFs in space (**SFig. 12I, SFig. 13C**). While a thick collagen layer surrounded tumors in groups A-B, SHG revealed dense and disorganized collagen deposition inside Group C tumor (**SFig. 12J**). Finally, IF staining identified abundant Lamc2+/YFP+ tumor cells invading a Tnc-rich stroma in Group C tumors (**Fig. 7 I-J**, **SFig. 12 K-L**).

Together, these findings demonstrate that Ncam1+ myCAFs are sufficient to remodel the tumor microenvironment and promote the emergence of invasive tumor phenotypes.

## Discussion

Knowledge of the initial steps of TME remodeling is essential for early disease interception, however clinical samples typically provide isolated and heterogeneous snapshots of the tumorigenesis process. Albeit distinct from the stochastic initiation and latency of sporadic tumors, the simultaneous activation of three oncogenes in our model accelerated tumor development along a genetically-defined trajectory and enabled the study of early TME remodeling.

Following oncogene activation in the luminal compartment, mammary epithelial cells consistently acquired basal-like molecular features. This aligns with previous studies from us and colleagues demonstrating that basal-like TNBC tumors can arise from luminal progenitors ^42,43,59–61^. In particular, our experimental design sampling multiple asynchronous lesions across early timepoints enriched for transdifferentiating cells rather than tumor endpoints, which are typically identified in human TNBC. This enabled us to use L2B transdifferentiation as a robust molecular clock to order 100+ tumorigenesis snapshots into 4 progressive tumor stages. While lineage tracing would shed light into the origin of individual tumor populations and potential evolutionary branching events, histological review orthogonally validated tumor molecular staging and supported the use of our framework to investigate how the TME is dynamically remodeled in response to changes in intraductal tumor phenotypes.

This revealed that resident populations surrounding healthy mammary ducts underwent rapid remodeling following tumor initiation: myoepithelial cells were reprogrammed to TABACs and periductal fibroblasts were replaced by iCAFs (early) and myCAFs (late). Although iCAFs preceded the appearance of myCAFs at the tumor-stromal interface and TGF-β signaling has already been shown to drive iCAF-to-myCAF transitions^62–64^, our spatiotemporal framework cannot definitively establish lineage relationships. Hence, alternative cellular origins for myCAFs remain possible (i.e. pericytes as described previously^65^).

Mapping cell-cell interactions dynamics identified which signaling axes are spatiotemporally aligned, hence likely drive, these events. Among the hundreds of ligand–receptor interactions analyzed, transforming growth factor β (TGFβ), a multifunctional cytokine that can stimulate diverse cellular responses, particularly critical to embryonic development, wound healing, tissue homeostasis, and immune homeostasis in health and diesase ^66–69^, emerged as (1) the earliest signaling program associated with TME remodeling, (2) TGF-β 2 was upregulated specifically by cell-of-origin tumor cells, (3) TGF-β receptors were broadly expressed by celltypes at the duct interface including resident fibroblasts, myoepithelial cells and recruited macrophages, (4) intracellular targets of TGF-β signaling were upregulated in all remodeled populations at the tumor-stromal interface. Pharmacological inhibition of TGFβ signaling using Galunisertib perturbed subsequent stromal remodeling, resulting in a iCAF-rich, myCAF- depleted (i.e. early) stroma. In addition to tumor-derived Tgfb2, our analysis identified a progressive increase in Tgfb1 signaling by recruited macrophages, TAMs and myCAFs at the advanced tumor-stromal interface. As Galunisertib blocks signaling from all TGFb isoforms^70^, Tgfb2 knockdown in tumor organoids before injection represents a compelling avenue to isolate the individual contributions of tumor-derived TGFb2 from stroma-derived TGFb1.

In advanced tumors, basal-like tumor cells adopted divergent transcriptomic phenotypes despite sharing the same oncogenic drivers. While somatic mutations may be acquired by tumor cells during progression, several lines of evidence suggest that their contribution to tumor heterogeneity in our model is limited: (1) the short 28-day timeframe likely precludes genetic evolution, (2) prior mouse model studies have shown that phenotypic diversity can emerge from genetically-defined oncogenic inputs alone ^21,28,43^, (3) tumor states were reproducibly organized in specific multicellular niches. In particular, keratinizing tumor cells were confined to the ductal core in close proximity to Wnt-high populations, while EMT-like tumor cells strictly colocalized with myCAFs at the invasive tumor–stroma interface. Although tumor phenotypic endpoints are limited by the genetic drivers (e.g. we and colleagues have previously linked overactive Wnt signaling with squamous differentiation ^43,60,71,72^), our data supports an active role of local microenvironments in shaping tumor cell phenotypes. Importantly, myCAFs, when co-transplanted with tumor cells into syngeneic, immunocompetent mice, were sufficient to shift tumor phenotypes away from keratinization towards EMT phenotypes. While recent work by Aleckovic and colleagues^73^ demonstrated that short-term treatment with *Galunisertib* in tumor-free pubertal mice prevented estrogen- and carcinogen-induced breast cancer *in vivo* targeting secretory basal cells, together our myCAF transplantation and TGFb inhibition experiments support a role for Galunisertib in tumor interception through delaying stromal remodeling and reducing the emergence of tumor- promoting myCAFs.

In line with previous studies linking TGFb signalling ^74,75^, myCAFs^63, 101,102^ and ECM remodeling^76^ with the induction of tumor EMT, in our data myCAFs are the main ECM organizers at the duct stromal interface, contributing to collagen and TNC deposition. These ECM components, together with matrix biomechanics, have been implicated in promoting tumor cell EMT and invasion through outside-in signaling ^76–80^ and mechanotransduction^81^. In our model, EMT-like tumor cells expressed Lamc2, a laminin subunit linked to transformed ECM interactions with receptors such as Sdc1 and Integrins in multiple tumor types^82–84^. Future studies combining pathway and biomechanical perturbations will be required to mechanistically investigate the relative contributions of ECM composition, matrix stiffness, and other myCAF-derived factors to the emergence of invasive tumor phenotypes.

In addition, tumor-infiltrating lymphocytes were rarely detected in our model and transplant experiments in line with the recently reported immunosuppressive role of Ncam1+ myCAFs ^85^. In fact, tumor-only grafts, where myCAFs were scarce, represented the only exception, featuring abundant B cell infiltration in the peritumoral stroma. Together, these findings suggest that targeting CAFs and the signaling pathways governing their emergence could represent an attractive therapeutic strategy to simultaneously restrain tumor invasion and modulate the immune microenvironment.

While the specific *PIKCA^H1047R^ / TP53 ^R175H (R172H^ ^in^ ^mouse)^ / CTNNB ^Δex3^* genotype is uncommon in TNBC patients, the coordinated activation of these pathways recapitulates pathway dysregulation recurrently observed across human TNBC subtypes. In particular, while CTNNB1 mutations are only observed in less than 3% of TNBC samples ^31,86,87^, Wnt/b-Catenin pathway activity is increased in 15-30% of cases with reports up to 80%, especially in claudin- low and metaplastic subtypes ^88,89^. In line with previous studies from us and colleagues of Wnt- driven ‘supra-basal’ tumors ^29,43,60,90–92^ our model displayed characteristic features of squamous differentiation and keratinization. Such aggressive histological features and evidence of the early stromal infiltration in our model are characteristic of metaplastic breast cancers, one of the most aggressive and therapeutically intractable forms of TNBC^93^.

Further supporting their translational relevance, the stromal programs identified here were conserved across genetically diverse PDX models and treatment-naïve human TNBC. Cross-species comparison of mouse and human single-cell datasets identified conserved iCAF and myCAF transcriptional programs, while spatial transcriptomic analyses demonstrated preferential enrichment of iCAFs in carcinoma *in situ* and myCAFs in invasive tumor regions, recapitulating the stromal transition reconstructed from the temporally ordered mouse trajectory. The emergence and spatial organization of CAFs, TAMs and ECM components across genetically diverse PDX models further suggests that this stromal remodeling program is largely independent of tumor genetic drivers.

Beyond fibroblasts, myoepithelial cells and macrophages also emerged as key components of the tumor–stroma interface. The functional role of myoepithelial cells in cancer remains controversial, with evidence supporting both tumor-suppressive and pro-tumor functions ^94,95^. Our spatiotemporal analyses identify TABACs as a distinct myoepithelial state that emerges early during tumor progression possibly from Shh-induced myoepithelial remodeling and persists at the tumor–stroma interface, suggesting that myoepithelial remodeling represents an integral component of early microenvironmental reorganization. Likewise, our analyses distinguish two spatially and functionally distinct tumor-associated macrophage populations, highlighting coordinated remodeling of both fibroblast and immune compartments during tumor progression.

The identification of surface markers specific for remodeled stromal populations, including Ncam1, Cd34, p73, Mrc1, Tgfbr1 and Gpnmb, provides a framework for future lineage-tracing and functional perturbation studies aimed at resolving the cellular origins and functions of individual stromal states. Moreover, because syngeneic transplantation in our system reduces tumorigenesis to a single, experimentally tractable lesion, it provides an ideal platform for CRISPR-based perturbation screens targeting the signaling axes and stromal cell states identified by our spatiotemporal atlas. Such approaches should enable systematic dissection of the molecular events driving early tumor–microenvironment remodeling and help identify candidate targets for intercepting tumor progression before establishment of the mature invasive niche.

In summary, we demonstrate that temporal ordering of spatially resolved disease snapshots allows identifying some of the **molecular ‘forces’** that propel cells/microenvironments along an early disease trajectory. By integrating high-resolution spatial transcriptomics, single-nucleus RNA sequencing, computational modelling, functional perturbation and validation in treatment-naïve human TNBC, we identify a conserved sequence of stromal remodeling in which early TGFβ-associated signaling precedes the establishment of a myCAF-rich invasive niche. As these early signaling axes were rapidly switched off upon tumor progression, profiling of advanced tumors may fail to identify the initial triggers of TME remodeling, which nevertheless represent attractive therapeutic targets. More broadly, we anticipate that combining spatial genomics with temporal reconstruction will provide a general framework for identifying early multicellular events underlying disease progression and for developing strategies aimed at disease interception. Moreover, the advent of computational methods that systematically collect vast amount of published data from health and disease trajectories (including human data) in a unified framework ^96^ may enable to identify molecular space-time hallmarks of tumorigenesis trajectories across different cancers.

## Author contribution

K.L. generated and characterized the mouse model, derived and characterized *in vitro* lines derived from the model, supervised the injection experiment. I.T. performed snRNA-seq, spatial transcriptomics experiments, initial processing and computational analyses of the sequencing data. T.M.P. performed computational analyses. S.B. and F.H provided support for immunofluorescence experiments and organoid characterization. D.L.P., A.A. and N.K. provided support for computational analyses. M.M and A.X. provided support for the mouse model characterization, IFs, mouse colony handling and genotyping. H.R., B.S. and S.F. performed histology characterization. S.R. and A.M. performed imaging of IF and second harmonic generation imaging experiments. J.L., M.S. and A.B. provided technical support for snRNA-seq and spatial-transcriptomics experiments. T.C. assisted with mouse model characterization and helped with manuscript revising. D.I.R., helped with manuscript revising S.D. performed spatial transcriptomics analyses under the supervision of M.N., S.K. performed and interpreted electron microscopy experiments. Eli.M., Elo. M., generated the xenograft sample cohort. W.B. provided mouse model supervision. K.L., W.B. and N.R. provided funding acquisition. K.L., I.T., T.M.P., and N.R. conceptualized and designed the project. N.R. supervised the project. K.L., I.T., T.M.P. and N.R. wrote the manuscript with input from all authors.

## Acknowledgements

K.L. was financially supported by the Deutsche Krebshilfe (DKH), grant number: 70113738 and Immunoprecept cluster of Excellence. I.T. was the recipient of an EMBO long term post-doctoral fellowship. T.M.P. is financially supported by the Berlin School of Integrative Oncology through the GSSP program of the German Academy of Exchange Service (DAAD) and by the Add-on Fellowship of the Joachim Herz Foundation. N.R. thanks the Deutsche Forschungsgemeinschaft (DFG) grant number RA 838/5-1, Deutsches Zentrum für Herz-Kreislauf-Forschung (DZHK) grant numbers 81Z0100105 and 81X2100155, the Chan Zuckerberg Foundation (CZI)/Seed Network, and the NeuroCure and Immunoprecept Clusters of Excellence in the neurosciences at the Charité – Universitätsmedizin Berlin. We thank Jeannine Wilder, Tatiana Borodina and the genomics facility of MDC for sequencing the samples. We thank Caroline Braeuning and the BIMSB FACS facility for cell sorting. We thank Diana Behrens and the EPO Gmbh team for coordinating and performing the injection experiment. We thank Giuseppe Macino, Elena Splendiani and Jan Licha for helping with Open-ST protocol optimization during their stay in the Rajewsky lab. We thank Takako Sasaki for sharing the laminin gamma (Lamc2) antibody. We thank Elmir Mahammadov for computational support. The authors thank the whole Rajewsky lab for discussions and constructive feedback on the manuscript.

## Resource availability

### Lead contact

Further information and requests for resources and reagents may be directed to Nikolaus Rajewsky.

### Data and code availability

Open-ST RNA-seq and microscopy raw and processed data have been deposited at GEO and will be publicly available as of the date of peer-reviewed publication.

All original code will be publicly available as of the date of publication.

Any additional information required to reanalyze the data reported in this paper is available from the lead contact upon request.

## Methods

### Mouse Strains

All animal experiments were conducted in accordance with European, national, and MDC regulations. WAP-iCre (B6129-Tg(Wap-cre)11738Mam/J Stock No: #:003552), Ctnnb1 ex3 (Ctnnb1tm1Mmt), YFP (B6.129X1-Gt(ROSA) 26Sortm1(EYFP)Cos/J, Stock No: 006148) mice were crossed into FVB/N background and previously described^43^. 129S-Trp53^tm2Tyj/J (Stock No: 008652) and FVB.129S6-Gt(ROSA)26Sor^tm1 (Pik3ca*H1047R)Egan/J (Stock No: 016977) mice were purchased from Jackson Laboratories and crossed into FVBN mouse background for more than 10 generations. Animal experiments were approved by the ethical board Landesamt für Gesundheit und Soziales (LaGeSo), Berlin (Approval G#0213.18). Tumor dynamics analysis is described in the result section of the manuscript, in short: tumor mutant mice developed palpable tumors from the 8th week onward. This occurred without additional stimulation of the promoter with pregnancy, induction of the promoter occurred around the 5th week of life of mice. Tumors were developed selectively in all 10 mammary glands of each mouse carrying the oncogenic genotype (GOF mutation in Trp53, Ctnnb1, Pik3ca and YFP with promoter WAPiCre), no other organs were affected with tumor formation, no other phenotype apart from mammary gland tumors growth was observed. The tumors were harvested between 6 and 11 weeks of mouse age and reached a maximum size of 1 cm³. For genotyping, ear biopsies were digested in lysis buffer (100 mM Tris, pH 8.0–8.5, 10 mM EDTA, pH 8.0, 0.2% SDS, and 200 mM NaCl) with 1:30 10 mg/ml Proteinase K (Roche, Cat. #03 115 879 001). Lysates were then diluted 1:20 in nuclease-free water, and PCR was carried out using Taq DNA polymerase (Invitrogen). The following primers were used for genotyping:

### Kaplan Meier

Mouse survival was analyzed using the Kaplan–Meier method in GraphPad Prism (version 11; GraphPad Software, San Diego, CA, USA). Survival time was calculated from birth until euthanasia at the predefined humane endpoint, which was reached when a palpable mammary tumor exceeded 1 cm in diameter, in accordance with the approved animal welfare protocol and score sheet. Survival curves were compared using the log-rank (Mantel–Cox) test. A two-sided *P* value < 0.05 was considered statistically significant

### List of Genotyping primers

#### Target/Primer Name Sequence

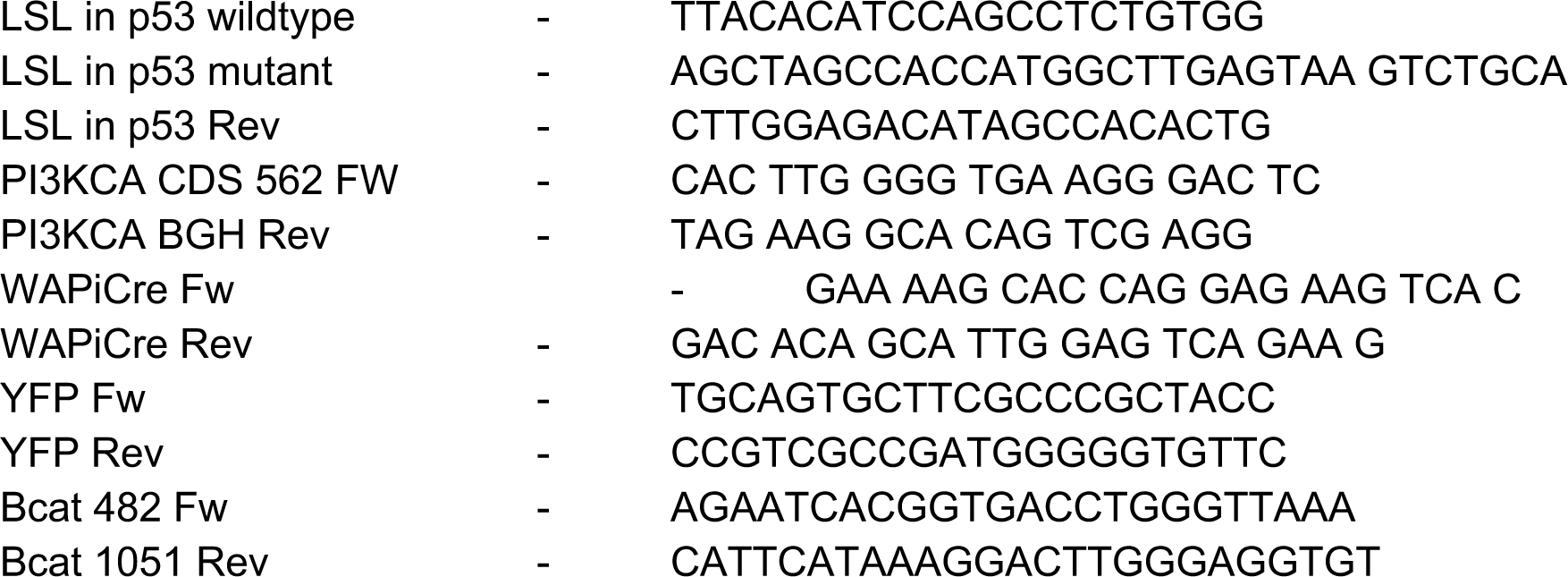

### Histology and Immunostaining

Mammary glands were fixed overnight at 4°C in 4% formaldehyde (Roth), then dehydrated and embedded in paraffin. For staining, tissue sections were deparaffinized, rehydrated, stained with Hematoxylin and Eosin (H&E), dehydrated, and mounted. Images were acquired with a bright-field Zeiss microscope. For immunostaining, antigen retrieval was performed on paraffin-embedded sections by boiling in Tris-EDTA buffer (10 mmol/L Tris, 1 mmol/L EDTA, pH 9.0) for 20 minutes. Cryosections were directly frozen in OCT after tissue was excised from the mice. Before the staining sections were fixed with 4% methanol-free paraformaldehyde for 15 minutes.

All samples were blocked with 10% horse serum for 1 hour at room temperature before overnight incubation with primary antibodies at 4°C (see STable5 for antibody list). For fluorescent staining, samples were incubated with secondary antibodies and DAPI for 1 hour at room temperature before mounting with Immu-Mount (Thermo Scientific).

For immunohistochemistry, after antigen retrieval, slides were washed, incubated with 3% H₂O₂ for 15 minutes at room temperature, washed three times for 5 minutes in PBS, and incubated with primary antibodies overnight at 4°C. Horseradish peroxidase-labeled secondary antibodies were diluted and applied on slides for 1 hour at room temperature. Slides were then washed, developed using DAB chromogenic substrate (DAKO), dehydrated, and mounted.

### Histology and Immunohistochemistry (IHC)

Brightfield imaging of hematoxylin and eosin (H&E) stained paraffin sections and chromogenic immunohistochemistry (IHC) was performed using the 3DHISTECH Pannoramic SCAN II slide scanner (20× Plan-Apochromat objective, NA 0.8). Full mammary gland sections were scanned at 20× magnification under brightfield illumination. Stitching and image rendering were performed using CaseViewer software. For IHC, diaminobenzidine (DAB) chromogen was used, and scanned slides were evaluated uniformly with matched exposure settings. All scans included full-gland imaging for downstream analysis of lesion heterogeneity and tumor progression stages.

### Immunofluorescence imaging

Fluorescently stained cryosections were imaged using both widefield and confocal platforms. Widefield fluorescence imaging was conducted on a Leica Thunder Imager 3D Tissue system using a DFC9000 GTC sCMOS camera and a 20× HC PL APO dry objective (NA 0.75). Images were acquired at 20× magnification and stitched using Leica LAS X software when required for whole-gland visualization.

High-resolution confocal imaging was carried out using a Leica Stellaris 5 system equipped with a White Light Laser and HyD detectors. Sections were scanned using a 20× HC PL APO CS2 glycerol objective (NA 0.75) with 1024×1024 pixel resolution and sequential laser scanning. Z-stacks were collected when necessary for visualization of spatial organization within tumor-stromal niches. All image acquisitions were performed using standardized laser settings and exposure times to allow quantitative comparisons across conditions.

### Isolation of mammary gland cells

Tumors were minced and digested in DMEM/F12 HAM (Invitrogen) supplemented with 5% FBS (Invitrogen), 5 mg/mL insulin (Sigma-Aldrich), 0.5 mg/mL hydrocortisone (Sigma-Aldrich), and 10 ng/mL EGF (Sigma-Aldrich), forming a digestion medium containing 300 U/mL collagenase type III (Worthington), 100 U/mL hyaluronidase (Worthington), and 20 mg/mL Liberase (Sigma). Digestion was carried out at 37°C for 1.5 hours with shaking.

The resulting organoids were resuspended in 0.25% trypsin-EDTA (Invitrogen) at 37°C for 1 minute, then dissociated in digestion medium containing 2 mg/mL Dispase (Invitrogen) and 0.1 mg/mL DNase I (Worthington) at 37°C for 45 minutes while shaking. Samples were filtered using 40 μm cell strainers (BD Biosciences) and incubated with 0.8% NH₄Cl solution on ice for 3 minutes for red blood cell lysis. Lysis was stopped by washing in 30 mL Dulbecco’s Phosphate-Buffered Saline (DPBS) without Ca²⁺ and Mg²⁺ (Gibco, catalog no. 14190169). The resulting pellets were used for downstream applications as outlined below.

### Organotypic Stem Cell-Enriched 3D Cultures

Organotypic stem cell-enriched cultures were generated using previously published protocols, with minor modifications. Single cells from digested mammary glands were resuspended and plated on collagen I-coated plates (50μg/mL) in stem cell-enriching medium MEBM (Lonza Cat. #CC-3151), supplemented with 2% B27 (Invitrogen, Cat. #17504044), 20 ng/mL bFGF (Invitrogen, Cat. #13256029), 20 ng/mL EGF (Sigma, Cat. #SRP3196-500 μg), 4μg/mL heparin (Sigma, Cat. #H3149), 5μg/mL insulin (Sigma, Cat. #I0516-5ml), 0.5μg/mL hydrocortisone (Sigma, Cat. #H0888-1G), and 1X Gentamicin (Sigma, Cat. #G1397-100ML) for 16–18 hours.

Cells were washed twice with DPBS, treated with 0.25% trypsin-EDTA for 30 seconds, and then incubated with TrypLE (Thermo Fisher#12604021) at RT for 5–7 minutes until detachment. Cells were resuspended in stem cell-enriching medium, counted, and seeded in 25μL droplets containing 95% reduced growth factor Matrigel at a density of 100 cells/μL. Plates were carefully inverted, and Matrigel was allowed to solidify at 37°C for 45 minutes to 1 hour.

Then, 0.5 mL of stem cell medium was added per well of 24-well plates containing a 25 μL droplet. Medium was changed every second day. For secondary sphere formation, spheres were dissociated for 10 minutes in TrypLE, and cells were filtered using 0.45μm filters. Single-cell suspensions were re-seeded as described for primary cells. All treatments with verteporfin were performed under minimal lighting. Tumor cell were enriched and isolated by sorting them for the presence of YFP using Fluorescence -activated cell sorting (FACS)

Fibroblast cultures were isolated the same way as epithelial compartment but were grown in DMEM, FBS 10%, P/S 1% on uncoated plastic dishes. After plating cells were detached, FACS sorted for the absence of CD45 to exclude all of possible contamination with immune cells and YFP to exclude the chance of tumor cells growth.

### Fluorescence -activated cell sorting (FACS)

Single-cell suspensions from dissociated tumors were resuspended at 10,000 cells/μl and incubated with conjugated primary antibodies at 4°C for 15 minutes. Cells were then washed three times in DPBS and subsequently incubated with 7-AAD (5 μl per 10⁶ cells) at room temperature for 5 minutes to stain dead/dying cells. Cells were sorted using the FACSAria II or III (BD Biosciences) or analyzed using the LSRFortessa (BD Biosciences). Compensation and unstained controls were carried out before FACS experiment. Data is shown for three independent biological experiments from cell lines derived from three different animals, that were analyzed using FlowJo Analysis Software.

### Patient-derived xenograft models

Patient-derived xenograft (PDX) models were established from patients with triple-negative breast cancer with their informed consent as described previously (Coussy et al., 2019). The experimental protocol and animal housing were in accordance with institutional guidelines as proposed by the French Ethics Committee (agreement no. B75-05-18). PDXs were obtained from patients with their informed written consent.

### Cell grafting

Engraftment experiments were conducted at EPO GmbH, Berlin-Buch, and performed according to the German Animal Protection Law with approval from the responsible authorities (LAGeSo, approval number E0023-23). The in vivo procedures were consistent and in compliance with the UKCCCR guidelines.

Ncam1 was used as a marker of Cancer Associated Fibroblasts (myCAFs) to sort these cells by FACSs (FAB7820P, R&D Systems). Ncam1**^high^** population of CAFs were derived from transformed glands of T2 from tumor lines (P3WeY). Ncam1**^low^** Normal Fibroblasts (NFs) were derived from healthy ctrl mammary glands (WY genotype). Cells were screened for the absence of Immune cell marker Ptprc/Cd45 (25-0451-81, Invitrogen) and lack of YFP expression (it would indicate the contamination of fibroblasts with tumor cells).

Accordingly tumor cells were sorted for the presence of YFP as explained method section: mammary gland cells isolation.

2 biological replicates for each of the lines, organoids, Normal fibroblasts and CAFs were used for the injection experiments. First tumor organoids were dissociated to single cells in Tryple for 30min in 37°C and filtered through Flow me 40um mesh (FlowMi)Cell strainer, then counted in Trypan blue. Accordingly, Fibroblasts cultures were detached by Trypsin and counted in Trypan blue.

2 x 10^6^ tumor cells were injected into fat pads into immunocompetent, 7 week old, healthy mice either: alone (Group A), with 2 x 10^6^ of normal fibroblasts (NF) (Group B) or with 2 x 10^6^ of CAFs (Group C). Every injection was supplemented by equal volume of BME (Cultrex UltiMatrix RGF BME, R&D systems). Every injection consisted of cells resuspended in 20 ul of Medium that was additionally supplemented with 20ul of BME. Addition of BME aimed to help the cells to recover after dissociation procedure.

Each of 5 mice which we used per group was injected into two mammary fat pads making 2 experimental points per mouse. After injection mice were routinely (every 2 days) monitored for palpable tumor appearance and weighted. After the largest tumors reached the size of 1,5cm all mice were sacrificed at the same times. Glands were collected either for paraffin or direct freezing in OCT. iCAF cells were isolated from tumors in the same strategy as myCAFs, however Cd36 was used as a marker of iCAF to sort these cells by FACSs (17-0362-82, Themofisher). Cd36**^high^**

/Ncam1- population of CAFs were derived from transformed glands of T2 from tumor lines (P3WeY). CD36 was also reported previously as iCAF marker (Houthuijzen et al., 2023) Cells were then grown on soft collagen (9ug/ul) surface to retain the iCAF phenotype according to report from Crozier et al 2023. Cells were screened for the absence of Immune cell marker Ptprc/Cd45 (25-0451-81, Invitrogen) and lack of YFP expression.

### Galunisertib treatment

Normal Fibroblasts (NFs) that were derived from healthy control mammary glands (WY genotype) were injected as described in the Cell grafting section, together with the same YFP+ tumor cells (Mammary gland cells isolation). Tumor organoids were co-transplanted orthotopically with normal mammary fibroblasts into recipient mice. Following transplantation, mice were treated once a day by oral gavage with either vehicle control or the ALK5/TGFβ receptor I (TGFβRI) inhibitor galunisertib at a dose of 37.5 mg/kg. Treatment was continued for 26 days after transplantation, at which point mice were euthanized before substantial tumor expansion to enable the assessment of stromal remodeling independently of differences in tumor burden. Tumors were harvested, processed for histological and other downstream analyses.

### Electron microscopy

Mammary gland tissue, healthy and tumor as well as organoids were processed for ultrastructural analysis. All sample types were fixed by immersion in 2% (w/v) formaldehyde and 2.5% (v/v) glutaraldehyde in 0.1 M phosphate buffer for 2 hrs at room temperature followed by an overnight incubation at 4°C. Samples were post-fixed with 1% (v/ v) osmium tetroxide (Sigma-Aldrich), dehydrated in a graded series of EtOH and embedded in PolyBed 812 resin (Polysciences). Ultrathin sections (60–80 nm) were poststained with 0.5% (w/v) uranyl acetate (Sigma-Aldrich) and 3% lead citrate (Leica microsystems). Sections were imaged using back-scattered electrons at 2 kV, 0.8nA with scanning electron microscope Helios 5CX (Thermo Fisher Scientific). Acquisition was performed with the retractable DBS detector in immersion mode and the MAPS software (Thermo Fisher Scientific).

### snRNAseq data generation

Healthy or tumor pieces from the abdominal mammary gland were snap-frozen in liquid nitrogen. For processing, half a gland was used for healthy and early tumors and only a piece was used for middle and late tumors. Frozen tumor pieces were kept on ice and chopped into pieces with a cold scalpel in 500 µl of lysis buffer (10 mM Tris-HCl PH 7.4, 10 mM NaCl, 3 mM MgCl2, 0.1% NP-40, 1 mM DTT, 1u/µl Rnase Inhibitor). The tissue was further collected and processed on ice in 1.5 mL tubes for the rest of the procedure. First, 800 µl of lysis buffer was added, the tissue was crushed with a plastic pestle 10 to 15 times and incubated for 5 minutes with gentle resuspension from time to time using a 1000 µl pipette. Tissue pieces were then collected at the bottom of the tube with a fast spin down, the supernatant was strained in a new tube using a 100 µl filter (pluriselect) and spun down at 500G in a swing bucket rotor for 5 min at 4°C. The supernatant was removed and 1 ml of wash buffer (1% BSA, 1u/µl Rnase Inhibitor in BSA) was gently added without disturbing the nuclei pellet and incubated for 5 minutes. Nuclei were spun down at 500 G in a swing bucket rotor for 5 min at 4°C. An additional wash was performed in 1 ml of wash buffer and after a final centrifugation, nuclei were resuspended in 500 µl of wash buffer and filtered through a 40 µm strainer (pluriselect). DAPI was then added to a final concentration of 2 µM. In the case of T2 and T3 tumor samples that presented a higher concentration of debris, around 250,000 nuclei were sorted based on DAPI using a BD FACS Aria III with a 100 µm nozzle in 250 µl of sorting buffer (4% BSA, 4u/µl Rnase Inhibitor in BSA). Nuclei were then counted using a Neubauer counting chamber with a Keyence microscope and processed using the 10X Genomics 3’ V3.1 kit following the manufacturer instruction with a target of 8,000 to 9,000 recovered nuclei. The only adjustment concerned the number of PCR cycles which was increased by one accounting for the usual lower concentration of RNA in nuclei in comparison to cells. Libraries were sequenced with the recommended read lengths. The snRNA-seq dataset consisted of 12 biological samples spanning four experimental groups (healthy control and three tumor progression stages), with three biological replicates per group.

### snRNAseq data processing

Fastq files were generated using Cellranger v7.1.0 (‘Cellranger mkfastq’) and aligned to the mm10 mouse genome (‘Cellranger count’) to generate digital gene expression matrices with default parameters. Filtered gene expression matrices were imported in R v4.4.3 for downstream processing using Seurat v5.1.0^97^. Single samples were merged into a single Seurat object and nuclei with less than 1000 genes and more than 50% of mitochondrial transcripts were removed. Doublets were identified using *scDblFinder* v1.18.0^98^ and nuclei with a doublet score higher than 0.5 were removed.

### snRNAseq data normalization and clustering

Raw UMI counts were normalized using Seurat’s LogNormalize function with a scale factor of 10,000 transcripts per nucleus, followed by natural logarithm transformation and feature-wise scaling prior to dimensionality reduction and clustering. Principal component analysis was performed on the scaled data, after which a shared nearest-neighbor (SNN) graph was constructed and clusters were identified using the Louvain modularity optimization algorithm implemented in Seurat’s FindClusters() function. Clustering resolutions were selected according to the specific analysis performed, including a resolution of 1.5 for global cell-type identification and 0.4 for epithelial state identification and multicellular niche identification.

### snRNAseq cell type identification

The top 3,000 variable genes were used to compute 30 principal components and UMAP coordinates, while a subset of 2964 genes with standardized variance higher than 1 in both snRNAseq and Open-ST data was used to identify clusters (resolution= 1.5, dims=1:30). Cluster annotation were supported by marker gene expression (FindMarkers only.pos=T, min.pct= 0.2, min.diff.pct=0.1, max.cells.per.ident=500) and label transfer scores from a murine healthy and tumor mammary gland atlas^41^. For label transfer, the atlas was subsetted to 1000 cells per identity (‘CellTypesFinal’) before label transfer (FindTransferAnchors normalization.method=‘LogNormalize’, reduction=‘pcaproject’, dims=1:30, non.method=‘rann’, eps=0.5).

### Batch effect evaluation

Samples were sequenced in three sequencing batches and potential batch effects were evaluated using the CellMixS R package v1.20.0. Batch mixing was first assessed qualitatively by visualizing the distribution of sequencing batches in UMAP space (**Method supplementary Fig. S1A**). Quantitative assessment was then performed by calculating the Cell Mixing Score (CMS), which measures the diversity of metadata labels within each nucleus’s local transcriptional neighborhood. CMS was computed on the first 30 principal components using a *k*-nearest neighbor graph (*k* = 200) for four metadata variables: sequencing batch, sample identity, experimental timepoint, and tumor status (healthy versus tumor-bearing glands). Smoothed CMS values (`cms_smooth`) were used for downstream visualization and comparison both across the complete dataset (**Method supplementary Fig. S1B**) and within individual annotated cell types (**Method supplementary Fig. S1C**). Sequencing batch exhibited the highest degree of local mixing (median CMS = 0.31), followed by sample identity (median CMS = 0.27) and experimental timepoint (median CMS = 0.26), whereas tumor status showed substantially lower mixing (median CMS = 0.12). Across all annotated cell types, sequencing batch, sample identity, and experimental timepoint consistently exhibited higher mixing than tumor status, indicating that the transcriptional organization of the dataset was driven predominantly by biological differences associated with tumorigenesis rather than by technical batch effects.

### snRNAseq epithelial state identification

Epithelial cells (including ‘Hormone sensing’, ‘Luminal’, ‘Myoepithelial’ and ‘Tumor’) were integrated (IntegrateLayers method= CCAIntegration) and re-clustered. Doublets identified by means of marker gene expression (‘Ptprc’, ‘Mrc1’ and ‘Meg3’) and higher scDblFinder scores were excluded. UMAP coordinates (reduction= ‘epithelial.cca’, dims=1:30) and clusters were recomputed (resolution=0.4). Cluster annotations were supported by marker gene expression (FindMarkers only.pos=T, min.pct= 0.2, min.diff.pct=0.1, max.cells.per.ident=500), label transfer scores and enrichment for control and tumor-bearing samples. UMAP 2D densities were computed over a regular grid using MASS v7.3-65 (kde2d n=100) for each condition after balancing the number of epithelial cells from both conditions (slice_sample n=200) and then subtracted. Stromal densities were computed in a similar way, modified to account for the higher number of stromal cells (slice_sample n=4000, kde2d n=1000).

### Differential abundance analysis using MILO

Differential abundance analysis was performed using miloR R package v2.0.0^99^ to identify cellular neighborhoods associated with tumor progression. Analyses were performed independently for epithelial and stromal compartments. Seurat objects were converted to SingleCellExperiment objects and MILO neighborhood graphs were constructed using high-dimensional transcriptional representations (epithelial CCA space for epithelial cells and PCA space for stromal cells). k-nearest-neighbor graphs were generated using k = 20 and 30 dimensions, and refined neighborhoods were defined using the makeNhoods() function with a neighborhood proportion of 0.2. Neighborhood distances were calculated using the same reduced-dimensional representations. Cell counts within each neighborhood were aggregated by biological sample to account for biological replication. Differential abundance testing between healthy and tumor-bearing conditions was performed using the testNhoods() function with tumor status as the explanatory variable. The resulting neighborhood log-fold changes were used to identify tumor-enriched and healthy-enriched cellular neighborhoods. Differentially abundant neighborhoods were subsequently annotated according to their predominant epithelial or stromal cell states using annotatehoods() and visualized using neighborhood graph representations and differential abundance summaries. For visualization purposes only, MILO differential abundance results were projected onto UMAP embeddings; statistical testing was performed directly on the high-dimensional neighborhood graph.

### snRNAseq tumor pseudotime ordering

Tumor pseudotime was computed using Palantir v1.3.2.^100^ Healthy (i.e. ‘Hormone sensing’, ‘Luminal’ and ‘Myoepithelial’) and proliferating populations were excluded. 10 components were used as diffusion map input (run_diffusion_maps n_components=10, determine_multiscale_space). Pseudotime was computed starting from a Wap-expressing ‘Cell-of-origin’ cell from a T1 sample (run_palantir start_cell= ‘T1_Rep2-GTTACCCGTGTTCGTA’, num_waypoints=500). The robustness of the inferred trajectory was evaluated by stratified bootstrap resampling of the single-nucleus dataset. 50 independent bootstrap iterations were performed. For each bootstrap iteration, 20% of nuclei were randomly removed and the diffusion map manifold was recalculated using the same parameters, followed by independent Palantir trajectory inference using the same root cell and number of waypoints. Concordance between bootstrap-derived and original pseudotime assignments was quantified using Spearman correlation. Sensitivity to root-cell selection was assessed by repeating Palantir trajectory inference using all 60 Wap-expressing cells from the cell-of-origin population as alternative starting cells. Trajectories were reconstructed using the same diffusion map and Palantir parameters described above. Consistency between root-cell- derived trajectories was evaluated by calculating pairwise Spearman correlations between pseudotime assignments and the average per-cell pseudotime variance across inferred trajectories. Pseudotime remained highly concordant (median pairwise Spearman correlation 0.9789 and 0.9997 and a per-cell pseudotime variance of 0.0008 and 0.0005 for the bootstrap and root cell analyses, respectively), supporting the robustness of the inferred trajectory (**New Method Fig. ure 2**).

### Macrophage state characterization

Macrophage populations were subsetted from the annotated single-nucleus RNA-sequencing dataset based on previously assigned cell-state identities (healthy macrophages, tumor macrophages, TAM1, and TAM2). Differentially expressed genes between macrophage populations were identified using Seurat’s FindAllMarkers() function with the following parameters: max.cells.per.ident = 1000, min.pct = 0.10, and logfc.threshold = 0. The top 20 marker genes per macrophage population were selected based on adjusted P value, average log2 fold-change, and detection frequency difference and visualized using heatmaps. Pathway enrichment analysis was performed using ranked macrophage marker genes with the Gene Set Enrichment Analysis (GSEA) implementation from the clusterProfiler R package v4.12.6. For each macrophage population, marker genes were ranked by average log2 fold-change, and enrichment analysis was performed against Hallmark gene sets from the Molecular Signatures Database (MSigDB) using msigdbr R package v25.1.1. Enriched pathways were visualized based on normalized enrichment scores (NES) and adjusted P values.

### Open-ST data generation

Healthy and tumorigenic mammary tissue were embedded into OCT before being frozen and kept at −80°C. 12 µm sections were cut from blocks in a cryostat and placed on flow-cell pieces prepared according to the Open-ST protocol. Samples were processed according to the Open-ST protocol33. Briefly, tissue sections were fixed for 30 min in pre-chilled methanol at −20°C and stained with hematoxylin and eosin (H&E). Images of the stained sections were acquired at 10X or 20X magnification on a Keyence microscope. The tissue on the flow cell was incubated with permeabilization solution for 30 min at 37°C then washed in RT buffer (SSIV 1X RT buffer, Rnase inhibitor 1U/µl) before being incubated overnight at 42°C in RT (SSIV 1X RT buffer, 5 mM DTT, BSA 0.187 mg/ml, 1 mM dNTP mix, 6.67 U/µl, Ribolock Rnase inhibitor 1U/µl). The next day exonuclease treatment (1X Exo1 buffer, Exo1 1U/µl) was performed for 45 min at 37°C followed by tissue removal (100 mM Tris-Hcl PH 8.0, 200 mM NaCl, 2% SDS, 5 mM EDTA, Proteinase K 16 mU/µl) for 40 min at 37°C. Tissues were washed 3 times with ultrapure water, 3 times with 0.1 M NaoH (with 5 min incubation each time), 3 times with 0.1 M Tris PH 7.5 and 3 times with ultrapure water. Second strand synthesis (1X NEBuffer-2, 10 µM randomer primers, 1 mM dNTPs, Klenow exo (-) fragment enzyme 0.5 U/µl) was then performed for 2 h at 37°C. The flow-cell pieces were washed 3 times with ultrapure water, and the second strand product was eluted in 2 rounds of 100 µl 0.1M NaOH. The final 200 µl of second strand product was mixed with 28.6 µl of 1M Tris-HCl pH7.5. The product was purified using Ampure beads at a 1.8x ratio according to the manufacturer instructions and eluted in 82.5 µl of ultrapure water. 2.5 µl of purified product was used in a qPCR (1x blue S-Green qPCR mix + ROX, 1 µM forward primer, 1 µM reverse primer) reaction to determine the concentration. The threshold of 50% of the peak deltaRN was used to determine the number of cycles used in the subsequent PCR reaction using unique index barcodes for each sample. This number of cycles minus 5 was used in the PCR amplification of the cDNA where a 200 µl reaction was split into 4 50 µl tubes. The 200 µl PCR product was purified using Ampure XP beads at a 1X ratio according to the manufacturer instructions and eluted in 20 µl of ultrapure water. The size and concentration of the cDNA was checked on a BioAnalyzer or a Tapestation. The final size selection of fragments between 350 and 1100 bp was performed using a BluePippin or Pippin HT instrument according to the manufacturer instructions. The final library sizes were assessed using Tapestation or Bio Analyzer and the concentration was measured using Qubit. If a small contamination peak was still observed below 300bp, an additional AmPure bead purification was performed at a ratio of 1X. Open-ST profiling was performed on an independent set of 12 biological samples covering the same four experimental groups, with three biological replicates per group.

### Open-ST data processing

Raw BCL files were demultiplexed using BCL2FASTQ and mapped to the mm10 mouse genome using spacemake v0.7.976 with the default parameters of the Open-ST barcode flavor. For downstream analyses, neighboring spatial barcodes were aggregated into a 7 µm hexagonal mesh (∼120 µm² pseudocells). This binning strategy increased transcript capture robustness while preserving spatial resolution close to individual mammary cells, enabling identification of cellular compartments and cell-state distributions across entire tissue sections we used the 7 µm hexagonal mesh. When not done automatically during the spacemake pipeline, stitching of the images and data matrix were done using the stitch.py function of the Open-ST package. Samples were filtered to remove unwanted tiles, exported as 10X Genomics matrix folders, and imported into R for conversion into Seurat objects. Samples were filtered according to sample-specific threshold accounting for sequencing depth (T0_1:400, T0_2:100, T0_3:100, T1_1:250, T1_2:100, T1_3:100, T2_1:500, T2_2:200, T2_3:200, T3_1:100, T3_2:150, T3_3:200 genes). Raw gene expression counts were normalized and variance stabilized using SCTransform using Seurat.

### Open-ST cell type identification

Unsupervised Open-ST clustering was performed independently of snRNA-seq annotation to evaluate whether major transcriptional compartments identified in the spatial data recapitulated the cell types identified in the reference snRNA-seq dataset. The top 2,000 variable genes were used to compute 10 principal components and identify clusters (resolution=0.8). Cluster annotations were supported by marker gene expression.

### Gene set scoring in single nuclei and spatial data

Gene set scores in individual cells were computed using AUCell v1.26.078. For each cells in either single nuclei or spatial data, genes were ranked by their raw expression (AUCell_buildRankings) and the gene set area under the curve score was computed (AUCell_calcAUC). Scores were then added as a new assay and scaled (ScaleData). Stromal signatures were obtained from Mayer et. al., 2023 while gene sets were selected from the HALLMARK, REACTOME and KEGG databases using msigdbr v7.5.1.

Gene set activity was quantified in individual nuclei and pseudocells using the AUCell R package (v1.28.0). For each nucleus or spatial pseudocells, genes were ranked according to their raw expression values using AUCell_buildRankings(), and gene set enrichment was quantified by calculating the area under the recovery curve (AUC) with AUCell_calcAUC(). The resulting AUC scores were added as a separate assay in Seurat and scaled using ScaleData() for downstream visualization and analysis. Stromal gene signatures were obtained from Mayer et al. (2023), while additional gene sets were retrieved from the Hallmark, Reactome, and KEGG collections of the Molecular Signatures Database (MSigDB) using the msigdbr R package (v7.5.1). To characterize macrophage states, published transcriptional signatures for inflammatory (M1) and alternatively activated (M2) macrophages^45^, as well as TREM2/SPP1 tumor-infiltrating and FOLR2/LYVE1 tissue-resident macrophages^46^, were scored using the same AUCell workflow. A complete list of all gene signatures used in this study is provided in Supplementary Table 7.

### Open-ST Robust Cell Type Deconvolution

Robust Cell Type Deconvolution^40^ scores of snRNAseq cell states were computed using spacexr v2.2.1 to map cell states identified by snRNA-seq onto Open-ST spatial transcriptomic data. The snRNA-seq dataset was used as a reference atlas of annotated cell types and cell states after excluding proliferating populations (Immune proliferating, Fibroblast proliferating, and Tumor proliferating) and epithelial doublets. To avoid overrepresentation of abundant populations, a maximum of 500 nuclei per annotated cell state was retained (n_max_cells = 500). To allow the same reference atlas to be applied consistently across all samples, 12 primary and 3 transplant Open-ST spatial datasets were merged prior to RCTD analysis (run.RCTD) and spatial regions annotated as nerve or muscle were excluded as these populations were not represented in the snRNA-seq reference. Spatial pseudocells were not assigned to a single cell identity; instead, RCTD generates probabilistic estimates of the contribution of each reference cell state to each spatial pseudocell. These scores were used for downstream analyses of spatial organization, multicellular neighborhoods, and epithelial– stromal composition.

### Open-ST niche identification

To identify multicellular niches, we first identified the 30 closest spatial neighbors for each cell using dbscan v1.2.0^101^ on the stitched x,y coordinates (kNN, k=30). Neighbors further than 50 µm were excluded (pixel-to-micron scaling factor=1/30) and the neighborhood matrix was computed by averaging RCTD scores per cellular neighborhood. Dimensionality reduction and clustering were then performed on the scaled neighborhood matrix as described for snRNAseq (ScaleData, RunPCA, RunUMAP dims=1:6, FIindNeighbors dim=1:6, FindClusters resolution=0.4). Niches were annotated analyzing the average RCTD scores per cluster and their spatial localization.

### Open-ST pseudotime scoring

Seurat integration pipeline was used to map tumor pseudotime scores in space. To avoid missing values, tumor pseudotime scores were set to 0 for non-tumor cells in the single cell reference.

To simultaneously score all samples in a group, merged primary and EPO seurat objects were generated from 12 and 3 samples, respectively. Gene expression data was subset to shared 2,964 variable genes, normalized (SCTransform) and, following feature selection (SelectIntegrationFeatures), transfer anchors were identified between single nuclei and merged spatial objects (FIndTransferAnchors, normalization.method = “SCT”, nn.method= ‘rann’, eps = 0.5) to transfer pseudotime scores (TransferData prediction.assay = TRUE, weight.reduction = ‘pcaproject’, dims = 1:30, eps = 0.5).

### Open-ST duct segmentation

Duct segmentation was performed via manual lasso selection using plotly v4.10.4. Cells with a cumulative RCTD epithelial score higher than 0.5 were plotted interactively using shiny v1.9.1 to identify the barcodes of cells belonging to individual ducts. Each segmented duct was defined based on contiguous epithelial regions and manually assigned as a separate duct unit. Ducts with unclear epithelial boundaries or featuring less than 250 pseudocells were excluded. The duct-stromal interface was then defined as the cellular neighbors of duct cells as described for Open-ST niche identification.

### Open-ST duct pseudotime staging

Duct average tumor pseudotime scores computed across ductal cells were used to order individual ducts along tumor progression. Five stages were identified by binning average pseudotime scores in 0.15 intervals. RCTD scores, gene expression, ligand activity are then averaged per duct stage and scaled for visualization purposes.

### Open-ST analysis of stromal state distance to duct-stromal interface

For each stromal cell (i.e. with RCTD scores for macrophage and fibroblast states higher than 0.5), the closest duct-stromal interface was identified and their distance computed. To normalize for different duct radii and shapes, the distance to the duct interface was divided by the distance between the duct interface cells and the duct center of mass (i.e. the average x,y coordinates of ductal cells). Distances were then log-normalized so that positive scores identify extra-ductal stromal cells and negative values intraductal ones. Stromal cells further than 300 µm to any duct center were excluded.

### Open-ST cell-cell communication analysis

To investigate cell-cell communication in space, known mouse ligand-receptor pairs were obtained from CellChat v2.1.2^49^. For each ligand-receptor pair, an interaction score was computed in each cell as the geometric mean of receptor expression and average ligand expression in the 30 closest neighbors. Ligand activity scores were then computed by summing interaction scores of receptor-ligand pairs sharing the same ligand.

### Open-ST analysis of PDX model HBCx-14

Raw spatial sequencing data were processed using the same Open-ST preprocessing workflow described above. For the PDX sample, reads were aligned against a combined human and mouse reference genome to enable simultaneous detection of human tumor-derived and mouse host-derived transcripts. Following alignment, species-specific gene annotations were used to separate human (hg38) and mouse (mm10) expression profiles for downstream compartment-specific analyses. After filtering pseudocells with fewer than unique 100 genes, raw gene expression counts were log-normalized and scaled using Seurat SCTransform function as for primary Open-ST samples. The top 10 principal components have been used for Leiden clustering (resolution=0.8) and clusters were annotated based on spatial localization and quality metrics, including the overall number of transcripts and the percentage of human and mouse transcript counts quantified based on species-specific gene assignments. Clusters 3 and 7 were removed as ‘Low quality’ given their lower transcript counts and spatial mapping outside of the tissue region. Clusters 2, 8, 9 and 10 featured an enrichment for mouse transcripts and were retained for stromal analyses after removing pseudocells with less than 10 unique mouse genes or less than 50 mouse transcripts. The remaining clusters were annotated as ‘Human’. As before, mouse stromal cell-state composition was inferred using Robust Cell Type Deconvolution using the same reference dataset and parameters described for the primary Open-ST datasets. Similarly, extracellular matrix remodeling activity was quantified using AUCell v1.26.0, following the same workflow described for the primary Open-ST datasets.

### Cross-species identification of conserved CAF signatures

To identify conserved CAF transcriptional programs, we analyzed published treatment-naïve human TNBC single-cell RNA-sequencing data from Wu et al. (2021). The dataset was first restricted to patients with triple-negative breast cancer (TNBC), after which cells annotated as CAFs were selected for downstream analysis. CAFs annotated by the authors as inflammatory CAFs (iCAFs) or myofibroblastic CAFs (myCAFs) were selected from TNBC patients. To minimize biases due to unequal cell numbers, 50 iCAFs and 50 myCAFs were randomly sampled from each patient. Six TNBC patients containing at least 50 cells of each CAF subtype and were included in downstream pseudobulk analysis. Pseudobulk expression profiles were generated by summing raw gene counts across sampled cells within each patient and CAF subtype. Differential expression analysis was performed using the DESeq2 R package v1.44.0 with patient included as a blocking factor (design = ∼ patient + CAF_state). Genes with fewer than 10 counts in at least two pseudobulk samples were excluded prior to analysis. An equivalent pseudobulk differential expression analysis was performed on iCAFs and myCAFs from the mouse single-nucleus RNA-sequencing dataset using biological samples as replicates (design = ∼ sample + CAF_state). Mouse genes were mapped to their human orthologs using the orthogene R package v1.10.0. Conserved iCAF and myCAF signatures were defined as genes that were significantly differentially expressed (adjusted P < 0.05) in the corresponding CAF subtype in both human and mouse datasets after ortholog mapping. The significance of overlap between human and mouse marker genes was evaluated using one-sided Fisher’s exact test from the stats R package v4.4.3.

### Spatial mapping of conserved CAF signatures in human TNBC

To validate the spatial organization of conserved CAF programs, we analyzed an independent cohort of 94 treatment-naïve human TNBC samples profiled by poly(A)-based spatial transcriptomics (Wang et al., 2024). Pathologist-provided spot-level annotations were used to classify spatial locations based on annotation scores. Spatial spots were assigned to in situ, invasive tumor, stromal, lymphoid, or background categories when the corresponding pathology annotation score exceeded 10%. Samples containing more than five in situ and more than five invasive tumor spots were selected for paired analysis, resulting in 12 TNBC samples. Conserved iCAF and myCAF signatures identified from the cross-species single-cell analysis were projected onto spatial transcriptomic spots using the AUCell R package (v1.28.0) as described in ‘Gene set scoring in single nuclei and spatial data’. Briefly, genes were ranked according to their raw expression counts using AUCell_buildRankings(), and signature enrichment was calculated using AUCell_calcAUC() with aucMaxRank set to 5% of the ranked genes. AUCell scores were standardized across spatial spots by z-score scaling and matched to pathology annotations based on spatial coordinates. Spatial distributions of CAF signatures were visualized using annotated spatial coordinates. For statistical analysis, patient mean iCAF and myCAF signature scores were calculated for each annotation group. Paired comparisons between in situ and invasive tumor regions were performed using two-sided Wilcoxon signed-rank tests. Paired visualization and statistical annotation were performed using the ggpubr R package (v0.6.2).

---

**Supplementary Figure 1 related to Figure 1.**
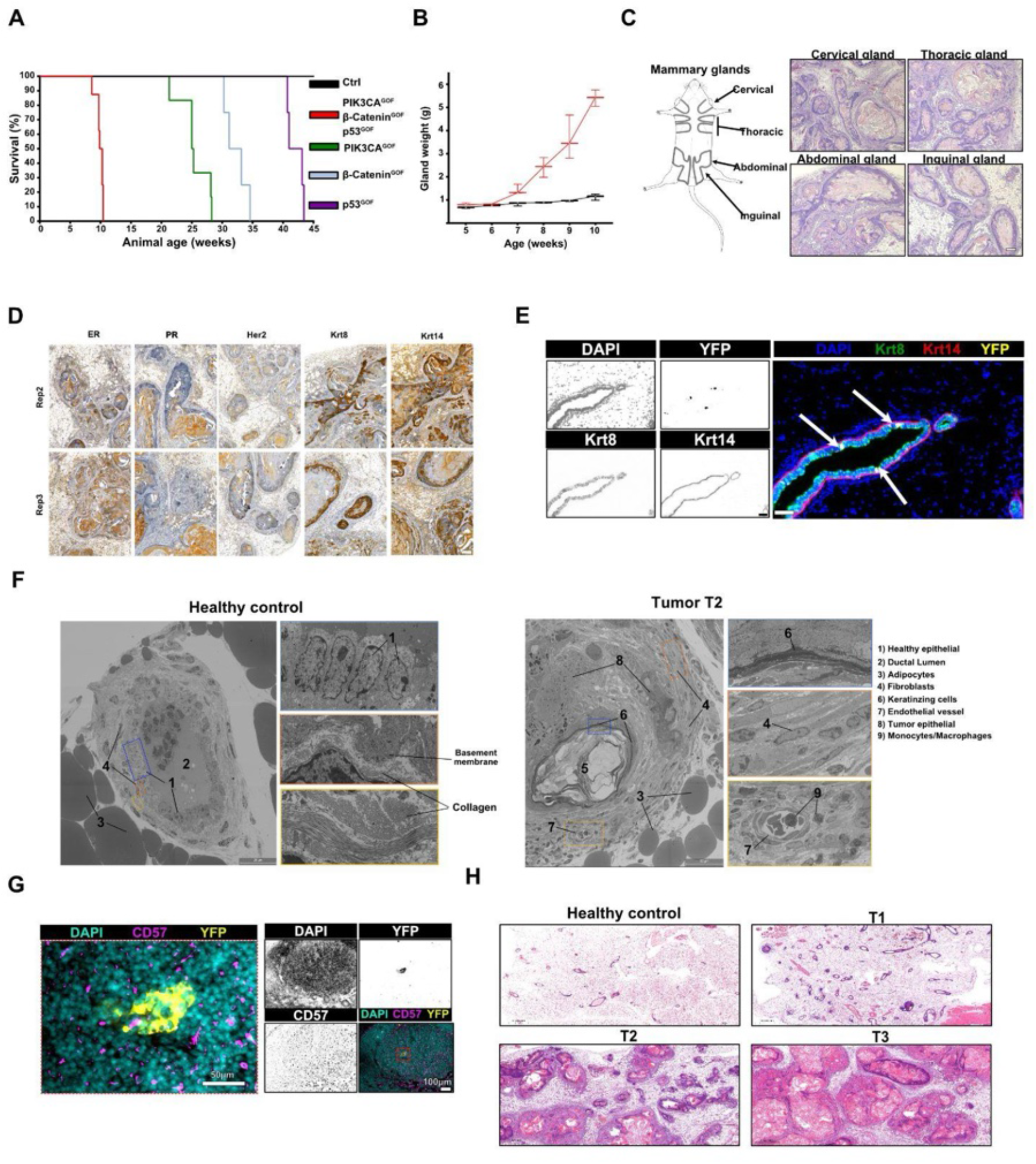
(A) Tumor growth is detectable after 6 postnatal weeks. Combined weight of thoracic and abdominal mammary glands in mutant and control (WAP-iCre and YFP but no oncogenic transgenes) animals. N = 3 per timepoint. (B) Tumors develop consistently across all mutant animals and tumor development is delayed in single-mutant animals. Kaplan–Meier survival curves comparing tumor onset in animals carrying individual oncogenic alleles to triple-mutants and control animals (all oncogenic transgenes but no WAP-iCre system). N = 8 per group. (C) Tumors arise across all mammary glands. Left: murine mammary gland anatomy. Right: H&E staining of cervical, thoracic, abdominal and inguinal glands from a mutant animal at postnatal week 8. Scale bar: 100 µm. (D) As in Fig. 1B, immunohistochemistry staining of mammary tumors in additional replicates at 9 postnatal weeks. Scale bars: 100 µm. (E) Oncogene activation (YFP^+^) is restricted to the luminal compartment (Krt8^+^/Krt14^-^) in early-stage glands. Single channel (left) and merged (right) immunofluorescence (IF) images of mammary glands from a mutant animal at postnatal week 5. Arrows: YFP⁺/Krt8^+^ cells. Scale bars: 50 µm. (F) Transmission electron microscopy of mammary gland ducts derived from healthy and tumor-bearing animals. T2 (8 postnatal weeks) tumor lesions show disrupted epithelial architecture, dense fibroblasts layer around the tumors and multiple macrophages infiltrating the stroma. (G) Tumor cells (YFP^+^) are detected in intramammary lymph nodes at study endpoint. Zoom-in merged (left) and overview (right) merged and single-channel IF images of lymph node tissue from mutant animals at 10 postnatal weeks. Scale bars: 100 µm (overview), 50 µm (zoom-in). (H) Histology across study timepoints. H&E staining of mammary glands collected at Healthy control (12 weeks), T1 (5–6 weeks), T2 (8–9 weeks), and T3 (10–11 weeks). Scale bars: 500 µm.

**Supplementary Figure 2 related to Figure 2.**
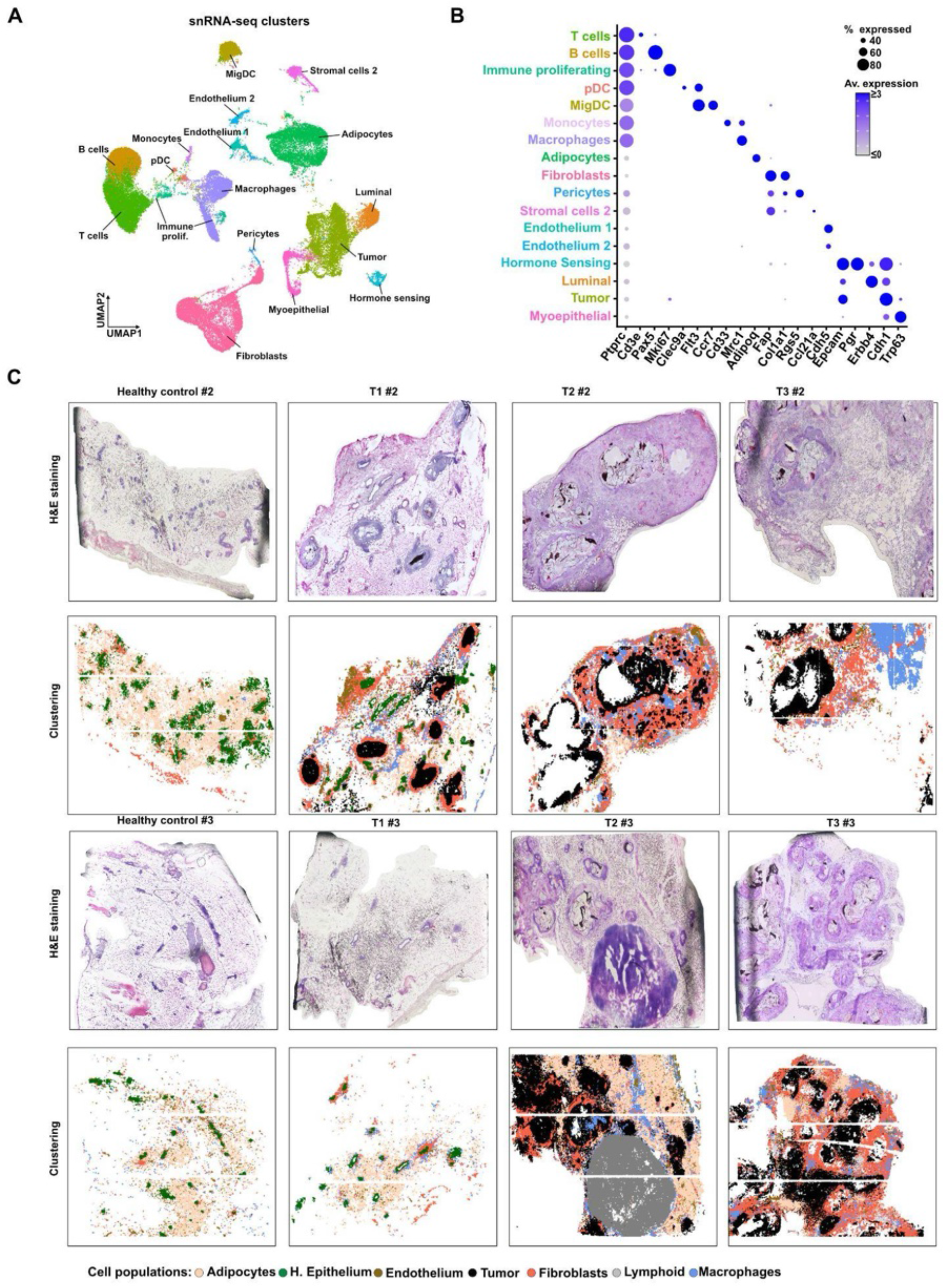
(A) Uniform manifold approximation and projection (UMAP) plot of healthy and transformed mammary gland cell types. RNA from *N* = 58,742 nuclei was sequenced (12 samples, 3 biological replicates/timepoint, Methods). (B) Marker gene expression across cell types. Dot size: percentage of cells (≥20% shown), for each cell type, where ≥1 transcript of a specific gene was detected. Color: Z-scores of mean gene expression values per cell type. (C) As in Fig. 2B, H & E images of tissue slices (top) and unbiased clustering results of the genome-wide, high-resolution spatial transcriptomics data (Open-ST) for biological replicates not shown in Fig. 2B.

**Supplementary Figure 3 related to Figure 2.**
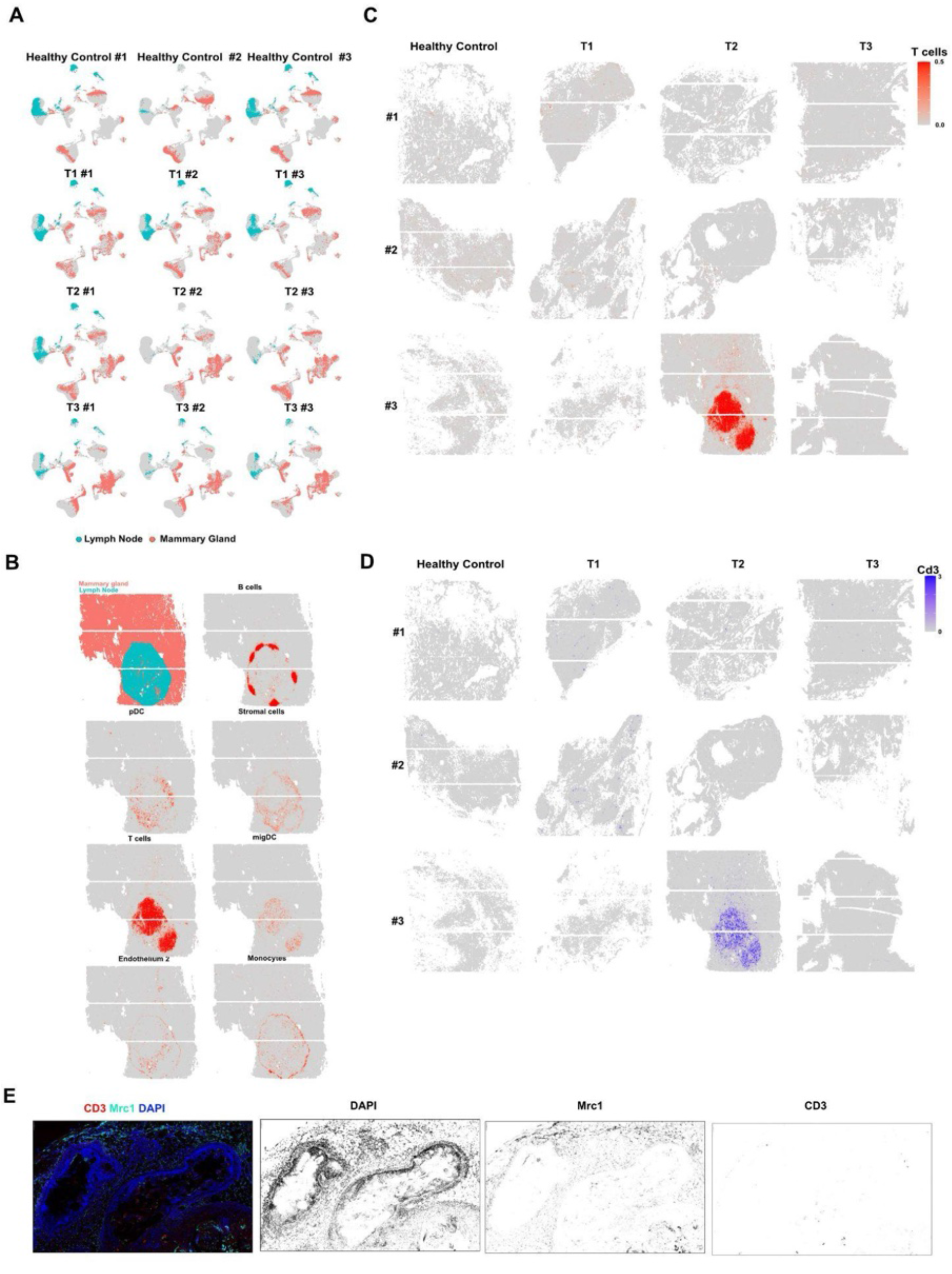
(A) UMAPs of the individual snRNA-seq samples can contain lymph node–associated immune and stromal populations (blue). snRNAseq and Open-ST replicates collected from independent mammary glands. (B) RCTD deconvolution scores (as in Fig. 2C, Methods) from lymph-node associated snRNAseq populations map to the intramammary lymph node captured in Open-ST sample T2-#3. (C) T cells are scarce in the healthy and peritumoral mammary stroma. Pseudocells are colored by T cell RCTD deconvolution scores. (D) CD3 expressing cells are scarce in the healthy and peritumoral mammary stroma. Pseudocells are colored by aggregate Cd3 (Cd3d, Cd3e and Cd3f) gene expression. (E) CD3+ cells are scarce in the healthy and peritumoral mammary stroma. Single channel (left) and merged (right) immunofluorescence (IF) images of macrophages (Mrc1+) and T cells (Cd3+) in the mammary gland from a mutant animal at postnatal week 8. Scale bars: 50 µm.

**Supplementary Figure 4 related to Figure 3.**
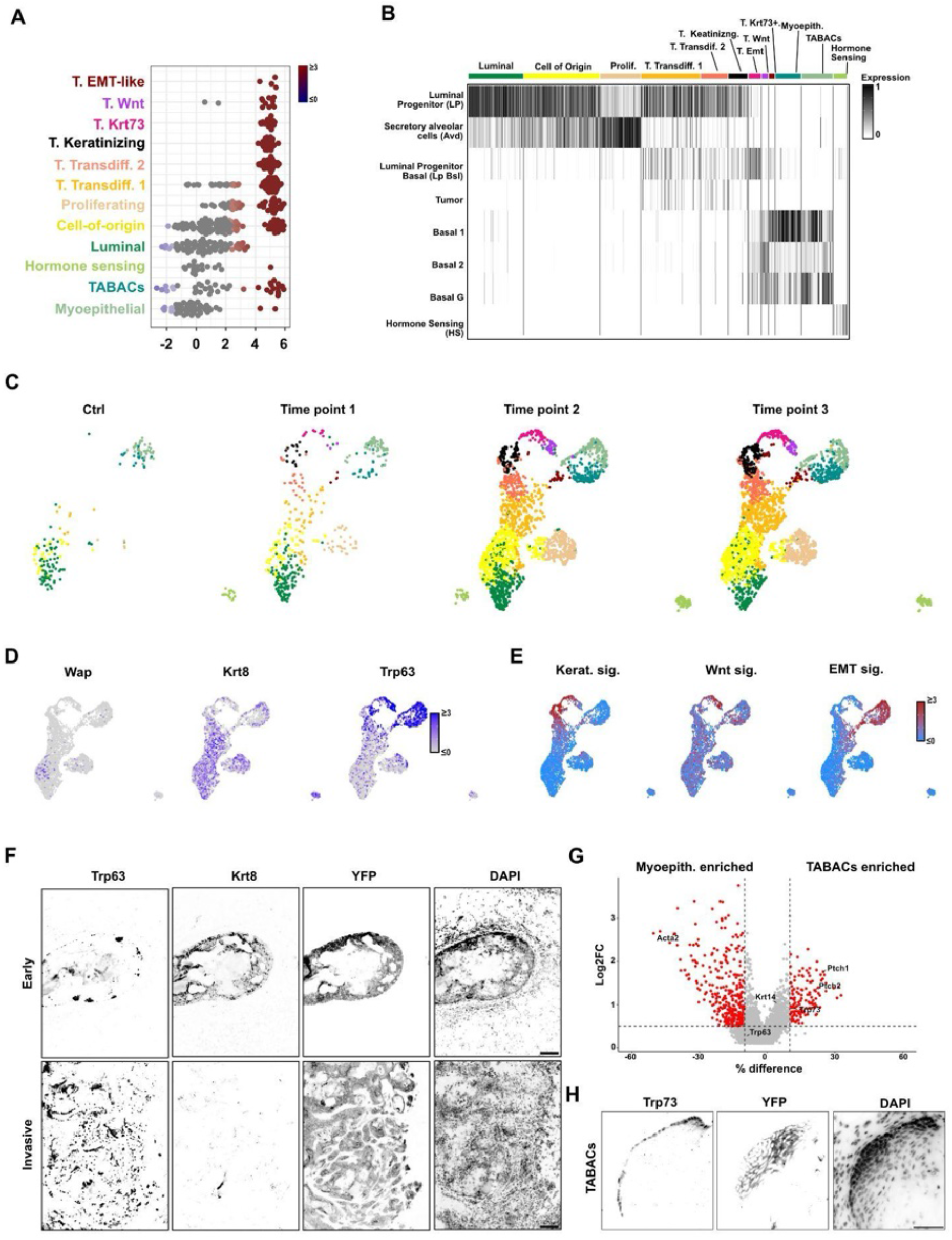
(A) Quantification of differential neighborhood abundance across epithelial cell states. Differential abundance analysis (as in Fig. 3A) showing the distribution of neighborhood log fold changes between tumor-bearing (T1–T3) and healthy samples for each epithelial cell state. Each point represents one neighborhood; colored points indicate neighborhoods with significant differential abundance. (B) Comparison of epithelial cell-state annotations with a published murine mammary gland tumor atlas. Heatmap showing label transfer scores from the reference atlas of Bach *et al.*^38^ across the epithelial populations identified in this study (Methods). Columns correspond to epithelial cell states identified in our dataset, and rows to reference populations. (C) Uniform manifold approximation and projection (UMAP) of epithelial single-nucleus RNA-seq data (as in Fig. 3C), colored by epithelial cell-state annotations and split by experimental timepoint. (D) Luminal-to-basal transdifferentiation is associated with progressive changes in marker gene expression. UMAPs (as in Fig. 3C) colored by Z-scores for expression of the luminal marker Krt8, the basal marker Trp63, and Wap, which marks the luminal cell-of-origin. (E) Basal-like tumor states activate distinct transcriptional programs. UMAPs showing AUCell enrichment scores for HALLMARK pathways keratinization, Wnt/β-catenin signaling, and epithelial-to-mesenchymal transition (EMT) gene signatures (Methods). (F) Single-channel IF images corresponding to Fig. 3E. Scale bars: 50 µm. (G) Volcano plot comparing gene expression between TABACs (right) and myoepithelial cells (left). Selected differentially expressed genes are highlighted. X: Difference in the percentage between TABACs and myoepithelial cells where ≥1 transcript of a specific gene was detected, Y: log2 fold change in average gene expression. (H) Single-channel IF images corresponding to Fig. 3G. Scale bars: 50 µm.

**Supplementary Figure 5 related to Figure 3.**
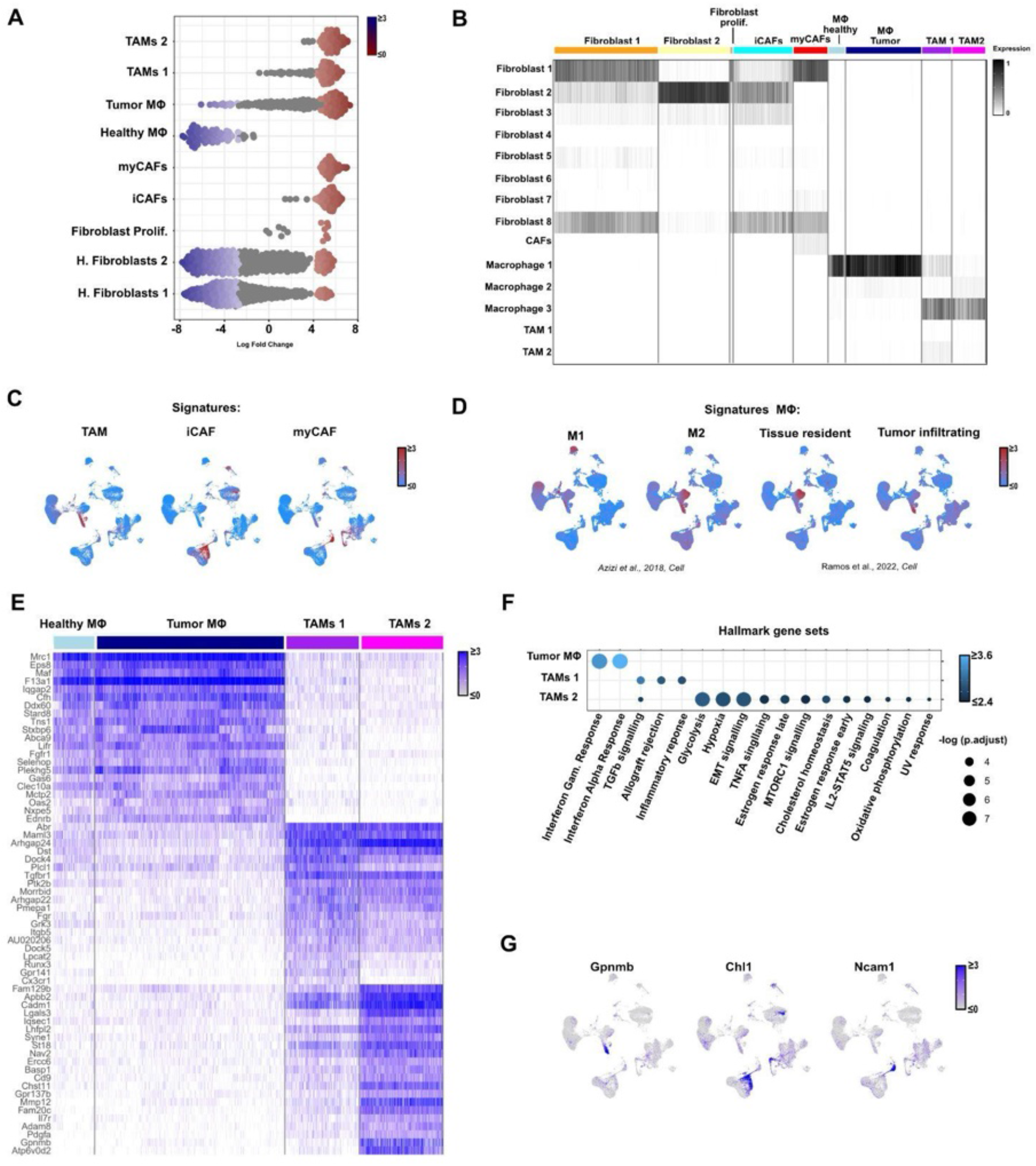
(A) Quantification of differential neighborhood abundance across stromal cell states. As in SFig. 4A but for fibroblast and macrophage populations. Each point represents one neighborhood; colored points indicate neighborhoods with significant differential abundance. (B) As in SFig. 4B, comparison of stromal cell-state annotations with a published mammary gland atlas. (C) Stromal populations acquire published cancer-associated transcriptional programs. UMAPs colored by AUCell enrichment scores for published tumor-associated macrophage (TAM), inflammatory cancer-associated fibroblast (iCAF), and myofibroblastic CAF (myCAF) gene signatures described by Mayer et al. (2023). (D) Macrophage populations display distinct polarization and activation programs. UMAPs colored by AU Cell scores (Methods) for M1 and M2 macrophage signatures (Azizi et al., 2018) together with tissue-resident and tumor-infiltrating macrophage signatures (Ramos et al., 2022). (E) Marker gene expression across macrophage populations. Heatmap showing normalized expression of the top 20 marker genes for tumor macrophages, TAM1, and TAM2 (Methods). (F) Hallmark pathway enrichment across tumor-associated macrophage populations. Dot plot summarizing significantly enriched Hallmark gene sets identified by differential expression analysis. Dot color represents normalized enrichment score, and dot size indicates statistical significance (−log adjusted P value). (G) Remodeled stromal populations upregulate specific marker genes. UMAPs (as in Fig. 3H) colored by Z-scores for expression of Gpnmb, Chl1, and Ncam1, marking TAMs, iCAFs, and myCAFs, respectively.

**Supplementary Figure 6 related to Figure 4.**
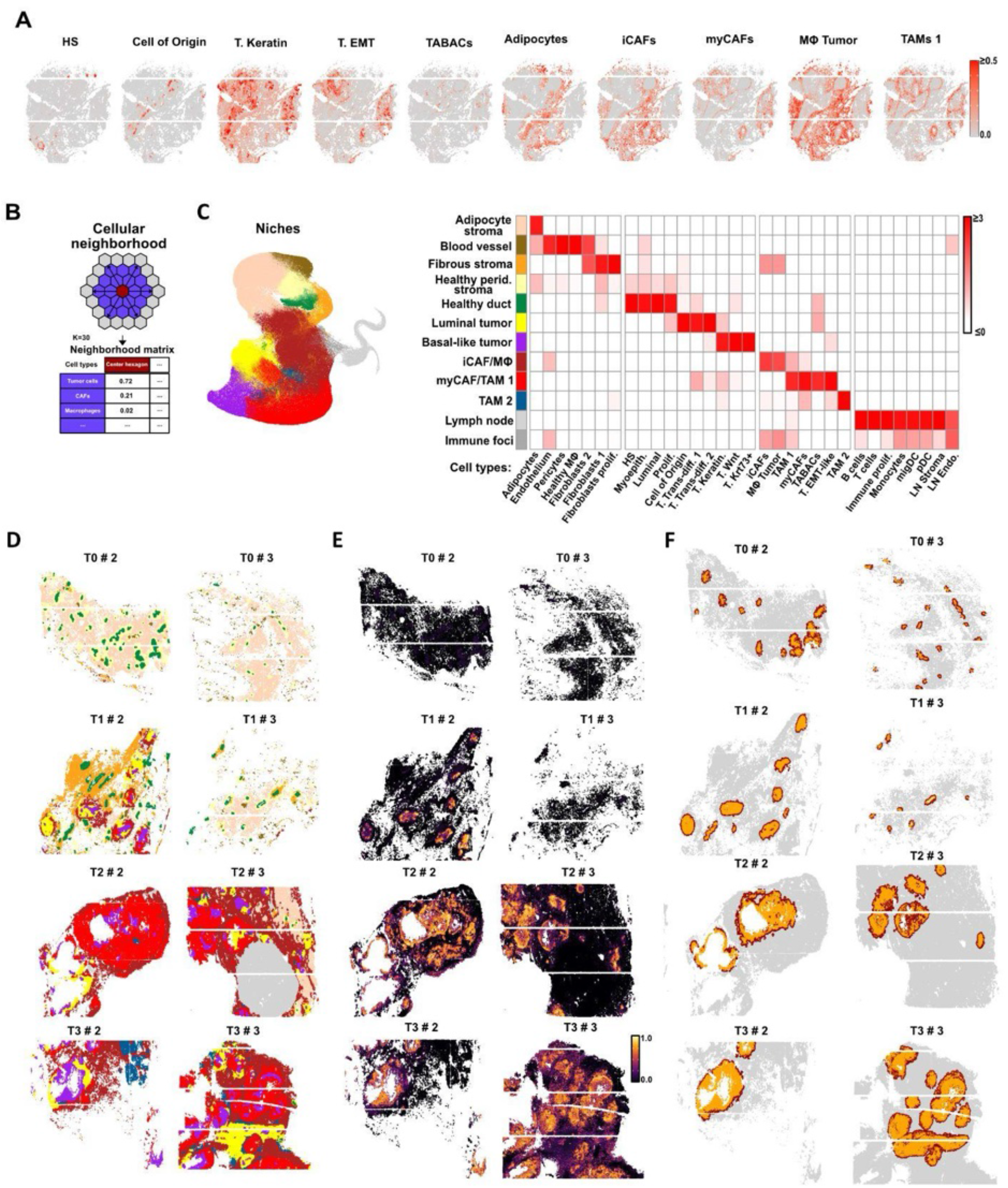
(A) Single-nucleus RNA-seq epithelial and stromal states mapped to spatial transcriptomics. RCTD deconvolution scores (see Fig. 2C, Methods) for selected epithelial and stromal populations are shown for Open-ST sample T2 #1. (B) Definition of cellular neighborhoods. RCTD deconvolution scores are averaged per cellular neighborhood (*i.e.,* 30 closest pseudocells). (C) Epithelial and stromal populations are spatially organized. Cellular neighborhoods with similar cell state deconvolution scores are grouped into niches (Methods). Left: UMAP colored by niche identity. Right: Heatmap of Z-scores for mean RCTD deconvolution scores of epithelial, stromal, and immune populations across all identified multicellular niches. (D) Spatial mapping of multicellular niches across additional biological replicates not shown in Fig. 4A. Color legend in B. (E) Spatial mapping of tumor pseudotime scores across additional biological replicates not shown in Fig. 4B. (F) Spatial segmentation of individual ducts across additional biological replicates not shown in Fig. 4C.

**Supplementary Figure 7 related to Figure 4.**
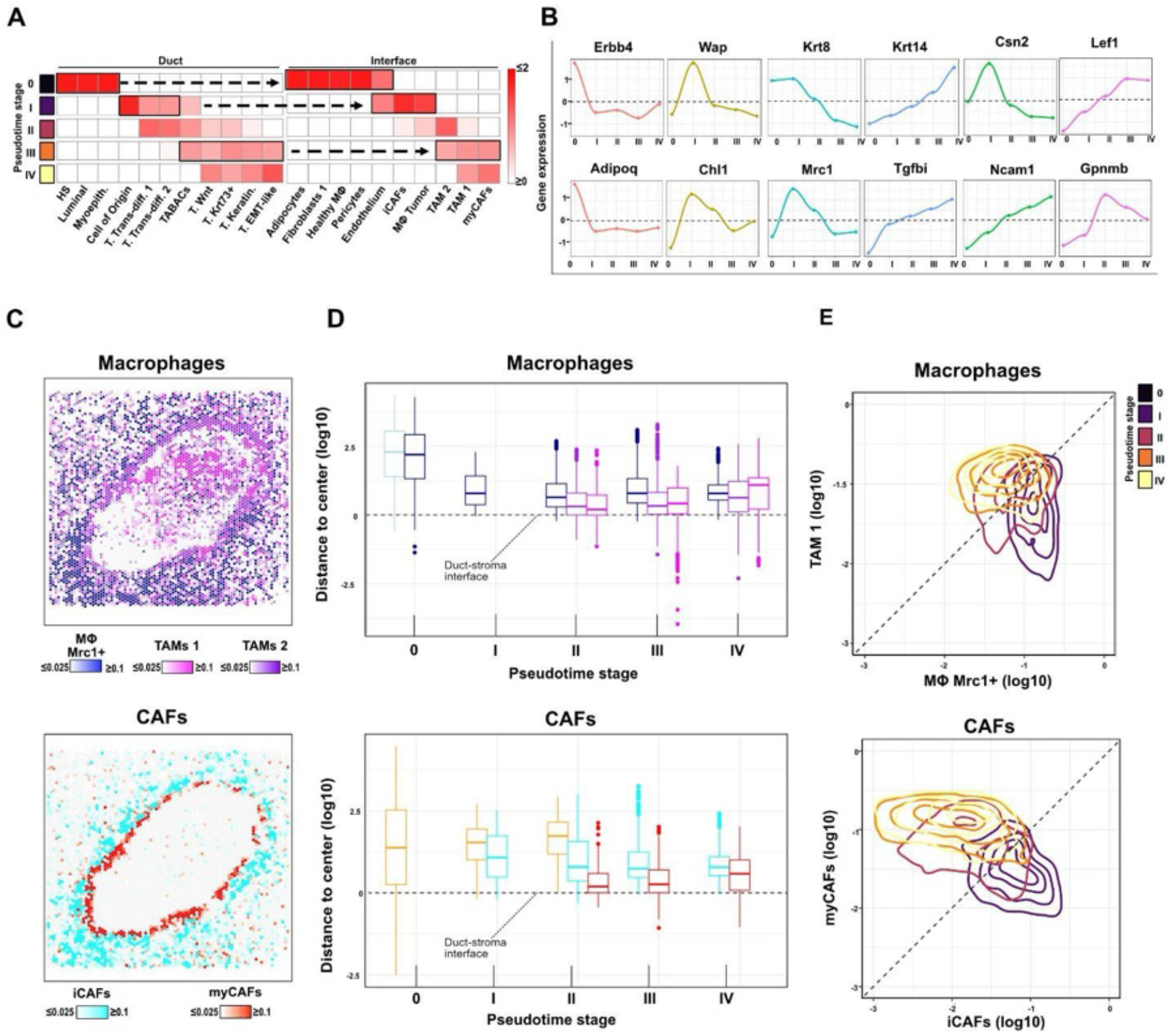
(A) Specific stromal compositions at the duct-stromal interface follow specific epithelial phenotypes within ducts. Cell-state composition (Z-scores of mean RCTD deconvolution scores) of epithelial populations in the duct (as in Fig. 5B) and stromal populations in the duct-stromal interface (as in Fig. 5C). (B) Marker gene expression follows population dynamics. Z-scores of mean expression of epithelial marker genes in the duct compartment and stromal markers genes in the duct-stromal interface. (C) Stromal phenotypes are spatially organized in the periductal stroma. Spatial transcriptomics of the same duct shown in (C), displaying macrophage (top) and CAF (bottom) cell-state composition. (D) Spatial organization of stromal phenotypes across duct stages (time). Box plots show the distance of individual macrophage (top) and fibroblast (bottom) populations to the tumor–stroma interface (Methods). Boxes omitted for populations with <150 stage-specific pseudocells. Color legend as in (D). (E) Progressive remodeling of stromal states at the tumor-stromal interface. Phase diagrams of stromal dynamics during tumor progression. Cell-state composition of neighborhoods at the duct-stromal interface (here shown in log10 scale) are used to compute 2D kernel densities for each duct stage. Stage 0 not shown as remodeled stromal populations only appear later (see (C).

**Supplementary Figure 8 related to Figure 5.**
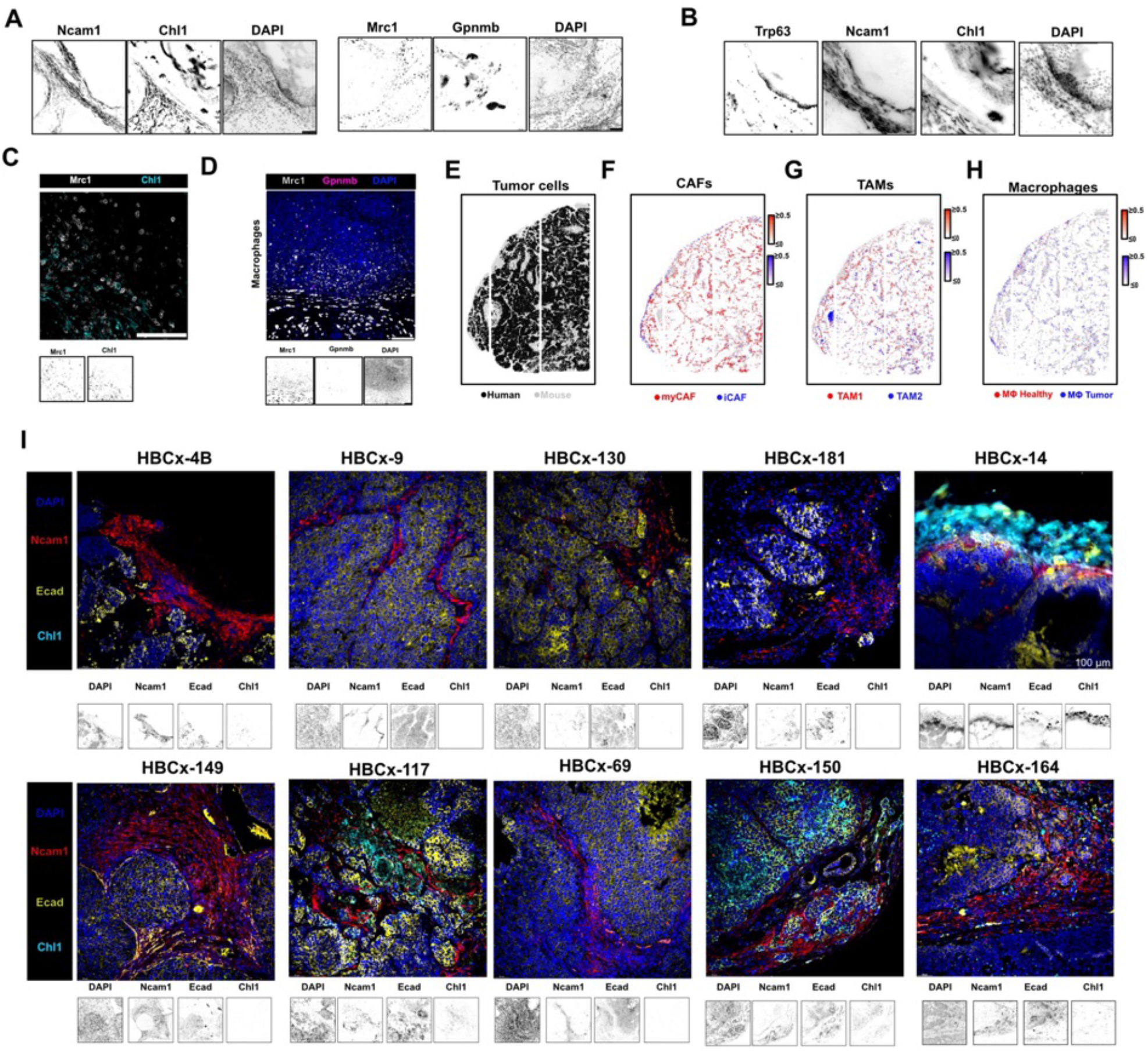
(A-B) Single-channel IF images corresponding to Fig. 5 A-B. Scale bars: 50 µm. (C) iCAFs are closely associated with macrophages in the periductal stroma, from T2. IF staining of iCAFs (Chl1⁺) and macrophages (Mrc1⁺) in a T2 tumor. (D) Cell type distribution at the duct-stromal interface. IF staining of macrophages (Mrc1⁺), and TAMs (Gpnmb⁺) a PDX (#3) model (right). Scale bars: 50 µm (E) As in Fig. 5D, Open-ST analysis of one TNBC PDX model (HBCx-14). Species-aware transcript assignment distinguishes human tumor-cell regions from mouse stromal regions. (F-H) Deconvolution scores in mouse pseudocells for CAFs (F), TAMs (G) and macrophages (H) showing conserved stromal remodeling (RCTD, Methods). (I) Validation of stromal remodeling in 10 TNBC patient-derived xenograft (PDX) models with diverse genetic drivers (STable 6, Coussy et al., 2019). As in (A), IF distribution of stromal markers (Chl1 iCAF, NCAM1 myCAF) at the tumor (marked by Ecad)-stromal interface.

**Supplementary Figure 9 related to Figure 5.**
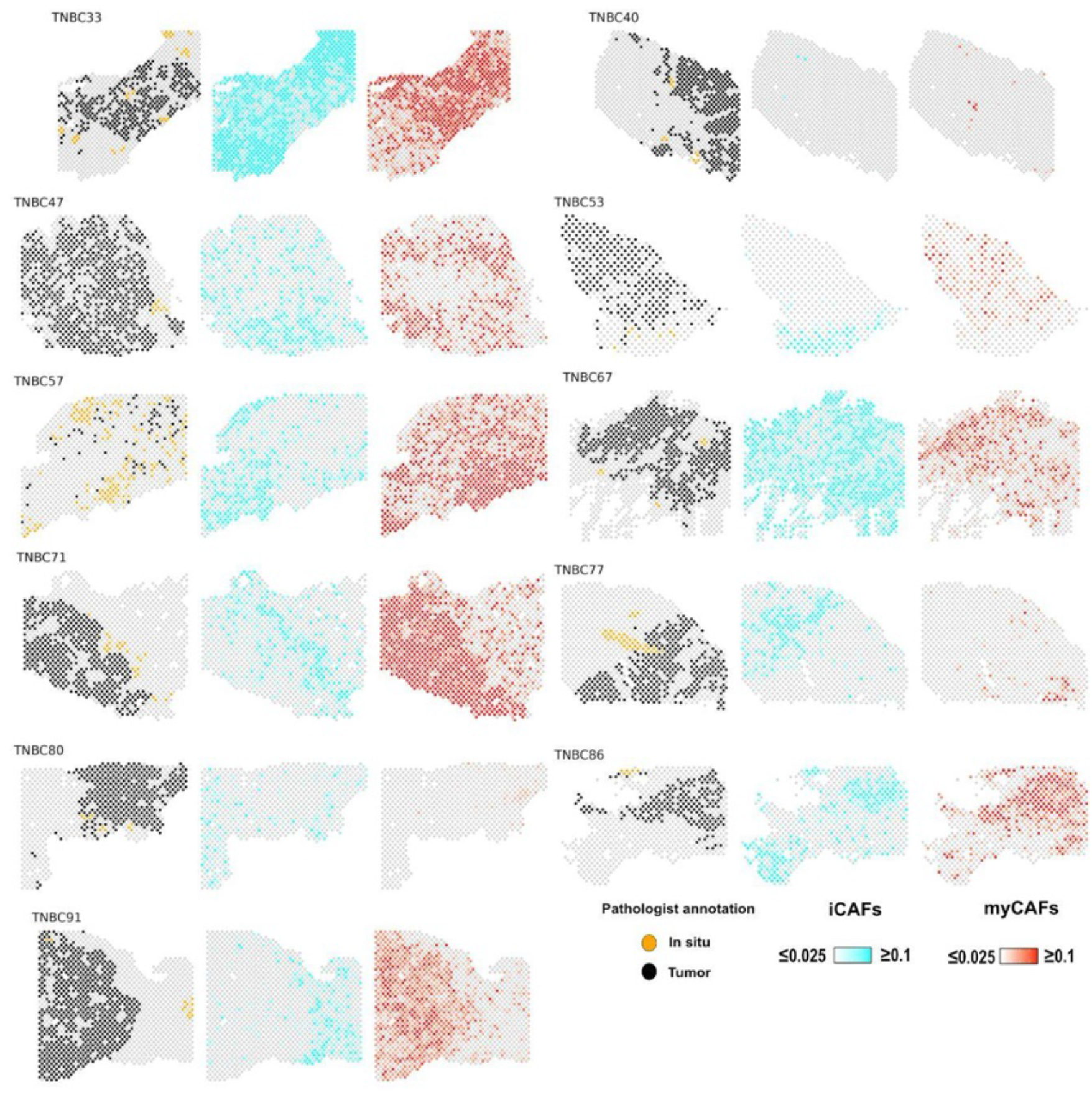
As in Fig. 5G, spatial projection of conserved stromal signatures in human TNBC across additional 11 patients (low-resolution spatial transcriptomics data from Wang et al 2024) with pathologist’s annotation of in situ and invasive tumor regions (left) together with spatial AU Cell scores of conserved iCAF and myCAF signatures.

**Supplementary Figure 10 related to Figure 6.**
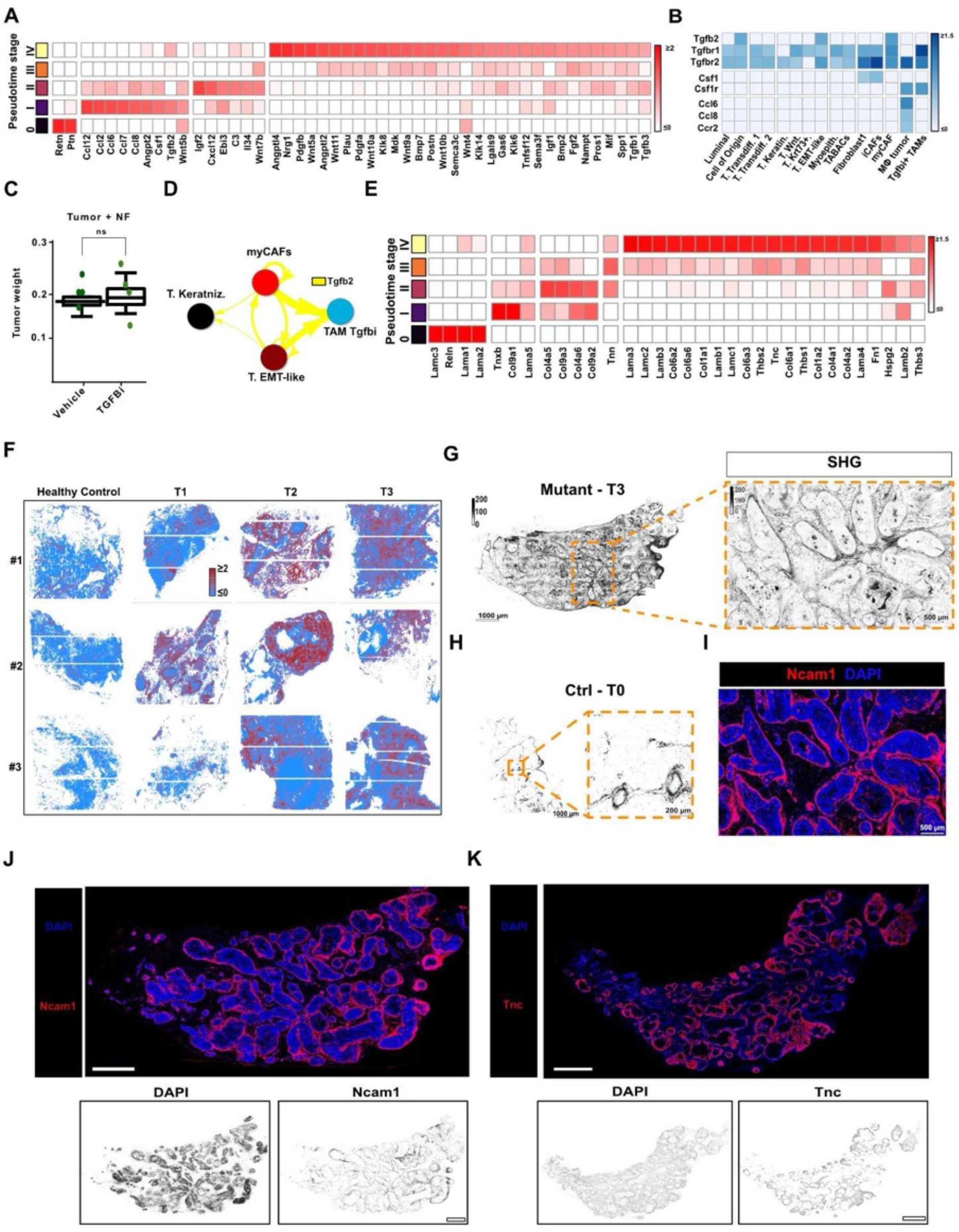
(A) Stage-specific secreted interactions at the tumor-stromal interface. Heatmap: Z-scores of mean spatial ligand activity (computed for spatial pseudocells, as in Fig. 6B). Stage-specificity is defined by log2FC >0 in average ligand activity and non-zero score in at least 5% of stage-specific interface pseudocells. (B) Early interface ligand and receptor expression in single-nuclei populations. Heatmap: average expression of selected early-stage ligands and their receptors across snRNA-seq populations. (C) Treatment of co-injected lesions does not affect the tumor weight at study termination. Fatpad weights (in gram) at the day of study termination for Groups A, B. Bars: endpoint tumor mass in grams. *N* =4. (D) myCAFs and EMT tumor cells secrete Tgfb2 towards TAMs at the late tumor-stromal interface. As in Fig. 6C but showing Tgfb2 interactions in snRNA-seq population mapping to the advanced (stage III/IV) tumor–stromal interface. (E) Stage-specific ECM ligand-receptor interactions at the tumor-stromal interface. As in (A) but related to ECM ligands. (F) ECM remodeling at the tumor-stromal interface. As in Fig. 6J but spatial, AU Cell Z-scores for the REACTOME ‘ECM organization’ pathway across Open-ST samples. (G) Collagen is visualized at the tumor-stromal interface. Second harmonic generation (SHG) imaging of collagen fibers in healthy control (left) and T3 tumor (right) tissue sections. Zoom-out and zoom-in views shown. Scale bars: 1000 µm (overview), 200 µm (inset). (H) Collagen fibers colocalize with myCAFs at the tumor-stromal interface. IF and SHG imaging of the same tumor region from consecutive sections. Top: Ncam1⁺ myCAFs shown by IF. Bottom: corresponding SHG image showing collagen deposition in the same field. Scale bars: 50 µm. (I) Whole-gland immunofluorescence image corresponding to (F). Merged and single-channel images for Ncam1 and DAPI are shown in a T3 tumor section. (J) Whole-gland immunofluorescence merged image and single-channel images for Tnc and DAPI are shown in examples from healthy and all tumor stages T1-T3 tumor section.

**Supplementary Figure 11 related to Figure 6.**
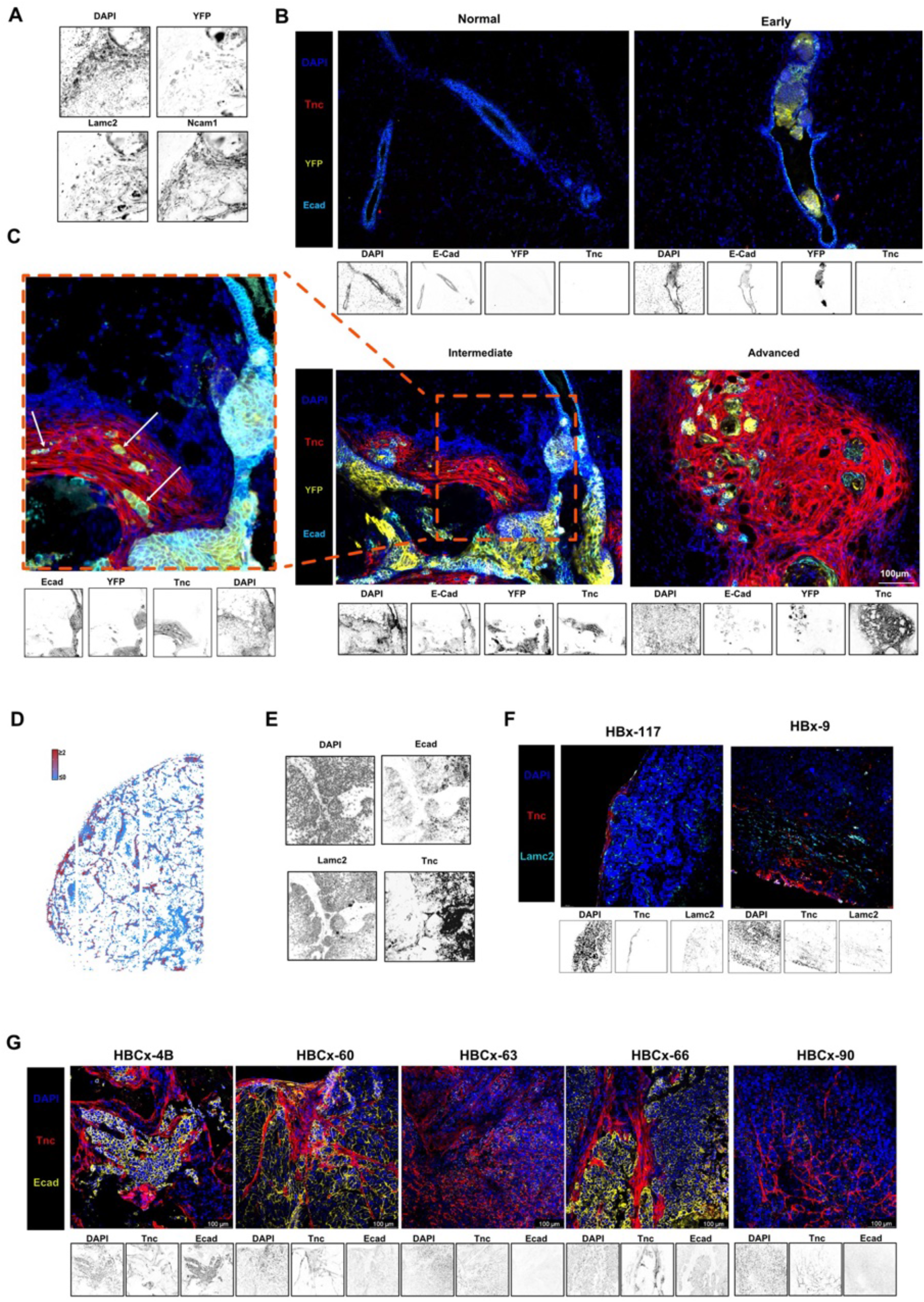
(A) Individual channels of IF images shown for Ncam1⁺, Lamc2, YFP, and DAPI corresponding to Fig. 6L. Scale bars: 50 µm. (B) Merged and single-channel immunofluorescence images for Ecad, Tnc and DAPI in representative normal, early, intermediate and advanced lesions. Scale bars: 100 µm. (C) Zoom in of Merged and single-channel immunofluorescence images for Ecad, Tnc and DAPI in intermediate lesions showing the infiltrating tumor cells (arrows) in TNC positive stroma Scale bars: 100 µm. (D) Spatial ECM pathway scoring in myCAF enriched regions in PDX model. AU Cell Z-scores for the REACTOME ‘ECM organization’ pathway across Open-ST samples. (E) Individual channels of IF images shown for Tnc⁺ myCAFs, Lamc2, Ecad, and DAPI corresponding to Fig. 6M. Scale bars: 50 µm. (F) Conserved expression of Lamc2 and TNC in PDX replicates. (G) Immunofluorescence images and single-channel images for Ecad, Tnc and DAPI are shown in a panel of human TNBC PDXs sections. Scale bars: 100 µm.

**Supplementary Figure 12 related to Figure 7.**
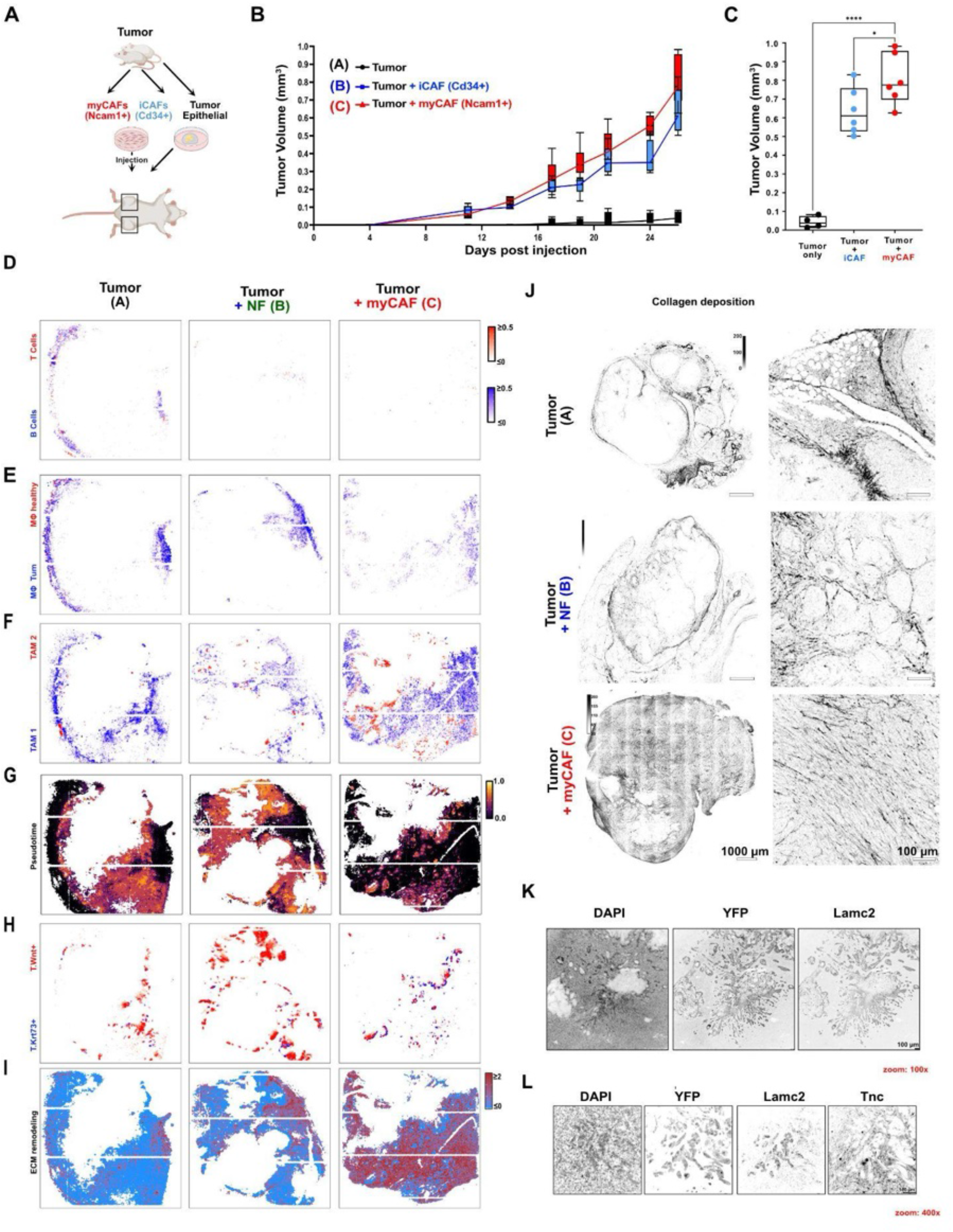
(A) Experimental design for orthotopic co-transplantation. Tumor cells were injected with healthy fibroblasts (Group A), with tumor-derived Cd34+/ Ncam1 negative (Group B), or with tumor-derived Cd34negative/ Ncam1⁺ myCAFs (Group C) into the mammary fat pad of immunocompetent FVB/N mice. (B) Both myCAFs and iCAFs promote tumor growth when co-transplanted with tumor cells in healthy immunocompetent mice. Tumor volume was measured daily from days 8 - 26, the experimental endpoint defined by maximum allowable tumor size. Tumor growth at individual injection sites is shown over time in mm³ (N = 4 per group). (C) myCAFs and iCAF co-injection leads to increased tumor volume at study termination. Fatpad size (in mm^3^) at the day of study termination for Groups A, B, and C. *N* = 4. (D - F) Spatial mapping of lymphocytes (D), macrophages (E) and TAM (F) snRNA-seq populations in Open-ST transplant samples (as in Fig. 7C). (G) Spatial mapping of tumor pseudotime (as in Fig. 4B) but related to transplant samples in (E-G). (H) As in (E-G) but related to Wnt+ and Krt73+ tumor populations. (I) myCAFs promote ECM remodeling. As in Sup Fig. 10 F but related to transplant tumors in C. J) myCAFs promote the deposition of a pro-invasive ECM. Second harmonic generation imaging of collagen deposition at tumor injection sites for Groups A–C. For each group, a zoomed-out view (scale bar: 1000 µm) and a corresponding zoom in (scale bar: 100 µm) are shown. Collagen fiber organization and density are visualized across the tumor and surrounding stroma K) Immunofluorescence staining of tumors from group (A - C). Channels: DAPI, YFP, Ncam1. Scale bars: 100 µm. L) Single immunofluorescence channels related to Fig. 7E. Single-channel immunofluorescence images corresponding to Fig. 7E. Channels: Lamc2, YFP, Tnc, and DAPI. Scale bars: 100 µm.

**Supplementary Figure 13 related to Figure 7.**
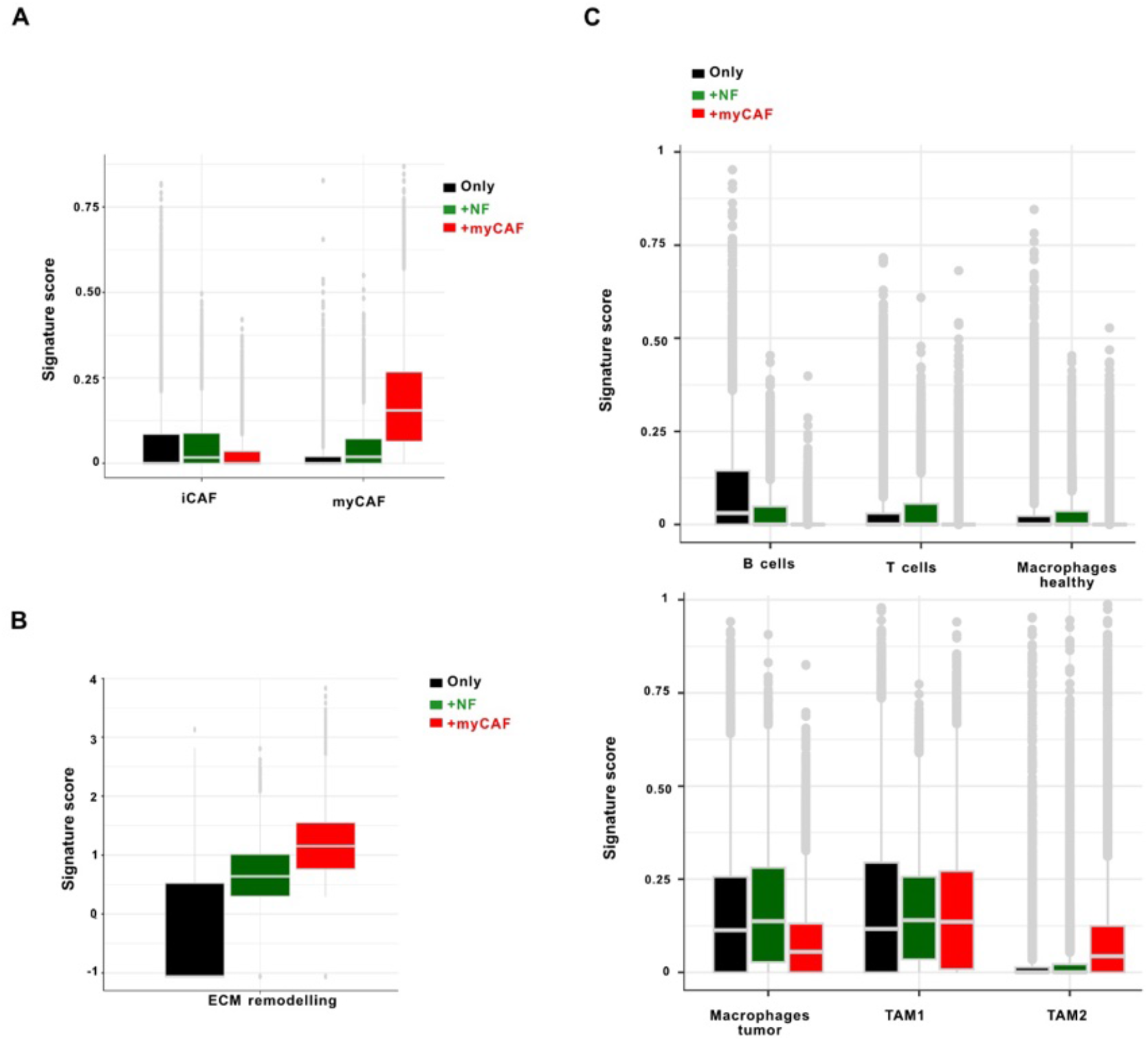
(A) Quantification of CAF phenotypes across conditions. Boxplot: RCTD scores per tumor pseudocells (tumor pseudotime > 0.3). (B) Quantification of ECM pathway across conditions. Boxplot: RCTD scores per tumor pseudocells (tumor pseudotime > 0.3). (C) Quantification of immune cells phenotypes across conditions. Boxplot: RCTD scores per tumor pseudocells (tumor pseudotime > 0.3).

## Method Figures

**Supplementary Method Figure 1 related to Figure 2.**
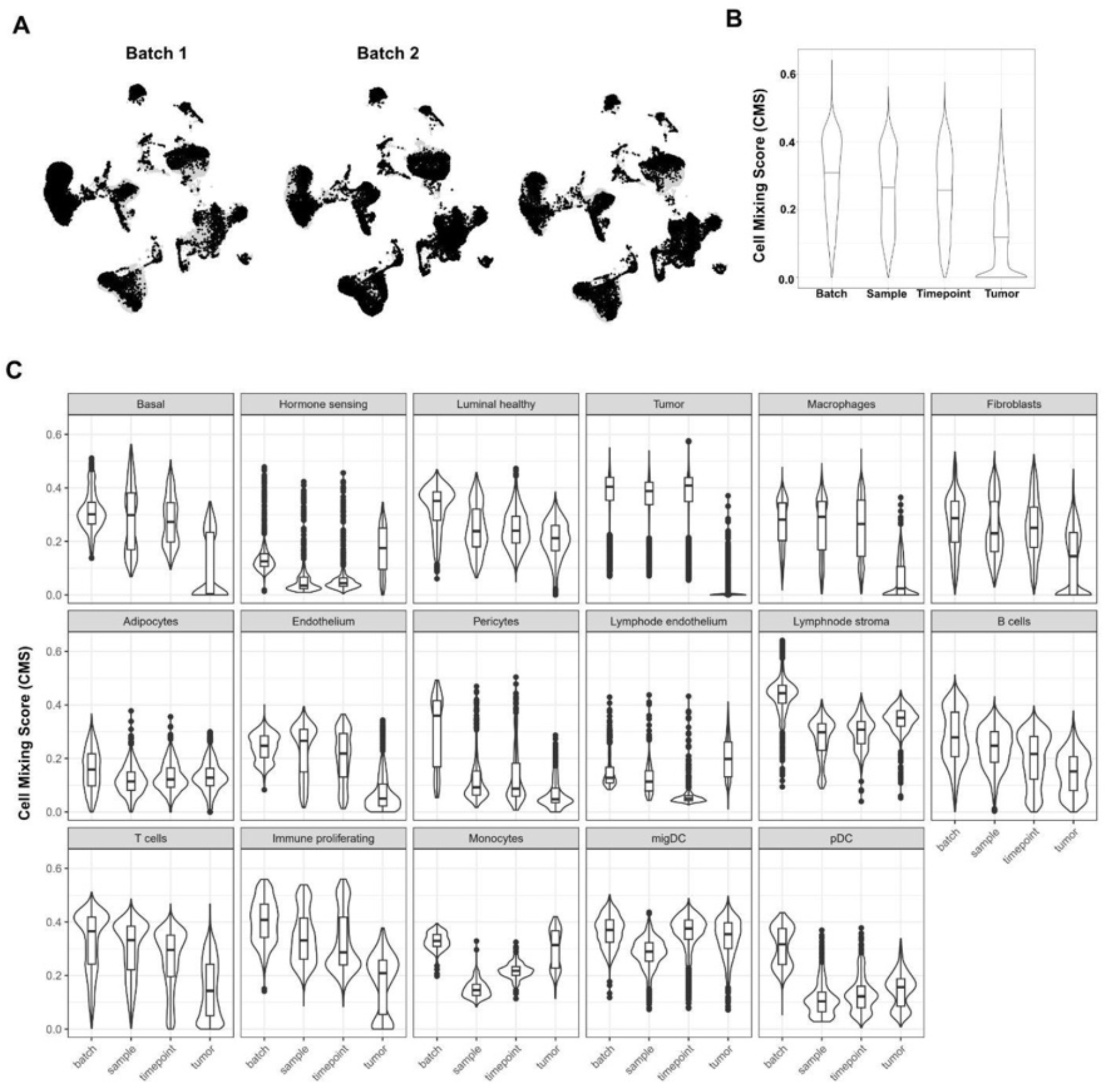
(A) UMAPs of individual batches in the snRNA-seq data. Black: cells from the given batch, grey: cells from other batches. (B) Quantification of batch effects in the snRNA-seq data. Distribution of cell mixing scores across sequencing batches, biological samples, experimental timepoints, and tumor status (Methods). Higher values indicate stronger local mixing of cells across the corresponding grouping variable, whereas lower values indicate greater separation. Batch, timepoint and samples exhibit higher mixing than tumor status, arguing for the presence of biological differences over batch effects. (C) As in (B), cell mixing scores by cell type.

**Supplementary Method Figure 2 related to Figure 3.**
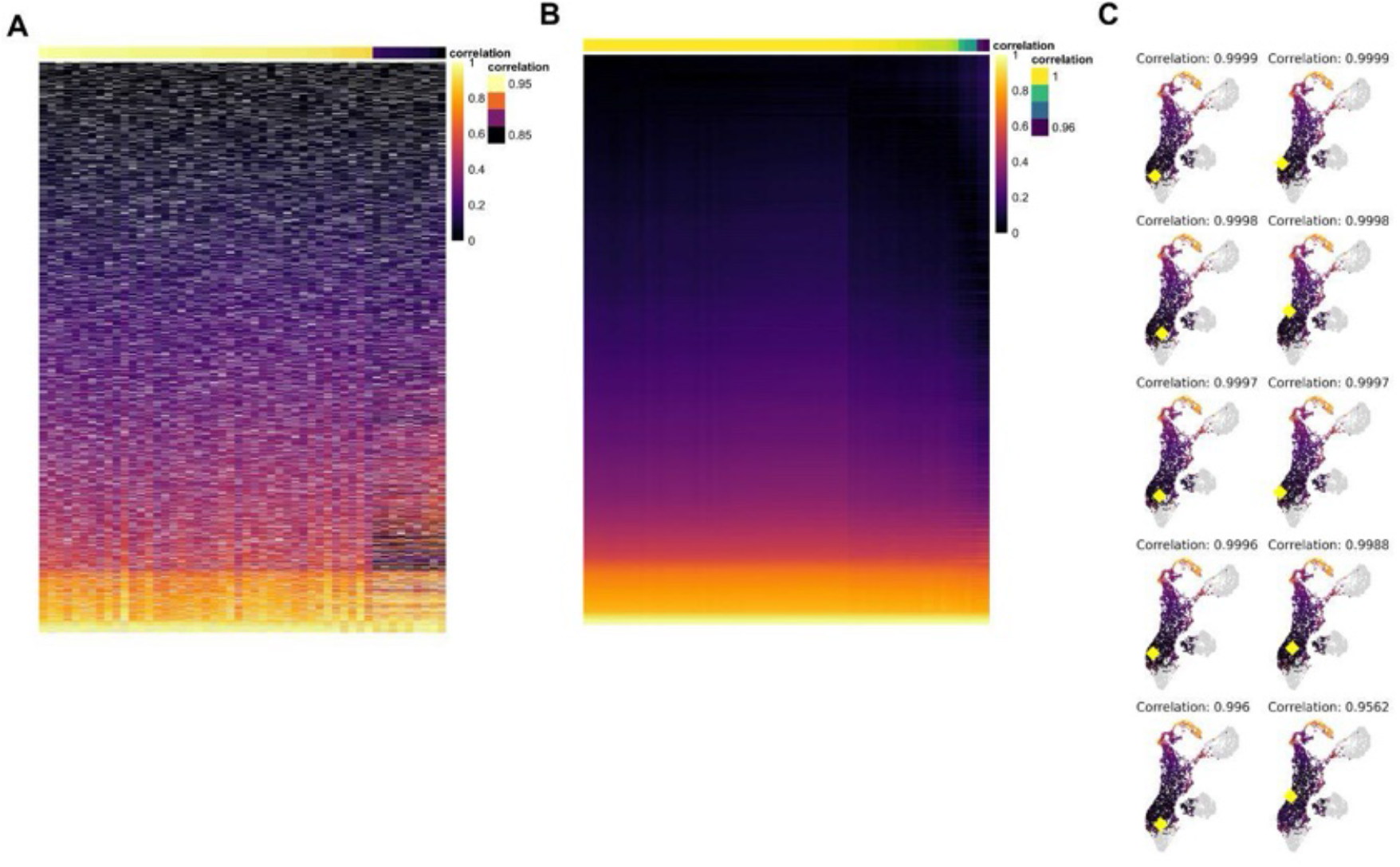
(A) Tumor pseudotime ordering is robust to bootstrapping. Correlation matrix showing the similarity of pseudotime assignments generated from 50 bootstrapping iterations. Columns correspond to one run where 20% of cells were randomly removed (indicated with white dashes) and color intensity represents the pseudotime score. Columns are ordered by decreasing Spearman correlation to the pseudotime score in Fig. 3D. (B) Tumor pseudotime ordering is robust to the choice of root cell. All 66 Wap-positive cells were used independently as starting points for trajectory inference. Correlation matrix showing the similarity of pseudotime assignments generated from all 66 alternative Wap-expressing root cells. Columns correspond to individual root-cell-derived trajectories, and color intensity represents the pseudotime score. Columns are ordered according to decreasing Spearman correlation to the pseudotime score in Fig. 3D. (C) Representative Palantir pseudotime trajectories generated using alternative Wap-expressing cell-of-origin root cells. Representative trajectories are shown projected onto the epithelial UMAP embedding, with cells colored according to inferred pseudotime and the selected root cell highlighted in yellow. Spearman correlation coefficients between each representative trajectory and the reference trajectory are indicated above each panel.

**Supplementary Method Figure 3 related to Figure 6.**
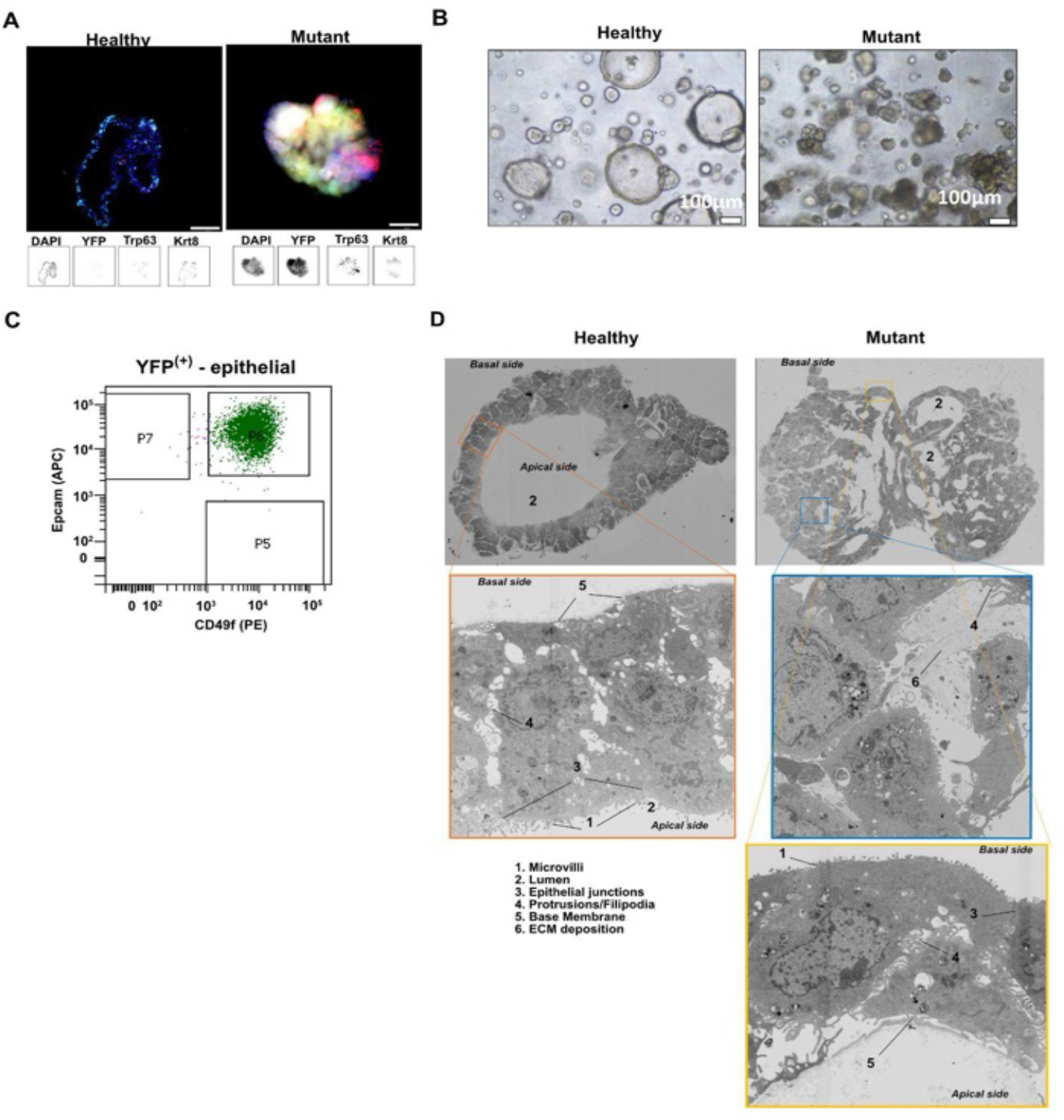
(A) Tumor organoids display a basal phenotype. IF staining of organoid lines from healthy (left) and mutant (right) mammary glands. Channels: DAPI, YFP, Trp63, and Krt8.Scale bars: 100 µm (B) Tumor organoids display a disorganized architecture with loss of the central lumen. Brightfield images of organoid cultures showing the structural organization of epithelial cells. Healthy organoids form acinar-like structures with central lumens; tumor organoids display disorganized architecture. Scale bars: 100 µm. (C) Epithelial cells used for injection experiment were enriched for LP state. Flow cytometry plot showing the sorting strategy for YFP⁺ epithelial cells using PE-CD49f and APC-EpCAM staining, based on the luminal progenitor gating approach described in Rosenbluth et al., 2020. (D) Tumor organoids display protrusions/filipodia invading the surrounding ECM. Transmission electron microscopy of organoids derived from healthy and tumor-bearing mammary glands. Tumor organoids show disrupted epithelial architecture, loss of apical–basal polarity, and protrusions extending into the extracellular matrix. Scale bars: as indicated.

**Supplementary Method Figure 4 related to Figure 7.**
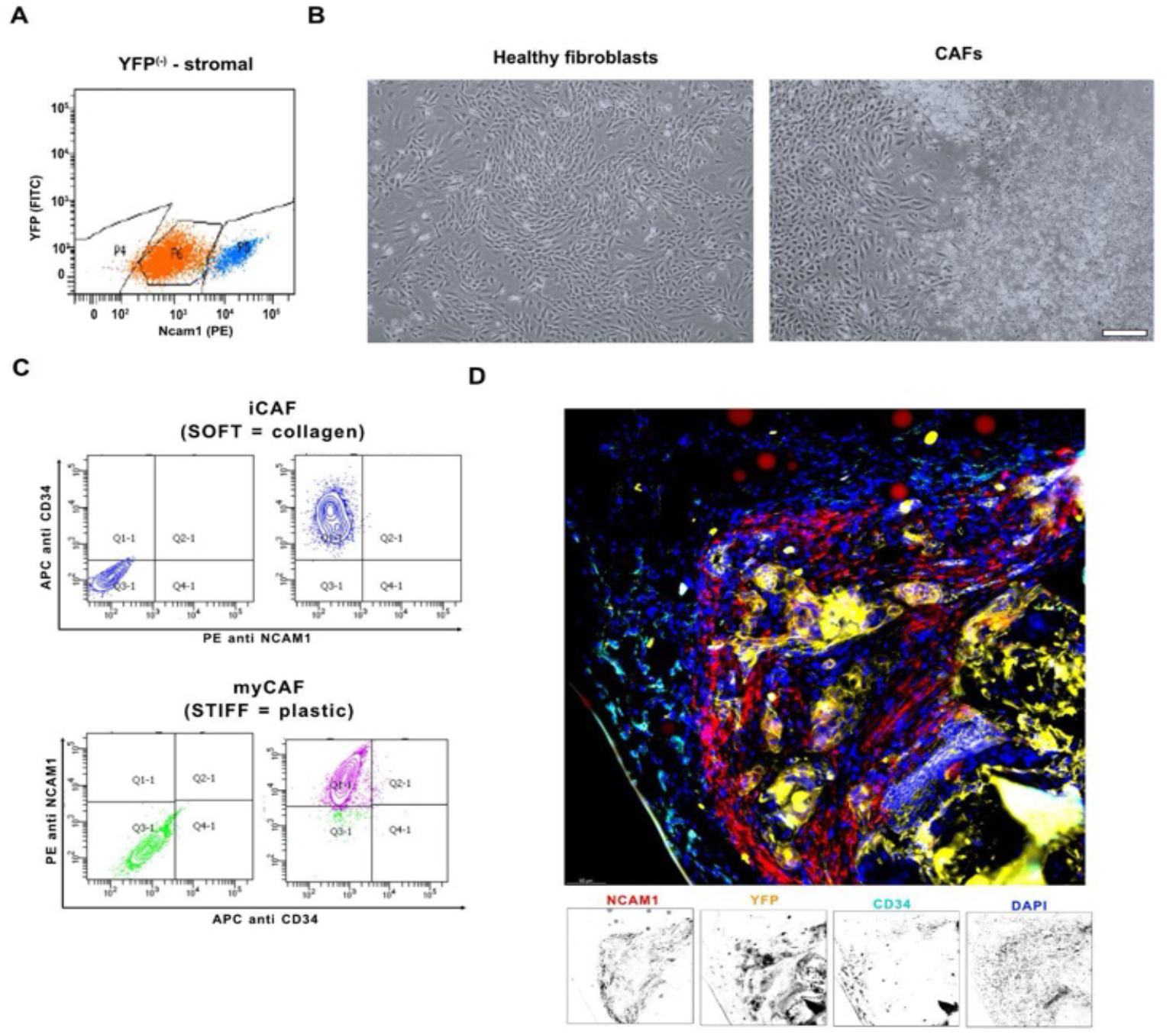
(A) A Tumor fibroblast cultures Ncam1+ and Ncam1- negative populations. Flow cytometry analysis of tumor-derived fibroblasts stained with PE-conjugated anti-Ncam1 antibody. Cells were gated negative for Ptprc and EpCAM. Distinct Ncam1⁺ and Ncam1⁻ populations are shown. (B) Tumor fibroblast cultures display heterogenous morphologies. Brightfield images of fibroblast cultures derived from healthy and tumor-bearing mammary glands showing differences in morphology. Scale bars: 100 µm. (C) Tumor fibroblast cultures strategy for sorting Ncam1+ (myCAF) Cd34+ (iCAF) populations. Flow cytometry analysis of tumor-derived fibroblasts stained with PE- conjugated anti-Ncam1, APC conjugated Cd34 antibody. Cells were gated negative for Ptprc and EpCAM. Distinct Cd34-/Ncam1⁺ (grown on soft matrix formed by collagen (9ug/ul) coating of plastic) and Cd34+/Ncam1⁻ (grown on plastic) populations are shown. (D) iCAF and myCAFs validated in T2 tumor tissue. IF staining of a T2 tumor. Region displays distinct localization of Cd34 (iCAF), Ncam1 (myCAF) around the tumor cells (YFP). Scale bar: 100 µm.

